# S-nitrosoglutathione reductase deficiency causes aberrant placental S-nitrosylation and preeclampsia

**DOI:** 10.1101/2020.07.01.183012

**Authors:** Shathiyah Kulandavelu, Raul A Dulce, Christopher I Murray, Michael A Bellio, Julia Fritsch, Rosemeire Kanashiro-Takeuchi, Himanshu Arora, Ellena Paulino, Daniel Soetkamp, Wayne Balkan, Jenny E Van Eyk, Joshua M Hare

## Abstract

Preeclampsia (PE), a leading cause of maternal and fetal mortality and morbidity, is characterized by an increase in S-nitrosylated (SNO) proteins and reactive oxygen species (ROS), suggesting a pathophysiologic role for dysregulation in nitrosylation and nitrosative stress. Here we show that mice lacking S-nitrosoglutathione reductase (*GSNOR^−/−^*), a denitrosylase regulating protein S-nitrosylation, exhibit a PE phenotype, including hypertension, proteinuria, renal pathology, cardiac concentric hypertrophy, decreased placental vascularization, and fetal growth retardation. ROS, nitric oxide (NO) and peroxynitrite levels are elevated. Importantly, mass spectrometry reveals elevated placental SNO-amino acid residues in *GSNOR^−/−^* mice. Ascorbate reverses the phenotype except for fetal weight, reduces the difference in the S-nitrosoproteome, and identifies a unique set of SNO-proteins in *GSNOR^−/−^* mice. Therefore, deficiency of GSNOR creates dysregulation of placental S-nitrosylation and preeclampsia in mice, which can be rescued by ascorbate. These findings offer valuable insights and therapeutic implications for PE.

Graphical Abstract:
Dysregulation in nitrosylation contributes to nitroso-redox imbalance and nitrosative stress contributing to clinical features of PE including hypertension, proteinuria, concentric hypertrophy in the heart, decrease placental vascularization and fetal growth restriction. Antioxidant treatment rescued the PE-like phenotype in the mother.

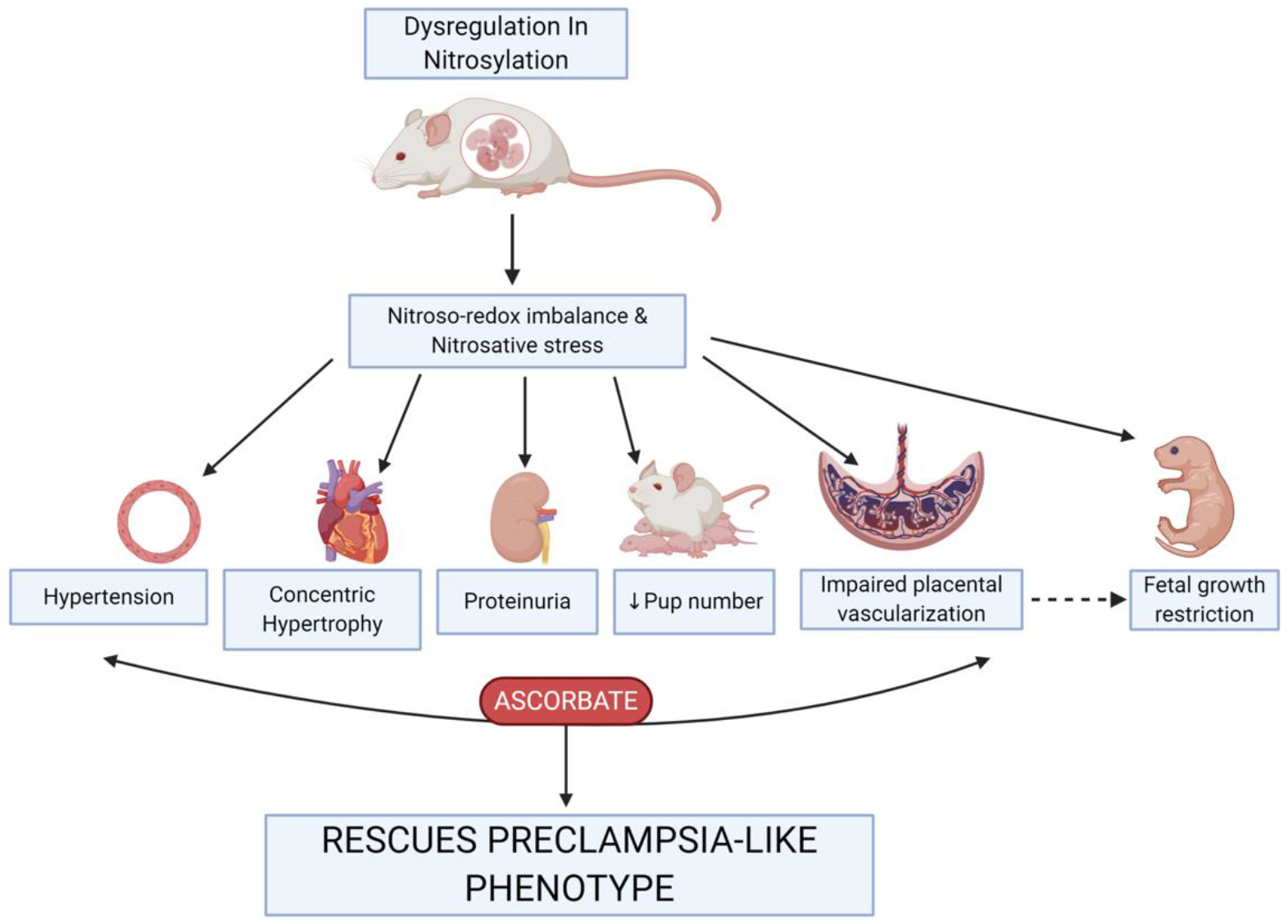

## INTRODUCTION

Preeclampsia (PE), is a life-threatening disorder of pregnancy, characterized by new onset hypertension, proteinuria, abnormal maternal cardiovascular and renal adaptations, poor placental vascularization and fetal growth restriction. PE affects up to 10% of pregnancies, is a leading cause of maternal and fetal/neonatal mortality and morbidity worldwide^1, 2^, and maternal mortality rates have been steadily rising over the past 30 years in large part due to cardiovascular complications^3, 4^. The pathogenesis of PE is incompletely understood, but emerging data support a role for impaired protein S-nitrosylation, nitration, increased ROS, contributing to alterations in NO bioavailability and nitroso-redox imbalance^5–8^. A paradoxical finding in preeclampsia is the elevation in circulating S-nitrosylated (SNO)-albumin^6, 7^ because elevated SNO-albumin would traditionally be considered a vasorelaxant and an anti-oxidant^9^. Furthermore, cellular SNO level is elevated despite factors that are assumed to abrogate the formation of nitroso-thiols, including oxidative stress and defects in nitric oxide (NO) bioavailability^7, 10^. These findings raise two alternative pathogenic possibilities: Either SNO increases to compensate for the increased ROS levels or elevated SNO directly reflects abnormal regulation of SNO in preeclampsia (nitrosative stress). To differentiate between these two possible roles, we studied the impact of the loss of an important denitrosylase, S-nitrosogluathione reductase (*GSNOR^−/−^*) in pregnant mice.

Protein S-nitrosylation participates in numerous pregnancy-related processes including placental trophoblast cell migration, apoptosis, angiogenesis, immunomodulation, and oxygen delivery^11, 12^. Protein SNO is enhanced in a transnitrosylation reaction using S-nitrosogluathione (GSNO), which thus acts as a second messenger to transduce NO bioactivity^13^. This process is tightly regulated by GSNOR, which selectively metabolizes GSNO thereby depleting the levels of S-nitrosylated proteins in equilibrium with GSNO. Ascorbate, with its dual roles as an antioxidant, and a reductant is required for the release of biologically active NO from these nitrosylated thiols. Although homozygous deletion *GSNOR^−/−^* mice have increased numbers of S-nitrosylated proteins^14, 15^, which can in some circumstances lead to favorable outcomes (e.g. recovery from myocardial infarction)^16^, deficiency of this enzyme can also disrupt physiological nitrosylation-denitrosylation dynamic cycles leading to unfavorable outcomes (e.g. increase in oxidative stress)^15, 17, 18^. Thus, to assess the role of S-nitrosylation and GSNOR in pregnancy outcome, we examined multiple organ systems, including the heart, kidney, placenta, and the offspring during pregnancy in *GSNOR^−/−^* mice. Initially, we anticipated that *GSNOR^−/−^* mice would exhibit favorable maternal and fetal adaptations to pregnancy and were surprised to discover that pregnant *GSNOR^−/−^* mice recapitulate many of the features of preeclampsia. As a result, we subsequently tested the predictions that placental nitrosylation would exhibit widespread derangement in the knockout compared with control mice, and that ascorbate would restore the intact animal and nitroso-proteome phenotypes towards normal.

## RESULTS

### Pregnant *GSNOR^−/−^* mice exhibit hallmark features of PE, hypertension and proteinuria

Development of hypertension and proteinuria are hallmarks of PE, and *GSNOR^−/−^* mice are hypotensive compared to wild type mice prior to pregnancy^14^, consistent with GSNOR regulation of endothelium-dependent vasodilation (Figure 1A). At day 17.5 of pregnancy, *GSNOR^−/−^* mice develop hypertension (Figure 1A) and proteinuria (Figure 1B), characterized by elevated urine macroglobulin levels, associated with renal pathology including enlarged glomeruli, swelling of the endothelial cells and loss of fenestration and corrugation of the glomerular basement membrane (Figures 1C-G); changes that recapitulate human preeclampsia^19, 20^.

**Figure 1:**
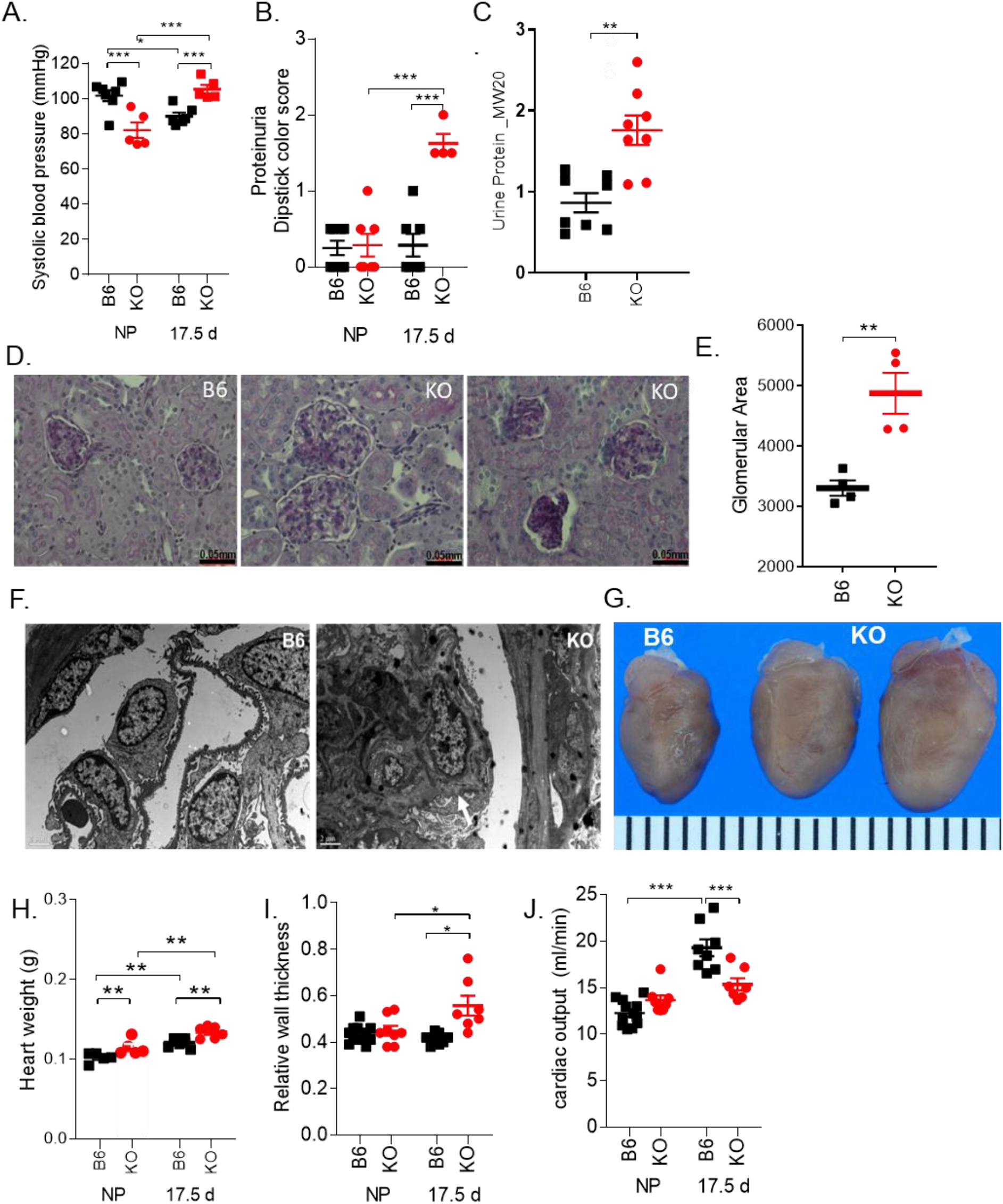
Pregnant *GSNOR^−/−^* mice exhibit hallmark features of PE including hypertension, proteinuria, and concentric hypertrophy in the heart. Non-pregnant and pregnant (17.5 d) (N=4-12 mothers per group) C57Bl/6J (B6) and *GSNOR^−/−^* (KO) mice were examined. At late gestation, KO mothers exhibited (**A**) hypertension, (**B**) proteinuria and elevated (**C**) urine macroglobulin levels. (**D, E**) Kidney sections at late gestation stained with periodic acid-Schiff showed enlarged glomeruli and focal and sclerosis with collapsed glomerular capillaries. (**F**) Electron microscopy on renal tissues showed that GSNOR^−/−^kidneys exhibited glomerular endotheliosis comprised of endothelial cell swelling, along with loss in fenestration and corrugation of the glomerular basement membrane (N=2 was examined per group). (**G, H**) Heart weight and (**I**) relative wall thickness were significantly bigger in KO mice as compared to controls, indicating the presence of concentric hypertrophy. (**J**) The normal increase in cardiac output was absent in KO mice at late gestation. Results are shown as mean ± SEM. *** P<0.001, **P<0.01, *P<0.05. NP, non-pregnant; B6, C57Bl/6J mice; KO, *GSNOR^−/−^* mice. For systolic blood pressure, proteinuria, heart weight, relative wall thickness and cardiac output, a 2-way ANOVA with Newman-Keuls for post hoc analysis was performed. For the other variables, Student’s T-test was performed.

### *GSNOR^−/−^* hearts exhibit concentric hypertrophy during pregnancy

We next studied cardiac responses to pregnancy in the *GSNOR^−/−^* mouse. The heart responds to sustained hypertension by an increase in wall thickness leading to concentric hypertrophy, and cconcentric hypertrophy independently predicts adverse outcome in preeclamptic pregnancies^21^. At late gestation, left ventricular end-diastolic dimension was lower, whereas the anterior wall at diastole (Table 1) was thicker, contributing to higher relative wall thicknesses in pregnant *GSNOR^−/−^* mice as compared to controls (Figures 1H,I, Table 1). We next isolated cardiomyocytes (CMs) and observed that CM width at late gestation was greater in *GSNOR^−/−^* mice, whereas CM length was not different between the two strains (Table 1). These factors likely contributed to the enlarged *GSNOR^−/−^* hearts at late gestation even when normalized to tibia length (Figure 1H, Table 1). Furthermore, the normal physiologic increases in maternal cardiac output and stroke volume were completely abrogated at late gestation in *GSNOR^−/−^* mice (Figure 1J, Table 1), consistent with the phenotype of lower cardiac output and stroke volume in preeclamptic patients with concentric heart geometry^21^. This data suggests that SNO homeostasis plays an important role in limiting pathological hypertrophy^22^.

**Table:** KO mice exhibited concentric hypertrophy and abnormal maternal cardiovascular adaptation to pregnancy, which was rescued with ascorbate treatment. Results are shown as mean ± SEM. *P<0.05 vs. pregnancy (same strain); # vs. strain difference. Asc, ascorbate. LV, left ventricle. Two-way ANOVA with Newman-Keuls for post hoc analysis. * P<0.05 vs pregnancy (same strain); # vs strain difference.

|  | Non-pregnant |  | Pregnant |  | Asc-treated Pregnant |  |
| --- | --- | --- | --- | --- | --- | --- |
|  | B6 | KO | B6 | KO | B6 | KO |
| Maternal body weight (g) | 20.3 ± 0.36* | 20.3 ± 0.70* | 35.4 ± 0.56 | 31.1 ± 1.55<sup>#</sup> | 36.24 ± 0.78 | 34.2 ± 0.55* |
| LV end-diastolic dimension, diastole (mm) | 3.40 ± 0.04* | 3.50 ± 0.05 | 3.78 ± 0.04 | 3.52 ± 0.12<sup>#</sup> | 3.67 ± 0.07 | 4.00 ± 0.09*<sup>#</sup> |
| LV anterior wall thickness, diastole (mm) | 0.80 ± 0.02 | 0.82 ± 0.04* | 0.91 ± 0.03 | 1.07 ± 0.07<sup>#</sup> | 0.84 ± 0.03 | 0.88 ± 0.04* |
| Relative wall thickness | 0.43 ± 0.01 | 0.45 ± 0.02* | 0.41 ± 0.01 | 0.56 ± 0.04<sup>#</sup> | 0.44 ± 0.01 | 0.45 ± 0.03* |
| Myocyte length (μM) | 138 ± 2* | 164 ± 4*<sup>#</sup> | 158 ± 2 | 154 ± 2 | 158 ± 7 | 150 ± 2 |
| Myocyte width (μM) | 25.7 ± 0.28 | 25.1 ± 0.46* | 25.2 ± 0.53 | 29.4 ± 0.67<sup>#</sup> | 25.3 ± 0.34 | 22.4 ± 0.29*<sup>#</sup> |
| Stroke volume (ul) | 27.1 ± 0.50* | 27.8 ± 1.16 | 39.2 ± 1.96 | 30.9 ± 1.27<sup>#</sup> | 34.5 ± 1.54* | 40.1 ± 1.60*<sup>#</sup> |
| Heart weight normalized to tibial length (mg) | 5.88 ± 0.13* | 7.09 ± 0.31*<sup>#</sup> | 7.07 ± 0.10 | 7.98 ± 0.34<sup>#</sup> | 7.01 ± 0.63 | 9.98 ± 3.38<sup>#</sup> |

### *GSNOR^−/−^* mice exhibited placental insufficiency during pregnancy

Placental insufficiency (fetal weight to placental weight ratio) is believed to contribute directly to the development of preeclampsia. The placentas of preeclamptic pregnancies often exhibit fetoplacental hypovascularity and decreased fetoplacental perfusion^23^, both of which decrease transfer of oxygen and nutrients to the placenta, thereby limiting fetal growth. Fetal litter size was significantly lower in *GSNOR^−/−^* mice at 17.5 d of gestation (Figure 2A). Fetal body weights were significantly lower in *GSNOR^−/−^* mice at 17.5 d of gestation (Figure 2B). To evaluate the role of GSNOR on placental vascularization and fetoplacental perfusion, we examined the placentas at late gestation. GSNOR activity was present in the control placentas, and as anticipated was absent in the knockout mice (Figure 2C). Placental weight (Figure 2D) and umbilical arterial blood flow (Figure S1) were not significantly different between the two strains. However, placental efficiency and umbilical venous blood flow were significantly lower in *GSNOR^−/−^* placentas as compared to controls at late gestation (Figures 2E, S1). In addition, placental vascularization was decreased in *GSNOR^−/−^* placentas (Figures 2F,G). These findings suggest that GSNOR plays an essential role in placental development and function during pregnancy.

**Figure 2:**
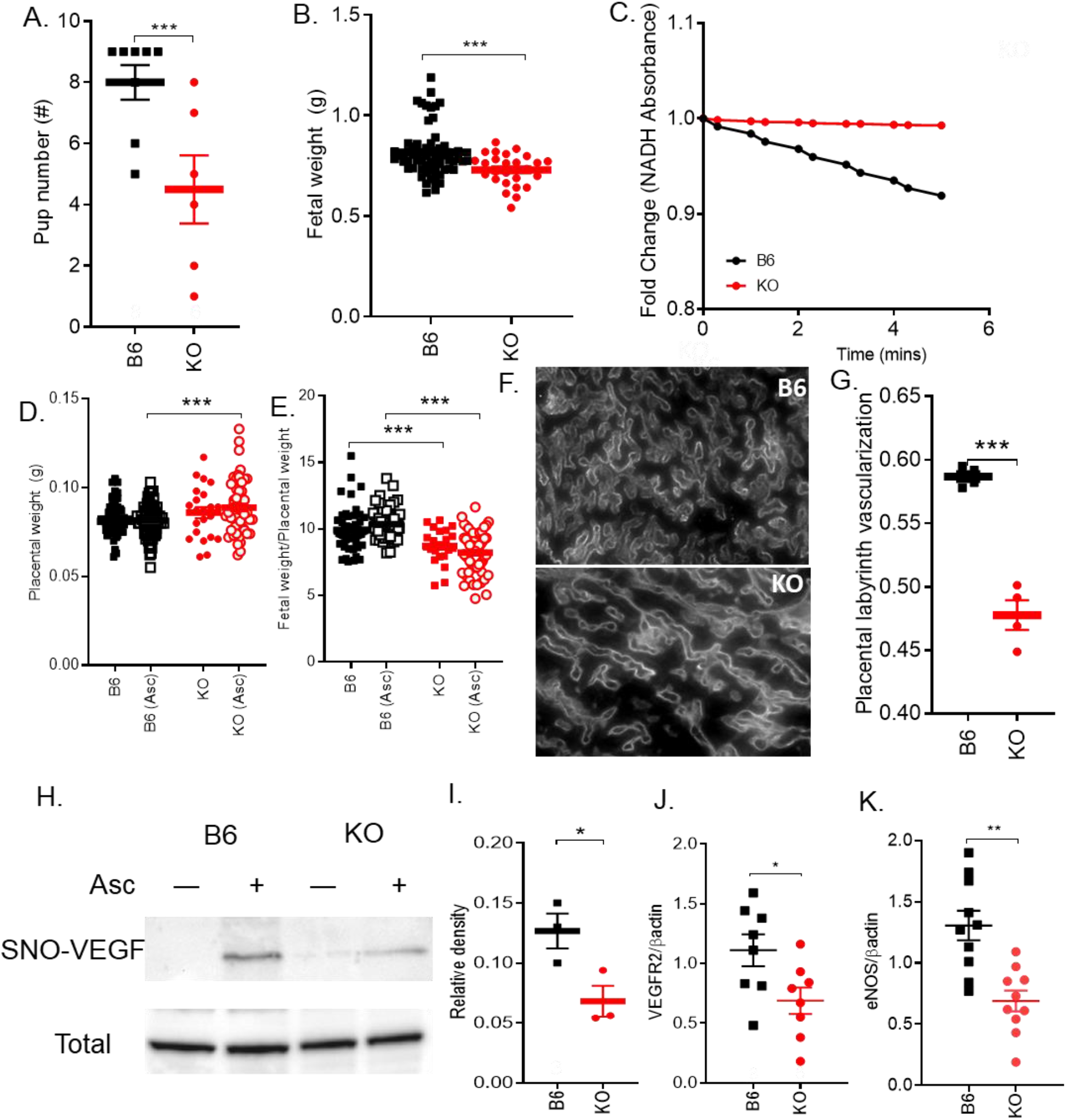
*GSNOR^−/−^* mice exhibited placental insufficiency during pregnancy and showed alteration of VEGF pathway. Non-pregnant and pregnant (17.5 d) (N=4-12 mothers per group) C57Bl/6J (B6) and *GSNOR^−/−^* (KO) mice were examined. (**A**) Fetal number and (**B**) weight were significantly lower in KO mice. (**C**) NADH-dependent GSNOR enzymatic activity was determined in placental tissue at 17.5 days of gestation. GSNOR activity is enriched in B6 placentas, whereas it is completely absent in the KO placentas. (**D, E**) Placental weight was not significantly different between the two strains, whereas placental efficiency was significantly lower in KO mice as compared to control. (**F, G**) Placental vascularization determined using isolectin immunostaining, was significantly lower in KO as compared to B6 at late gestation. SNO-VEGF was measured using Biotin-switch assay. (**H**) Representative blots shown for S-nitrosylated and total VEGF. Omission of ascorbate was used as the negative control. (**I**) Nitrosylation of VEGF was significantly lower in KO placentas as compared to B6 at late gestation. (**J, K**) VEGFR2 and eNOS protein levels were determined in the placentas at 17.d of gestation. Both VEGFR2 and eNOS protein levels were significantly lower in in GSNOR^−/−^ placentas as compared to B6 placentas at 17.5 d of gestation. Results are shown as mean ± SEM. *** P<0.001, **P<0.01, *P<0.05. NP, non-pregnant; B6, C57Bl/6J mice; KO, *GSNOR^−/−^* mice. For statistical significance, 2-way ANOVA with Newman-Keuls for post hoc analysis or Student’s T-test were performed.

### VEGF pathway was blunted in *GSNOR^−/−^* placentas at late gestation

We next examine the vascular endothelial growth factor (VEGF) pathway as a potential mechanism for the impaired placental vascularization as VEGFR2 levels have been shown to be decreased in human preeclamptic pregnancies^24^. Whereas, VEGF protein abundance in the placentas was not different between the two strains at late gestation (Figure 2H), nitrosylation of VEGF protein was significantly lower in *GSNOR^−/−^* placentas (Figures 2H,I). VEGF binds to its receptor VEGFR2 and signals through eNOS pathways. Like human PE, we found that VEGFR2 (P<0.05) and eNOS (P<0.01) total protein quantities were lower in the *GSNOR^−/−^* placentas as compared to controls (Figures 2J,K). Thus, alterations in the VEGF signaling may represent a contributory mechanism for impaired placental development in GSNOR^−/−^ mice.

### *GSNOR^−/−^* mice exhibit nitroso-redox imbalance and nitrosative stress during pregnancy

Aberrant ROS and nitric oxide signaling has been implicated in the pathogenesis of PE ^25, 26^. To address this issue, we measured the cellular ROS levels in *GSNOR^−/−^* mice. Prior to pregnancy, cellular ROS generation was higher in cardiomyocyte (CMs) isolated from *GSNOR^−/−^* mice as compared to control mice (Figure 3A), yet ROS levels were not significantly different between the two groups (Figure 3B), suggesting increased involvement of antioxidant scavengers. As in the case with humans, pregnancy^27^ itself increases ROS generation as seen in hearts obtained from control pregnant animals as compared to those from non-pregnant animals (Figures 3A,B). Physiologic oxidative stress is proposed to be necessary for a wide array of physiological functions, including antioxidant defense and continuous placental remodeling^27^. While in pregnancy, ROS generation and levels were significantly higher in *GSNOR^−/−^* CM as compared to controls (Figures 3A,B) confirming the presence of oxidative stress. Increased oxidative stress can lead to alterations in NO/SNO signaling, so we next measured NO levels. As expected, there was a significant increase in NO levels in isolated CMs of control mice during pregnancy (Figure 3C). This increase likely plays a critical role in vasodilation leading to the normal adaptations to pregnancy. Furthermore, NO levels were significantly higher in CMs isolated from pregnant as compared to non-pregnant *GSNOR^−/−^* mice. With the presence of elevated ROS and NO/SNO levels in *GSNOR^−/−^* mice, we predicted an increase production of the potent pro-oxidant, peroxynitrite. Prior to pregnancy, peroxynitrite levels were significantly higher in *GSNOR^−/−^* CMs as compared to controls (Figure 3D). This preexisting elevation in peroxynitrite levels suggests that the *GSNOR^−/−^* mice were less able to respond to the stress of pregnancy. At late gestation, peroxynitrite levels were, indeed, significantly higher in isolated CMs of *GSNOR^−/−^* mice (Figure 3D), suggesting the presence of nitrosative stress. Furthermore, superoxide dismutase (SOD) levels were significantly lower in *GSNOR^−/−^* placentas as compared to controls suggesting decrease in antioxidant capacities (Figure 3E). NO signals via SNO-proteins may exert a protective role against an oxidative environment by competing with other post-translational modifications and shielding critical cysteine residues from the damaging effects of irreversible oxidation as shown in ischemic preconditioning^28, 29^. These findings suggests that SNO-based mechanisms that protect physiologic signaling may be impaired in pregnant *GSNOR^−/−^* mice contributing to the PE phenotype. Alternatively, increase levels of NO, ROS can form peroxynitrite, which can irreversibly lead to protein nitration^8^. Protein nitration plays a relevant role in post-translational modification of protein and increased nitration of proteins have been implicated in human PE pregnancies^8^. Thus, protein nitration may be another mechanism involved in the development of PE-like conditions in *GSNOR^−/−^* animals.

**Figure 3:**
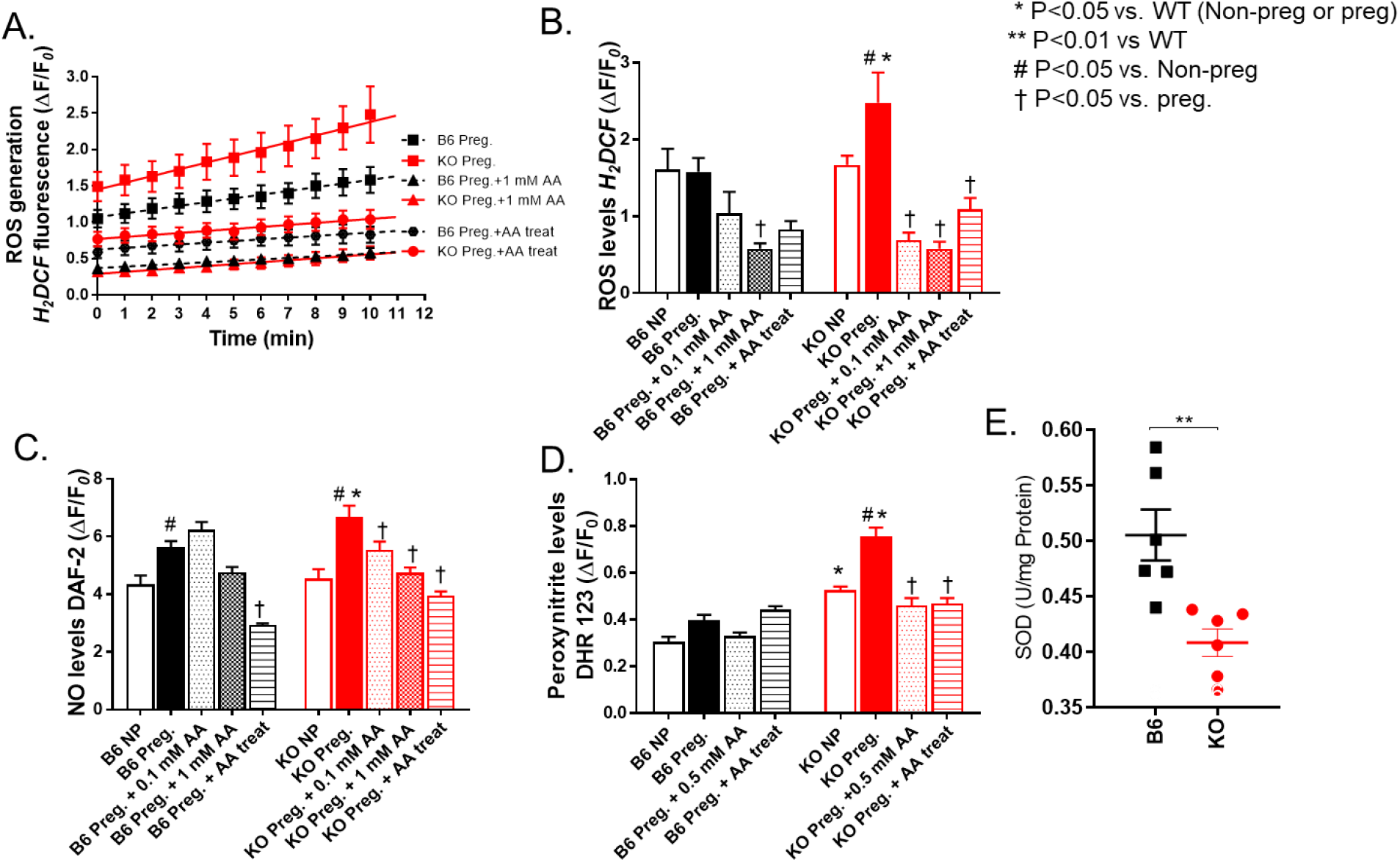
*GSNOR^−/−^* mice exhibit nitroso-redox imbalance and nitrosative stress that is rescued with ascorbate treatment. In non-pregnant (NP) and pregnant (17.5 d) C57Bl/6J (B6) and *GSNOR^−/−^* (KO) mice, reactive oxygen species (ROS), nitric oxide (NO) and peroxynitrite levels were determined in isolated cardiomyocyte using epifluorescence by 2’,7’dichlorodihydro-fluoresceine (H2DCF‒DA), 4,5-diaminofluorescein (DAF-2DA) and dihydrorhodamine 123 (DHR 123), respectively. Ascorbate (AA) studies included both acute (0.1 mM, 0.5 mM, 1 mM) and chronic (AA provided in drinking water from day 0.5) treatments. (**A-B**) ROS generation and levels, and (**C**) NO and (**D**) peroxynitrite levels were significantly higher in CM isolated from pregnant *GSNOR^−/−^* mice. This increase was prevented with acute and chronic AA treatment. (**E**) Superoxide dismutase (SOD) levels were measured in the placenta using lucingenin-enhanced chemiluminescence. SOD levels were significantly lower in the KO placentas as compared to controls. Results are shown as mean ± SEM. N=3-5 animals per experiment. NP, non-pregnant; B6, C57Bl/6J mice; KO, *GSNOR^−/−^* mice; AA, Ascorbate.1-way ANOVA with Newman Keuls post-hoc test.

### Antioxidant treatment rescued the PE phenotype in *GSNOR^−/−^* mice during pregnancy

To test whether antioxidant treatment can rescue the PE phenotype, we treated the animals or isolated CMs with ascorbate. In addition to being an antioxidant, ascorbate also functions as a reductant, promoting the release of biologically active NO from nitrosylated thiols, and plasma ascorbate levels are commonly diminished in preeclamptic patients^30^. Importantly, ascorbate treatment rescued the onset of hypertension, proteinuria, and urinary macroglobulin levels (Figures 4A-C). The enlarged anterior wall thickness, relative wall thickness and CM width, all indicative of concentric hypertrophy, returned to normal levels in ascorbate treated pregnant *GSNOR^−/−^* mice (Table 1). Ascorbate also significantly improved cardiac output and stroke volume in *GSNOR^−/−^* mothers at late gestation, whereas it lowered both parameters in pregnant B6 controls (Figure 4D, Table 1). Ascorbate treatment significantly improved placental vascularization along with placental VEGFR2 and eNOS protein levels (Figures 4G,J-K). In addition, ascorbate improved umbilical venous blood flow and litter size, but did not improve placental efficiency, which may account for the failure of fetal weights to improve at late gestation in the *GSNOR^−/−^* mice (Figures. 4E-I, S1). Acute and chronic treatment of ascorbate similarly reduced ROS, NO and peroxynitrite levels in isolated CMs from pregnant *GSNOR^−/−^* mice (Figure 3A-D). Thus, ascorbate may work as a potent scavenger of free radicals as seen in experimental models of hypertension^31, 32^ and in rat models of nephrotoxicity^33^ to balance the nitroso-redox system and in turn rescue the PE-like phenotype in pregnant *GSNOR^−/−^* mice.

**Figure 4:**
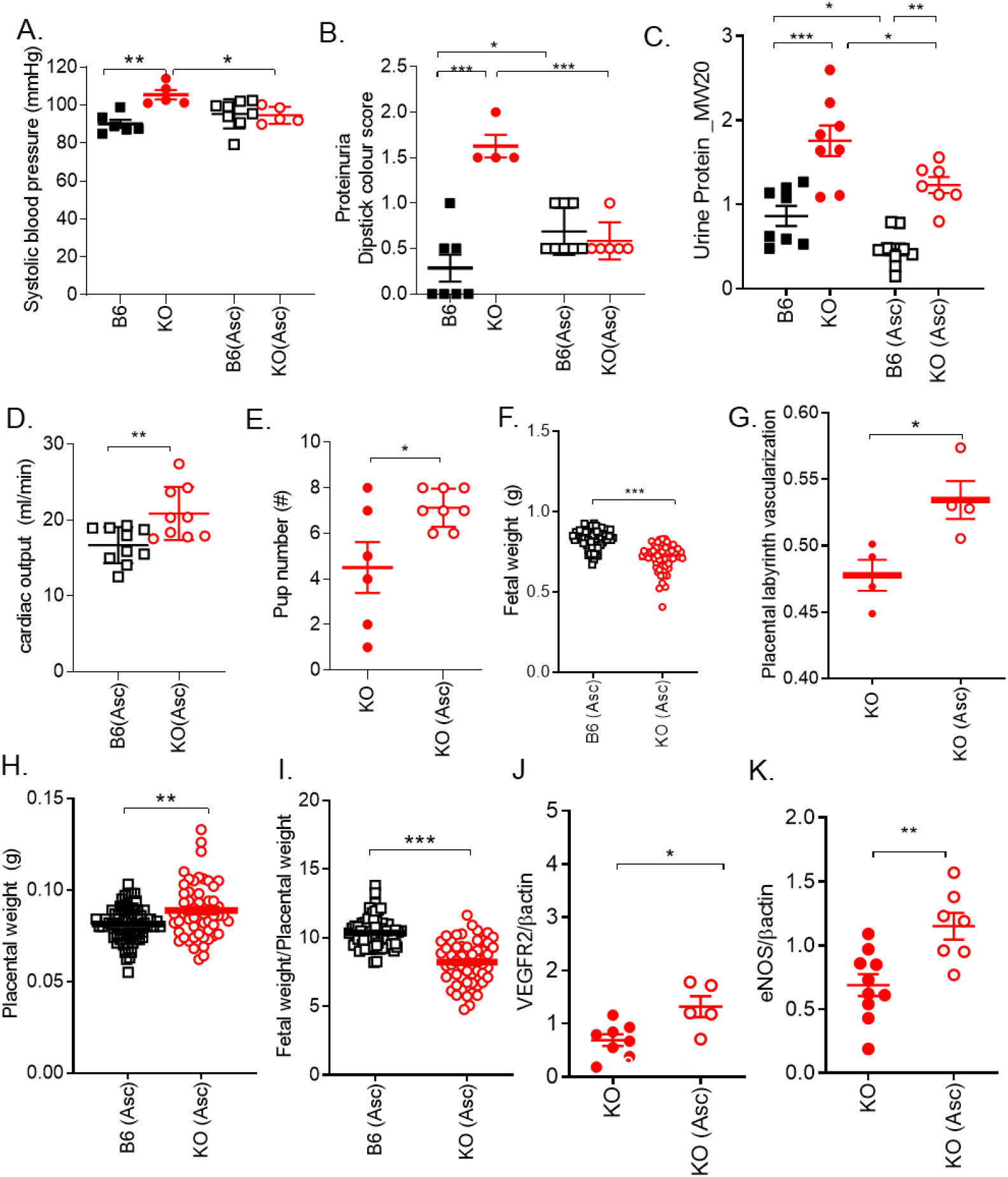
PE phenotype in the mother and litter size is rescued with ascorbate. Late pregnant (17.5 d) C57Bl/6J (B6) and *GSNOR^−/−^* (KO) mice were examined. N=4-10 mothers per group. (**A**) Hypertension, (**B**) proteinuria and (**C**) urine macroglobulin levels were rescued with ascorbate treatment. Ascorbate treatment increased (**D**) cardiac output in KO mice at late gestation. (**E**) Pup number was improved, whereas (**F**) fetal weight remained significantly lower in KO mice treated with ascorbate. (**G**) Impaired placental vascularization was rescued with ascorbate treatment in KO placentas. With ascorbate treatment, (**H**) placental weight was significantly higher in the KO treated animals as compared to B6 treated animals. Whereas (**I**) placental efficiency remained significantly lower in treated KO mice as compared to controls. (**J, K**) VEGFR2 and eNOS placental protein levels were significantly increased in KO (Asc) treated animals as compared to non-treated KO animals. Results are shown as mean ± SEM. ***P<0.001, **P<0.01, *P<0.05. 1-way or 2-way ANOVA with Newman Keuls post hoc test and Student’s T-test were performed. Asc, ascorbate; B6, C57Bl/6J mice; KO, *GSNOR^−/−^* mice.

### Mass Spectrometry revealed elevated placental SNO-amino acid residues in *GSNOR^−/−^* mice

Next in order to directly test the prediction that SNOylation is dysregulated in *GSNOR^−/−^* placentas, we performed mass spectrometry using both thiol reactive biotin-HPDP and cysTMT labeling to maximize coverage^34^. Consistent with our prediction, this analysis revealed a marked increased number of SNOylated residues in *GSNOR^−/−^* placentas (459 corresponding to 351 proteins) compared to controls (264 SNOylated residues corresponding to 198 proteins) (Figure 5A-C, S2, Table S1-3). Importantly, placentas from ascorbate treated mice exhibited an increased net number of SNOylated proteins in both control and *GSNOR^−/−^* placentas (consistent with the transnitrosylation properties of chronic ascorbate^35^), but this increase was less in the *GSNOR^−/−^* mice (Figure 5D) decreasing the difference in number of SNO-proteins between the two groups. To gain insights into the most important signaling pathways affected by excess S-nitrosylation we examined the subset of SNO-proteins found exclusively in GSNOR versus B6, and which exhibited denitrosylation with ascorbate. Of all the detected peptides, there were 50 SNO-residues unique to the *GSNOR^−/−^* placentas, but only 16 residues unique to the B6 placentas (Figures 5C,E, Table S1-3). All 50 SNO-residues were reversed by ascorbate (Table S2). From these 50 proteins, 14 have been linked to important roles in processes essential in pregnancy, including angiogenesis, inflammation, cell migration and apoptosis (Figure 5E), supporting the pathophysiological relevance of these proteins.

**Figure 5:**
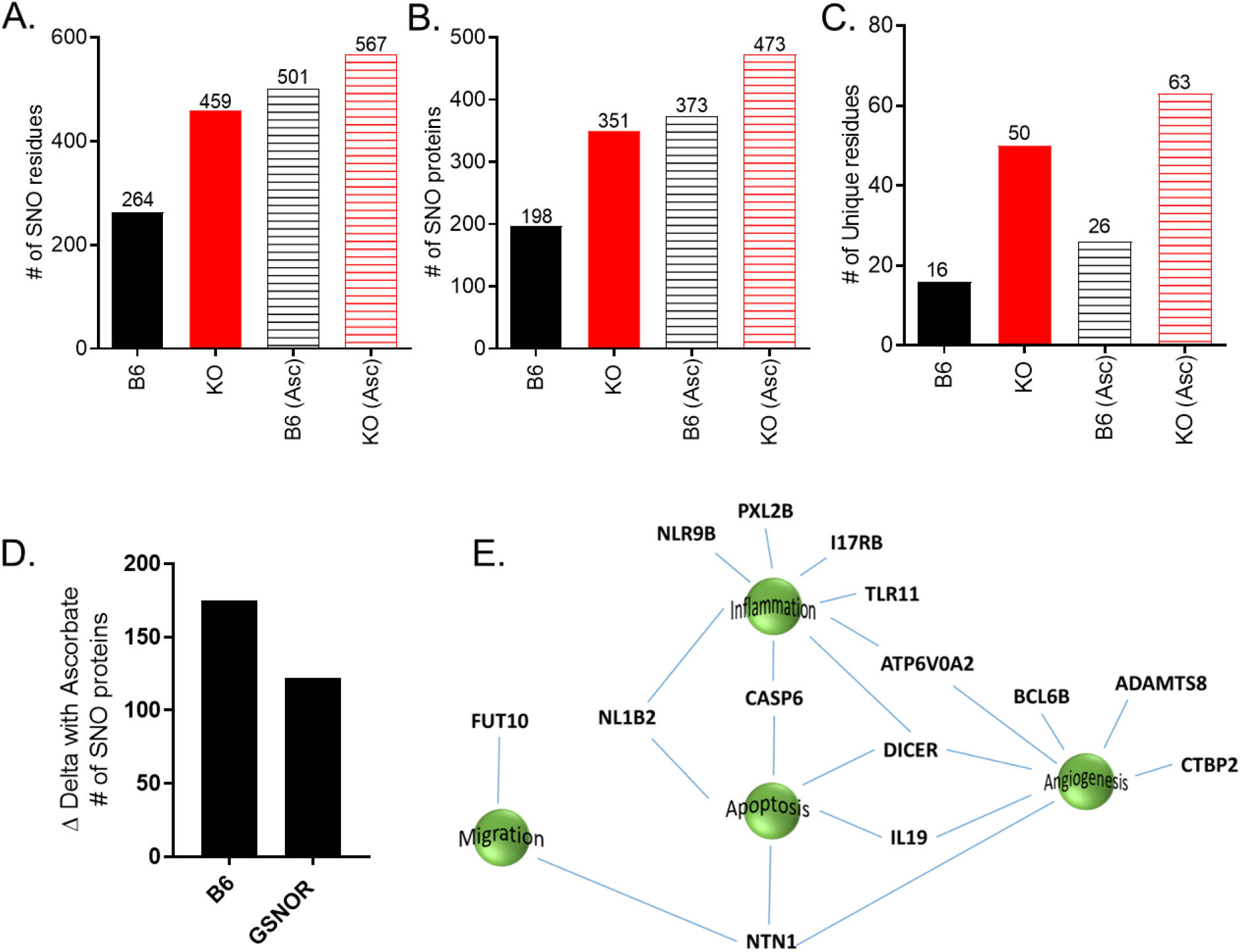
Duel-labeling mass spectrometry revealed an increased number of SNOylated proteins in the placentas from *GSNOR^−/−^* animals. (A, B) An increased number of SNOylated proteins were detected in the placentas of the KO animals, and of these there were more unique SNOylated residues (**C**) in *GSNOR^−/−^* placentas as compared to controls. Ascorbate treatment increased SNO-proteins in both groups, but this increase was less in the KO group as compared to control (D). (**E**) Schematic showing proteins unique in *GSNOR^−/−^* placentas but absent in the B6 placentas, and B6- and *GSNOR^−/−^* placentas treated with ascorbate. CASP6 = Caspase-6 (CASP-6); NLR9B = NACHT, LRR and PYD domains-containing protein 9B; TLR11 = Toll-like receptor 11; DICER = Endoribonuclease Dicer; PXL2B = Prostamide/prostaglandin F; IL17RB = IL-17 receptor B; FUT10 = Alpha-(1,3)-fucosyltransferase 10; NTN1 = Netrin-1; CASP6 = Caspase-6; NL1B2 = NACHT, LRR and PYD domains-containing protein 1b allele 2; IL-19 = Interleukin-19; BCL6B = B-cell CLL/lymphoma 6 member B protein; CTBP2 = C-terminal-binding protein 2; ADAMTS8 = A disintegrin and metalloproteinase with thrombospondin motifs 8; ATP6V0A2 = V-type proton ATPase 116 kDa subunit a isoform 2.

## DISCUSSION

We have identified that mice lacking S-nitrosoglutathione reductase (GSNOR−/−), a denitrosylase regulating S-nitrosylation, exhibit most of the clinical features of PE including hypertension, proteinuria, renal pathology, cardiac concentric hypertrophy, decreased placental vascularization and fetal growth restriction. The primary mechanisms involved in this PE phenotype appears to be nitrosative stress due to aberrant S-nitrosylation leading to the presence of nitroso-redox imbalance. In addition, we showed that antioxidant, ascorbate, rescued the nitrosative stress and the PE phenotype in the mother (Graphical Abstract). Our findings demonstrate that the absence of a single gene, GSNOR, alters large numbers of downstream signaling pathway. Ascorbate, which creates a net increase in the number of SNO-proteins, decreases the differences between the GSNOR^−/−^ and B6 mice. As such these results suggest that the regulation of the SNO-integrated post-translational modification system may account in large part for the phenotype of PE in the *GSNOR^−/−^* mice. Together, these results suggest that this system-wide alteration in SNO-proteins has a detrimental effect on the function of these pathways and as such, could be a key mechanism involved in the pathological phenotype seen in multiple organ systems, including the heart, kidney, placenta, and the offspring of the KO animals with PE.

One of the primary clinical features of PE is an elevation in blood pressure. Number of different mechanisms may be involved in alterations in blood pressure in the *GSNOR^−/−^* animals. Elevated plasma SNO can lead to adverse cardiovascular outcomes in patients with end-stage renal disease, which correlates with elevated blood pressure^36^. Gandley et al.^7^ postulated that the buffering function of SNO-albumin was impaired in preeclamptic patients, where the thiol of albumin acts as a sink for NO, therefore lowering NO bioavailability and thus raising blood pressure. Alternatively, denitrosylation of SNO-albumin may be regulated by glutathione (GSH). In preeclampsia, plasma GSH levels are low^37^, most likely due to oxidative stress, and this decrease may effect NO-dependent vasodilation of red blood cells, as GSH may facilitate their export of SNOs^38^. In addition, ROS which is increased in preeclampsia, potentiates protein S-nitrosylation^39^. Therefore, an altered redox state, which influences the thiol/nitrosothiol balance, may convey NO bioactivity, regulating free NO, thereby affecting blood pressure in *GSNOR^−/−^* mice.

Several clinical trials have examined the effectiveness of antioxidants (ascorbate) in preventing PE. Small, randomized placebo-controlled trials showed reduced PE with ascorbate treatment, whereas all large multicenter randomized trials have yielded disappointing results. These large trials showed that ascorbate did not decrease the risk of PE in either high risk (women with type-1 diabetes, nutritionally deficient, poor social economic status) or low risk populations ^40–42^. In turn, increased incidence of low birthweight, gestational hypertension, fetal loss, stillbirth, or premature rupture of membranes have been reported in some trials, but these findings were not confirmed across all studies; therefore their significance remains uncertain^40–42^. Dosage used and/or timing of the antioxidant treatment (8-22 weeks) may have played a role in the unsuccessful outcomes in these large clinical trials. Alternatively, oxidative stress may be relevant to the pathogenesis in only a subgroup of women, with no appreciable benefit of antioxidant therapy for the overall population. Furthermore, preeclampsia is a multi-systemic/multi-factorial disease and based on the heterogeneity of the clinical presentation, there may be different “subtypes” of preeclampsia^43^. This observation may explain, in part, why many of the clinical trials using ascorbate have yet to show favorable outcomes. Therefore, identifying women showing dysregulation in nitrosylation and/or altered GSNOR activity levels may be the ideal target sub-population for treatment with ascorbate, permitting a precision medicine approach for future clinical trials.

This study has several limitations. All the features of the murine phenotype may not translate into human pathophysiology given its heterogenous nature and involvement of multiple organ systems. Furthermore, *GSNOR^−/−^* mice do show pathological phenotypes of PE in the mother’s heart, kidney, placenta and in the offspring, we did not examine some other clinical features of PE such as thrombocytopenia, liver enzyme or spiral artery remodeling in the placenta.

Development of effective therapies for PE is hampered by a failure to understand the causative mechanisms involved and the absence of robust animal models that exhibit all the clinical features of PE. Therefore, the identification of the GSNOR^−/−^ mice as a model that exhibits essentially all the clinical features of preeclampsia, with the causative role of dysregulation in nitrosylation contributing to nitrosative stress and dysregulation of physiologic post-translational modification in a large number of signaling pathways as one of the primary mechanisms contributing to this disorder, has important implications for developing novel therapies and identifying clinically useful biomarkers for this difficult to treat maternal-fetal syndrome.

## MATERIALS AND METHODS

### Breeding

C57Bl/6J (wild type [B6]) controls (stock No. 000664) were purchased from Jackson Laboratories (Bar Harbour, Maine). GSNOR^−/−^ mice were raised in house. GSNOR^−/−^ mouse line was created from ES clones after ten consecutive backcrosses with C57Bl/6J^14^. Females were bred at 3-4 months of age and were studied in their first pregnancies. The presence of a sperm plug was defined as day 0.5 of gestation. Experimental time points included prior to breeding (non-pregnant), and day 17.5 (late gestation, 2 days prior to normal term delivery). Fetal, placental, and maternal organ weights were also recorded at time of sacrifice. For breeding, B6 females were bred with B6 males and GSNOR^−/−^ females were bred with GSNOR^−/−^ males.

### Blood pressure determination

Mice were anesthetized with isoflurane. A 1.4-F micromanometer-tipped catheter (SPR-839; Millar Instruments, City, TX) was inserted into the right carotid artery and advanced retrograde into the aorta. All analyses were performed using LabChart Pro 7 software (Millar Instruments).

### Urinary protein measurements

Urine samples were collected from non-pregnant and pregnant mice (17.5 d of gestation). Urinary protein levels were measured using dipstick. A score of 0 to 3 was given based on the color change on the dipstick following urine analysis. 24-hour urine was collected using metabolic cages (Tecniplast). Urine samples were analyzed for macroglobulin levels using Coomassie blue staining (Fisher).

### Echocardiography

Echocardiographic assessments were performed in anesthetized mice (1% isoflurane in oxygen) using a Vevo-770 micro-ultrasound (VisualSonics, Toronto, Ontario) equipped with a 30-MHz transducer. Cardiac dimensions including LV end-diastolic diameter, LV end-systolic diameter, and anterior and posterior wall thickness at systole and diastole were recorded from M-mode images; cardiac output and stroke volume were calculated from bi-dimensional long-axis parasternal views from three consecutive cardiac cycles. Doppler waveforms in the umbilical vein and artery were obtained near the placental end of the umbilical cord. Area under the peak velocity-time curve, and R-R interval were measured from three consecutive cardiac cycles and the results were averaged. Umbilical venous and arterial diameters were measured from B-mode images. Mean velocity (MV) over the cardiac cycle was calculated by dividing the area under the peak velocity-time curve by the R-R interval. A parabolic blood velocity distribution was assumed so that umbilical venous and arterial blood flows were determined by the formula: F= ½ п MV (D/2)2 (where MV = mean peak velocity (cm/s); D = diameter (cm); F = blood flow (ml/min)).

### Cardiomyocyte isolation

The isolation of myocytes was performed as previously described^44^. Briefly, hearts were perfused with Ca^2+^-free bicarbonate buffer containing 120 mM NaCl, 5.4 mM KCl, 1.2 mM MgSO_4_, 1.2 mM NaH_2_PO_4_, 5.6 mM glucose, 20 mM NaHCO_3_, 20 mM 2,3 butanedione monoxime (Sigma-Aldrich, St. Louis, MO), and 5 mM taurine (Sigma-Aldrich), gassed with 95%O_2_/5% CO_2_, followed by enzymatic digestion with collagenase type II (1 mg/mL) (Worthington, Lakewood, NJ) and protease type XIV (0.1 mg/mL; Sigma-Aldrich). Cardiomyocytes were obtained from digested hearts followed by mechanical disruption, filtration, centrifugation, and resuspension in a Tyrode solution containing 0.125 mM CaCl_2_ Tyrode buffer containing 144 mM NaCl, 1 mM MgCl_2_, 10 mM Hepes, 1.2 mM NaH_2_PO_4_, 5.6 mM glucose, and 5 mM KCl, adjusted to pH 7.4 with NaOH.

### ROS by DCF, NO by DAF, Peroxynitrite by DHR 123

ROS, intracellular nitric oxide (NO) and peroxynitrite (ONOO‒) were measured by epifluorescence using 2’,7’-dichlorodihydrofluoresceine (H_2_DCF‒DA, 10 μM; Molecular Probes), 4,5-diaminofluorescein (DAF-2DA, 10 mM; Cayman Chemical Co., Ann Arbor, MI) and dihydrorhodamine 123 (DHR 123, 25 mM; Sigma-Aldrich), respectively. Briefly, fresh isolated mouse cardiomyocytes were placed in the chamber of an IonOptix spectrofluorometer and the background fluorescence (F_*0*_) was acquired with an excitation wavelength of 488 nm and emission fluorescence collected at 510 ± 15 nm. Cardiomyocytes were incubated for 40 min at room temperature (23°C) with H_2_DCF‒DA or DAF-2DA or 20 min with DHR 123 and washed by superfusing fresh Tyrode (1.8 mM CaCl_2_) solution for 10 minutes. Fluorescence (F) was acquired at 37°C every 1 minute for 10 minutes. Myocytes were stimulated at 1 Hz during the 10-minute experiment. ROS, NO or ONOO‒ levels were expressed as:

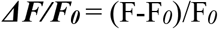

### Superoxide dismutase measurement

Sample preparation: frozen placentas were grinded in a Dounce homogenizer on liquid nitrogen. Then, pulverized tissue was homogenized in ice-cold PBS (10 μL per mg of tissue). The suspension was strained through a 250 μm-pore mesh. Then, samples were assessed for protein content by BCA (Pierce, Thermo Scientific) and diluted 1:10 in PBS for superoxide measurement. Superoxide was assessed by lucigenin-enhanced chemiluminescence.

Superoxide dismutase (SOD) assay: a superoxide-generator system (xanthine - xanthine oxidase (XO)) was used. XO (25 mU/mL) plus 5 μL of [SOD-containing] samples and 94.5 μL of PBS were added to the multiwell plate (in duplicate). Then, 100 μL of PBS containing 5 μM lucigenin (Enzo Life Sciences) and 200 μM xanthine (final concentration 100 μM, Sigma-Aldrich), were added. For positive control, a mix of 1 μL of 4 U/μL SOD (Sigma-Aldrich) with 99 μL of PBS was used. For superoxide generation control, 0.5 μL of 10 U/mL XO (Sigma-Aldrich) plus 99.5 μL of PBS (25 mU/mL final activity) was used.

Luminescence was acquired for 10 minutes every 30 seconds and the integrated luminescence units (LU) were used for calculations.

### Ascorbate treatment

Ascorbate was given in the drinking water starting from day 0.5 (time of plug detection). The water was changed every two days. In isolated CM, ascorbate (0.1 mM, 0.5 mM, or 1 mM) was incubated for 30 minutes prior to the start of the epifluorescence experiments.

### Tissue preparation and Histology and immunohistochemistry

At the end of the study, maternal organs (heart, kidney) and placenta were harvested, weighed, and processed for further analysis. Tissues were either flash-frozen in liquid nitrogen for total RNA isolation and protein analysis, while some tissues were fixed with 10% formalin for histology. Slides were stained with H&E and Masson’s trichrome staining for heart and kidney and PAS staining for kidney. Glomerular size was quantified using NIH Image J.

### Scanning Electron Microscope of the kidney

Tissue was fixed in 2% glutaraldehyde in 0.05M phosphate buffer and 100mM sucrose, post-fixed overnight in 1% osmium tetroxide in 0.1M phosphate buffer, dehydrated through a series of cold graded ethanols, and embedded in a mixture of EM-bed/Araldite (Electron Microscopy Sciences). 1μm thick sections were stained with Richardson’s stain for observation under a light microscope. 100nM sections were cut on a Leica Ultracut-R ultramicrotome and stained with uranyl acetate and lead citrate. The grids were viewed t 80 kV in a Philips CM-10 transmission electron microscope and images captured by a Gatan ES1000W digital camera. N=2 samples were examined per each group at 17.5 d of gestation.

### Isolectin immunofluorescence for paraffin-embedded tissues and analysis of placental capillary density

Paraffin placental sections were deparaffinized and rehydrated by immersion in xylene followed by a graded series of ethanol. Antigen retrieval was performed by a heat-induced method with citrate buffer (Dako, Carpinteria, CA). The slides were then blocked for 1 hour in 10% normal donkey serum to reduce background. Sections were then incubated with DyLight 594-GSL I-isolectin B4 (Vector Laboratories, Burlingame, CA) primary antibody for 1 hr. at 37°C. After washing with PBS, nuclei were counterstained with DAPI (Invitrogen, Carlsbad, CA). A stereological grid consisting of crosses was superimposed on images of placental sections stained with isolectin. The relative capillary density in the tissue was calculated on each section by dividing the number of crosses falling on capillary structure by the total number of points falling on the sampling area using Image J. For each placenta, four sections were analyzed.

### GSNOR activity in the mouse heart and mouse placenta

Heart and placental homogenate (100 μg/ml) were incubated with Tris-HCl (2mM, pH 8.0), EDTA (0.5 mM) and NADH (200 μM). The reaction was started by adding GSNO (400 μM) and activity was measured as GSNO-dependent NADH consumption at absorbance of 340 nm for 5 minutes in the mouse tissue.

### Protein immunoanalysis

Samples were electrophoresed using a NuPAGE 10% Bis-Tris gel (Invitrogen) and transferred to PVDF membranes (Bio-Rad Laboratories). Immunoblot detection was performed for VEGF (ab46154, 1:1000; Abcam, Cambridge, MA), VEGFR2 (55B11, 1:1000;Cell Signaling, Danvers, MA), eNOS (1:1000; BD Bioscience, BD Bioscience, San Jose, CA) in mouse tissue, B-actin (4957S, 1:1000, Cell Signaling, Danvers, MA), GAPDH (G8795, 1:1000, Sigma, St Louis, MO) and subsequent reaction with a goat anti-rabbit horseradish peroxidase-conjugated antibody (1:1000; Cell Signaling). Then, membranes were developed by enhanced chemiluminescence (Super Signal West Pico, Thermo Scientific, Hampton, NH) and analyzed by the QuantityOne software (Bio-Rad, Hercules, CA).

### VEGF nitrosylation

To assess S-nitrosylation, biotin-switch assay was used following methods described. Hearts were homogenized in HEN buffer (250 mM Hepes (pH 7.7), 1 mM EDTA, and 0.1 mM neocuproine). Free cysteine (Cys) residues were blocked with S-methyl methanethiosulfonate (MMTS) and labeled with N-[6-(biotinamido)hexyl]-3′-(2′-pyridyldithio) propionamide (HPDP-biotin) with or without sodium ascorbate. Biotinylated VEGF was individually immunoprecipitated with protein G-Sepharose beads, electrophoretically resolved, and immunoblotted with anti-biotin antibody. Blotted membranes were re-probed with related antibody for detection of protein load.

### RNA preparation and quantitative real-time PCR

Total RNA was extracted from tissues using the TRIzol method, and then reverse transcribed to complementary DNA using High-Capacity cDNA Reverse Transcription Kits (Applied Biosystems, USA) according to the manufacturer’s protocol. The quantitative RT-PCR for indicated genes was performed in TaqMan Universal PCR Master Mix (Applied Biosystems, USA). Quantitation of mRNAs was performed using Applied Biosystems™ TaqMan™ Gene Expression Assays according to the manufacturer’s protocol. Samples were analyzed using the BIORAD sequence detection system. All PCRs were performed in triplicate, and the specificity of the reaction was determined by melting curve analysis at the dissociation stage. The relative quantitative method was used for the quantitative analysis. The calibrator was the averaged ΔCt from the untreated cells. The endogenous control was glyceraldehyde 3-phosphate dehydrogenase (GAPDH).

### Mass Spectrometry Sample Preparation

All blocking and labeling steps were performed protected from light. Frozen placentas were individual minced in 0.9ml of cold homogenization buffer (PEN: PBS pH 8.0, 1mM ETDA, 0.1mM Neocupine; supplemented with 20 mM n-ethylmaleimide (NEM)). The tissue was then disrupted in 1ml mixer mill (Mixer Mill MM 400, retsch.com)) for 5 min and then subjected to probe sonication. Samples were adjusted to 2.5% SDS and clarified by centrifugation for 5 mins at 2000xg. The resulting supernatant was incubated for 10 mins at 50°C to completing the blocking step. The unreacted NEM was removed using a Zeba sin column (ThermoFisher Scientific) equilibrated with PEN buffer supplemented with 0.5% SDS. Each sample was divided and labeled with either 1 mM biotin-HPDP (ThermoFisher Scientific) or 0.3 mM iodoTMT^6^ (ThermoFisher Scientific) in the presence of 5 mM sodium ascorbate. As a labeling control, five pooled samples consisting of one replicate from each of the biological samples were prepared and reacted with each label in the absence of ascorbate. Samples were incubated for 1 hr. (HPDP) or 2 hr. (iodoTMT^6^) at 37°C. Excess label was removed from the HPDP treated samples by adding 2 volumes of cold acetone and incubating for 20 min at −20°C. Precipitated protein was pelleted by centrifugation and washed with 2 additional volumes of cold acetone. The pellets were resuspended 200 μl of PBS containing 1% (w/v) SDS aided by sonication. The excess IodoTMT^6^ label was removed by adding 5 volumes of cold acetone and precipitating as above. Samples were resuspended in 600 μl of PBS containing 1% (w/v) SDS aided by sonication. iodoTMT^6^ samples were then further reduced and alkylated using DTT and iodacetamide. Residual reagents were removed by Zeba spin column equilibrated with PBS. The protein concentration of each labeled sample was determined by BCA assay.

For HPDP labeled samples, 500 μg was digested overnight using 0.02 μg trypsin/μg of protein (Promega). iodoTMT^6^ labeled samples were combined according to the label’s isotope. Five-plexes were prepared containing 350 μg of each of the different biological samples and the pooled control. An additional set of six-plexes was prepared containing 250 μg of each biological replicates and a pooled control. The mixtures were digested overnight with 0.02 μg trypsin/μg of protein. Digestions were halted with 0.25 mM PMSF. The resulting peptides were captured using either streptavidin (HPDP) or TMT affinity resin (iodoTMT^6^). Peptides were enriched, washed, and eluted according to the manufacture’s protocol or as described here (PMID: 21036925). In the case of HPDP, eluted peptides were further alkylated with iodoacetamide.

### Mass Spectrometry analysis

The resulting peptides were desalted using Oasis HLB μ-elution plates (Waters, Milford, MA). Samples were eluted with 300 μL of 50% ACN, 0.1% FA dried in speedvac, then resuspended in 0.1% FA for LC/MS/MS analysis. LC/MS/MS analysis was performed using an Ultimate 3000 nano LC (Thermo Scientific) connected to an Orbitrap LUMOS mass spectrometer (Thermo Scientific) equipped with an EasySpray ion source. Peptides were loaded onto a PepMap RSLC C18 column (2 μm, 100 Å, 75 μm i.d. x 250 mm, Thermo Scientific) using a flow rate of 300 nL/min for 15 min at 1% B (mobile phase A was 0.1% formic acid in water and mobile phase B was 0.1 % formic acid in acetonitrile) after which point they were separated with a linear gradient of 2-20%B for 90 minutes, 20-32%B for 20 min, 32-95%B for 2 min, holding at 95%B for 8 minutes and re-equilibrating at 1%B for 5 minutes. Each sample was followed by a blank injection to both clean the column and re-equilibrated at 1%B. MS1 scans were acquired at a resolution of 240,000 Hz from mass range 400-1600 m/z. For MS1 scans the AGC target was set to 4×10^5^ ions with a max fill time of 50 ms. MS2 spectra were acquired using the TopSpeed method with a total cycle time of 3 seconds and an AGC target of 1×10^4^ and a max fill time of 100 ms, and an isolation width of 1.6 Da in the quadrapole. Precursor ions were fragmented using HCD with normalized collision energy of 30% and analyzed using rapid scan rates in the ion trap. Monoisotopic precursor selection was enabled and only MS1 signals exceeding 5000 counts triggered the MS2 scans, with +1 and unassigned charge states not being selected for MS2 analysis. Dynamic exclusion was enabled with a repeat count of 1 and exclusion duration of 15 seconds.

### Mass Spectrometry Data Analysis

Raw data was searched using a uniprot reviewed mouse database (09/18) with the X!Tandem (PMID: 14976030) algorithm version 2013.06.15.1 and Comet (PMID:23148064) algorithm version 2014.02 rev.2 search engines with the following parameters: Full Trypsin cleavage allowing for up to 2 missed cleavages, variable modifications +16 Da on Methionine (Oxidation), +57 Da, +125 Da and, in the case of IodoTMT6 labeled samples, +329 on Cysteine (Carbamidomethylation, NEM, TMT). Mass tolerance of MS1 error of 10 ppm, MS2 error of 1 Da were used. The mass spectrometry proteomics data have been deposited to the ProteomeXchange Consortium via the PRIDE (PMID: 30395289) partner repository with the dataset identifier PXD012706 (Reviewer account details: Username; Password: MkOzBjKi).

Differences SNO modification for HPDP labeled samples were determined by label-free quantitation using MS1 extracted ion chromatograms in Skyline (version 4.1) using a dot product cut off 0.8 (PMID: 22454539). Signal for each modified cysteine was summed within a replicate and normalized against the average of the pooled controls for that site. For iodoTMT^6^ labeled samples, only peptides with an iprofit score greater than 0.95 were considered. Reporter ion intensities were determined using the Libra module of the trans-proteomic pipeline (PMID: 20101611). Replicates were normalized against the pooled control present in the 5 or 6 plex. For all analysis, replicates need to be at least 1.5-fold greater than control and present in at least 40% of the replicates per group. Bioinformatic analysis was performed using Ingenuity Pathway Analysis (QIAGEN Inc., https://www.qiagenbioinformatics.com/products/ingenuity-pathway-analysis) and using gene ontology annotations in the Uniprot database (uniprot.org).

String v11 software was used to create pathway analysis of the data shown in Figure S2^45^.

### Statistics

The results are expressed as mean ± SEM. Differences between groups were examined for statistical significance using Student’s T-test or 1-way or 2-way analysis of variance (ANOVA), with Newman-Keuls for multiple comparisons for post hoc analysis where appropriate. Results P<0.05 were considered significant. All analysis were performed using SPSS and Prism Statistical software.

### Study Approval

All animal care was carried out in accordance with approval by the Institutional Animal Care and Use Committee.

## ACKNOWLEDGEMENTS

This study was funded by R01 HL09489 and R01 HL137355 to JMH and by Canadian Institute of Health Research postdoctoral fellowship and American Heart Association Career Development Award (19CDA34660102) to SK. JMH is also supported by NIH grants R01 HL134558, R01 HL101110, and 5UM 1HL113460 and by the Starr and Soffer Family Foundations. We acknowledge Vania Almeida and the UM Transmission Electron Microscopy Core, Dr. Wen Ding for urine collection.

## DATA AND MATERIAL AVAILABILITY

All data is available in the main text or the supplementary materials.

## AUTHOR CONTRIBUTIONS

S.K. conceived, designed, and executed the experiments and wrote the manuscript. R.A.D., M.A.B., J.F., R.K.T., E.P., H.A. performed experiments. C.I.M, D.S., J.E.V. performed mass spectrometry experiment and analysis. W.B. assisted in manuscript writing. J.M.H. conceived of and designed experiments, co-wrote the manuscript, and provided funding.

## COMPETING INTEREST STATEMENT

JMH reported having a patent for cardiac cell-based therapy. He holds equity in Vestion Inc. and maintains a professional relationship with Vestion Inc. as a consultant and member of the Board of Directors and Scientific Advisory Board. JMH is the Chief Scientific Officer, a compensated consultant and advisory board member for Longeveron, and holds equity in Longeveron. JMH is also the co-inventor of intellectual property licensed to Longeveron. Longeveron LLC and Vestion Inc. did not participate in funding this work. None of the other authors have anything to report.

## Notes

### Competing Interest Statement

The authors have declared no competing interest.

### Summary of Updates

This versions includes updated figures and methods and discussion.

## REFERENCES

1. Sibai B, Dekker G and Kupferminc M. Pre-eclampsia. Lancet. 2005;365:785–99.

2. Ives CW, Sinkey R, Rajapreyar I, Tita ATN and Oparil S. Preeclampsia-Pathophysiology and Clinical Presentations: JACC State-of-the-Art Review. J Am Coll Cardiol. 2020;76:1690–1702.

3. Creanga AA, Syverson C, Seed K and Callaghan WM. Pregnancy-Related Mortality in the United States, 2011-2013. Obstetrics and gynecology. 2017;130:366–373.

4. Molina RL and Pace LE. A Renewed Focus on Maternal Health in the United States. N Engl J Med. 2017;377:1705–1707.

5. Al-Gubory KH, Fowler PA and Garrel C. The roles of cellular reactive oxygen species, oxidative stress and antioxidants in pregnancy outcomes. The international journal of biochemistry & cell biology. 2010;42:1634–50.

6. Tyurin VA, Liu SX, Tyurina YY, Sussman NB, Hubel CA, Roberts JM, Taylor RN and Kagan VE. Elevated levels of S-nitrosoalbumin in preeclampsia plasma. Circulation research. 2001;88:1210–5.

7. Gandley RE, Tyurin VA, Huang W, Arroyo A, Daftary A, Harger G, Jiang J, Pitt B, Taylor RN, Hubel CA and Kagan VE. S-nitrosoalbumin-mediated relaxation is enhanced by ascorbate and copper: effects in pregnancy and preeclampsia plasma. Hypertension. 2005;45:21–7.

8. Webster RP, Roberts VH and Myatt L. Protein nitration in placenta - functional significance. Placenta. 2008;29:985–94.

9. Stamler JS, Jaraki O, Osborne J, Simon DI, Keaney J, Vita J, Singel D, Valeri CR and Loscalzo J. Nitric oxide circulates in mammalian plasma primarily as an S-nitroso adduct of serum albumin. Proceedings of the National Academy of Sciences of the United States of America. 1992;89:7674–7.

10. Foster MW, Pawloski JR, Singel DJ and Stamler JS. Role of circulating S-nitrosothiols in control of blood pressure. Hypertension. 2005;45:15–7.

11. Harris LK, McCormick J, Cartwright JE, Whitley GS and Dash PR. S-nitrosylation of proteins at the leading edge of migrating trophoblasts by inducible nitric oxide synthase promotes trophoblast invasion. Experimental cell research. 2008;314:1765–76.

12. Iyer AK, Rojanasakul Y and Azad N. Nitrosothiol signaling and protein nitrosation in cell death. Nitric oxide : biology and chemistry / official journal of the Nitric Oxide Society. 2014;42:9–18.

13. Lima B, Forrester MT, Hess DT and Stamler JS. S-nitrosylation in cardiovascular signaling. Circ Res. 2010;106:633–46.

14. Liu L, Yan Y, Zeng M, Zhang J, Hanes MA, Ahearn G, McMahon TJ, Dickfeld T, Marshall HE, Que LG and Stamler JS. Essential roles of S-nitrosothiols in vascular homeostasis and endotoxic shock. Cell. 2004;116:617–28.

15. Beigi F, Gonzalez DR, Minhas KM, Sun QA, Foster MW, Khan SA, Treuer AV, Dulce RA, Harrison RW, Saraiva RM, Premer C, Schulman IH, Stamler JS and Hare JM. Dynamic denitrosylation via S-nitrosoglutathione reductase regulates cardiovascular function. Proceedings of the National Academy of Sciences of the United States of America. 2012;109:4314–9.

16. Hatzistergos KE, Paulino EC, Dulce RA, Takeuchi LM, Bellio MA, Kulandavelu S, Cao Y, Balkan W, Kanashiro-Takeuchi RM and Hare JM. S-Nitrosoglutathione Reductase Deficiency Enhances the Proliferative Expansion of Adult Heart Progenitors and Myocytes Post Myocardial Infarction. Journal of the American Heart Association. 2015;4.

17. Barouch LA, Harrison RW, Skaf MW, Rosas GO, Cappola TP, Kobeissi ZA, Hobai IA, Lemmon CA, Burnett AL, O’Rourke B, Rodriguez ER, Huang PL, Lima JA, Berkowitz DE and Hare JM. Nitric oxide regulates the heart by spatial confinement of nitric oxide synthase isoforms. Nature. 2002;416:337–9.

18. Wei W, Li B, Hanes MA, Kakar S, Chen X and Liu L. S-nitrosylation from GSNOR deficiency impairs DNA repair and promotes hepatocarcinogenesis. Sci Transl Med. 2010;2:19ra13.

19. Stillman IE and Karumanchi SA. The glomerular injury of preeclampsia. Journal of the American Society of Nephrology : JASN. 2007;18:2281–4.

20. Kronborg C, Vittinghus E, Allen J and Knudsen UB. Excretion patterns of large and small proteins in pre-eclamptic pregnancies. Acta obstetricia et gynecologica Scandinavica. 2011;90:897–902.

21. Novelli GP, Valensise H, Vasapollo B, Larciprete G, Altomare F, Di Pierro G, Casalino B, Galante A and Arduini D. Left ventricular concentric geometry as a risk factor in gestational hypertension. Hypertension. 2003;41:469–75.

22. Schiattarella GG, Altamirano F, Tong D, French KM, Villalobos E, Kim SY, Luo X, Jiang N, May HI, Wang ZV, Hill TM, Mammen PPA, Huang J, Lee DI, Hahn VS, Sharma K, Kass DA, Lavandero S, Gillette TG and Hill JA. Nitrosative stress drives heart failure with preserved ejection fraction. Nature. 2019;568:351–356.

23. Karsdorp VH, van Vugt JM, van Geijn HP, Kostense PJ, Arduini D, Montenegro N and Todros T. Clinical significance of absent or reversed end diastolic velocity waveforms in umbilical artery. Lancet. 1994;344:1664–8.

24. Groten T, Gebhard N, Kreienberg R, Schleussner E, Reister F and Huppertz B. Differential expression of VE-cadherin and VEGFR2 in placental syncytiotrophoblast during preeclampsia - New perspectives to explain the pathophysiology. Placenta. 2010;31:339–43.

25. Sanchez-Aranguren LC, Prada CE, Riano-Medina CE and Lopez M. Endothelial dysfunction and preeclampsia: role of oxidative stress. Frontiers in physiology. 2014;5:372.

26. Phipps EA, Thadhani R, Benzing T and Karumanchi SA. Pre-eclampsia: pathogenesis, novel diagnostics and therapies. Nature reviews Nephrology. 2019;15:275–289.

27. Yang X, Guo L, Li H, Chen X and Tong X. Analysis of the original causes of placental oxidative stress in normal pregnancy and pre-eclampsia: a hypothesis. The journal of maternal-fetal & neonatal medicine : the official journal of the European Association of Perinatal Medicine, the Federation of Asia and Oceania Perinatal Societies, the International Society of Perinatal Obstet. 2012;25:884–8.

28. Kohr MJ, Sun J, Aponte A, Wang G, Gucek M, Murphy E and Steenbergen C. Simultaneous measurement of protein oxidation and S-nitrosylation during preconditioning and ischemia/reperfusion injury with resin-assisted capture. Circulation research. 2011;108:418–26.

29. Xu L, Eu JP, Meissner G and Stamler JS. Activation of the cardiac calcium release channel (ryanodine receptor) by poly-S-nitrosylation. Science. 1998;279:234–7.

30. Hubel CA, Kagan VE, Kisin ER, McLaughlin MK and Roberts JM. Increased ascorbate radical formation and ascorbate depletion in plasma from women with preeclampsia: implications for oxidative stress. Free radical biology & medicine. 1997;23:597–609.

31. Xu A, Vita JA and Keaney JF, Jr. Ascorbic acid and glutathione modulate the biological activity of S-nitrosoglutathione. Hypertension. 2000;36:291–5.

32. Chen X, Touyz RM, Park JB and Schiffrin EL. Antioxidant effects of vitamins C and E are associated with altered activation of vascular NADPH oxidase and superoxide dismutase in stroke-prone SHR. Hypertension. 2001;38:606–11.

33. Groebler LK, Wang XS, Kim HB, Shanu A, Hossain F, McMahon AC and Witting PK. Cosupplementation with a synthetic, lipid-soluble polyphenol and vitamin C inhibits oxidative damage and improves vascular function yet does not inhibit acute renal injury in an animal model of rhabdomyolysis. Free radical biology & medicine. 2012;52:1918–28.

34. Chung HS, Murray CI and Van Eyk JE. A Proteomics Workflow for Dual Labeling Biotin Switch Assay to Detect and Quantify Protein S-Nitroylation. Methods Mol Biol. 2018;1747:89–101.

35. Forrester MT, Foster MW and Stamler JS. Assessment and application of the biotin switch technique for examining protein S-nitrosylation under conditions of pharmacologically induced oxidative stress. The Journal of biological chemistry. 2007;282:13977–83.

36. Massy ZA, Fumeron C, Borderie D, Tuppin P, Nguyen-Khoa T, Benoit MO, Jacquot C, Buisson C, Drueke TB, Ekindjian OG, Lacour B and Iliou MC. Increased pasma S-nitrosothiol concentrations predict cardiovascular outcomes among patients with end-stage renal disease: a prospective study. Journal of the American Society of Nephrology : JASN. 2004;15:470–6.

37. Raijmakers MT, Zusterzeel PL, Steegers EA, Hectors MP, Demacker PN and Peters WH. Plasma thiol status in preeclampsia. Obstetrics and gynecology. 2000;95:180–4.

38. Pawloski JR, Hess DT and Stamler JS. Export by red blood cells of nitric oxide bioactivity. Nature. 2001;409:622–6.

39. Hlaing KH and Clement MV. Formation of protein S-nitrosylation by reactive oxygen species. Free radical research. 2014;48:996–1010.

40. McCance DR, Holmes VA, Maresh MJ, Patterson CC, Walker JD, Pearson DW, Young IS, Diabetes and Pre-eclampsia Intervention Trial Study G. Vitamins C and E for prevention of pre-eclampsia in women with type 1 diabetes (DAPIT): a randomised placebo-controlled trial. Lancet. 2010;376:259–66.

41. Poston L, Briley AL, Seed PT, Kelly FJ, Shennan AH and Vitamins in Pre-eclampsia Trial C. Vitamin C and vitamin E in pregnant women at risk for pre-eclampsia (VIP trial): randomised placebo-controlled trial. Lancet. 2006;367:1145–54.

42. Roberts JM, Myatt L, Spong CY, Thom EA, Hauth JC, Leveno KJ, Pearson GD, Wapner RJ, Varner MW, Thorp JM, Jr., Mercer BM, Peaceman AM, Ramin SM, Carpenter MW, Samuels P, Sciscione A, Harper M, Smith WJ, Saade G, Sorokin Y, Anderson GB, Eunice Kennedy Shriver National Institute of Child H and Human Development Maternal-Fetal Medicine Units N. Vitamins C and E to prevent complications of pregnancy-associated hypertension. The New England journal of medicine. 2010;362:1282–91.

43. Roberts JM. The perplexing pregnancy disorder preeclampsia: what next? Physiological genomics. 2018;50:459–467.

44. Khan SA, Skaf MW, Harrison RW, Lee K, Minhas KM, Kumar A, Fradley M, Shoukas AA, Berkowitz DE and Hare JM. Nitric oxide regulation of myocardial contractility and calcium cycling: independent impact of neuronal and endothelial nitric oxide synthases. Circulation research. 2003;92:1322–9.

45. Szklarczyk D, Gable AL, Lyon D, Junge A, Wyder S, Huerta-Cepas J, Simonovic M, Doncheva NT, Morris JH, Bork P, Jensen LJ and Mering CV. STRING v11: protein-protein association networks with increased coverage, supporting functional discovery in genome-wide experimental datasets. Nucleic acids research. 2019;47:D607–D613.

