## Supplement Figures and Tables for "S-nitrosoglutathione reductase deficiency causes aberrant placental S-nitrosylation and preeclampsia"

### Title: GSNOR deficiency causes aberrant placental S-nitrosylation and preeclampsia.

**This PDF file includes:**

Figs. S1 to S2
Tables S1 to S4

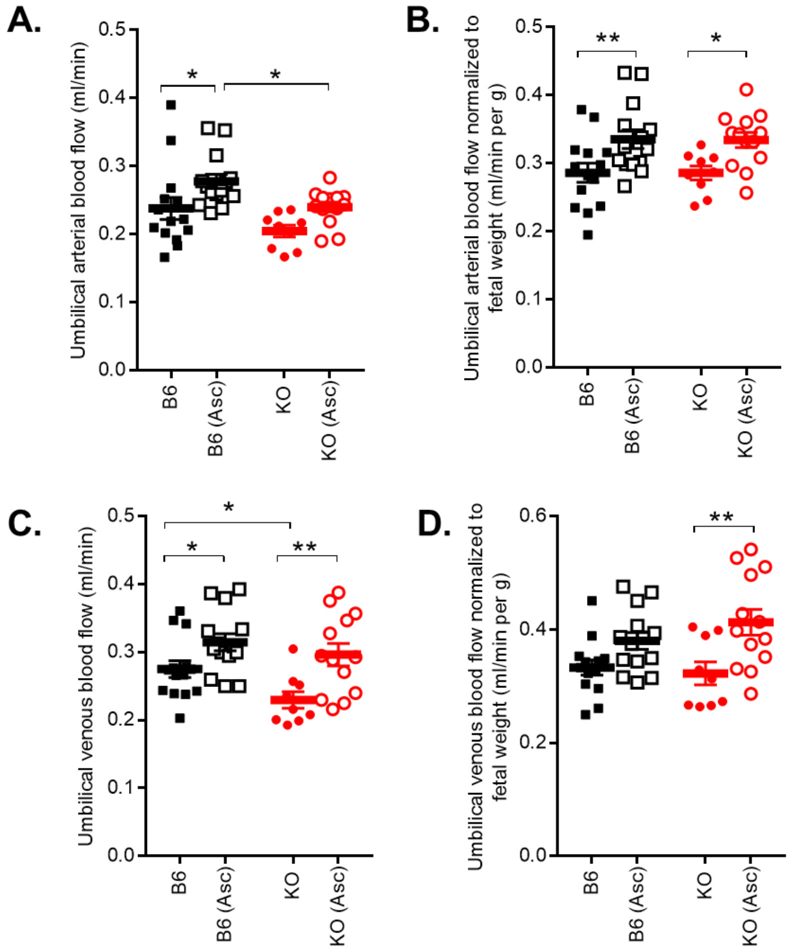

**Fig. S1:** Umbilical arterial and venous blood flow (**A, C**) and blood flow normalized to fetal weight **(B, D**) were determined using micro-ultrasound in isoflurane-anesthetized embryos on day 17.5 of gestation in C57Bl/6J (B6), GSNOR$^{-/-}$ (KO), and in mice treated with ascorbate (Asc). Umbilical venous blood flow was significantly lower in KO fetuses at 17.5 d of gestation and was rescued with Asc treatment. Results are shown as mean ± SEM. *P<0.05, **P<0.01. Two-way ANOVA with Newman-Keuls for post hoc analysis.

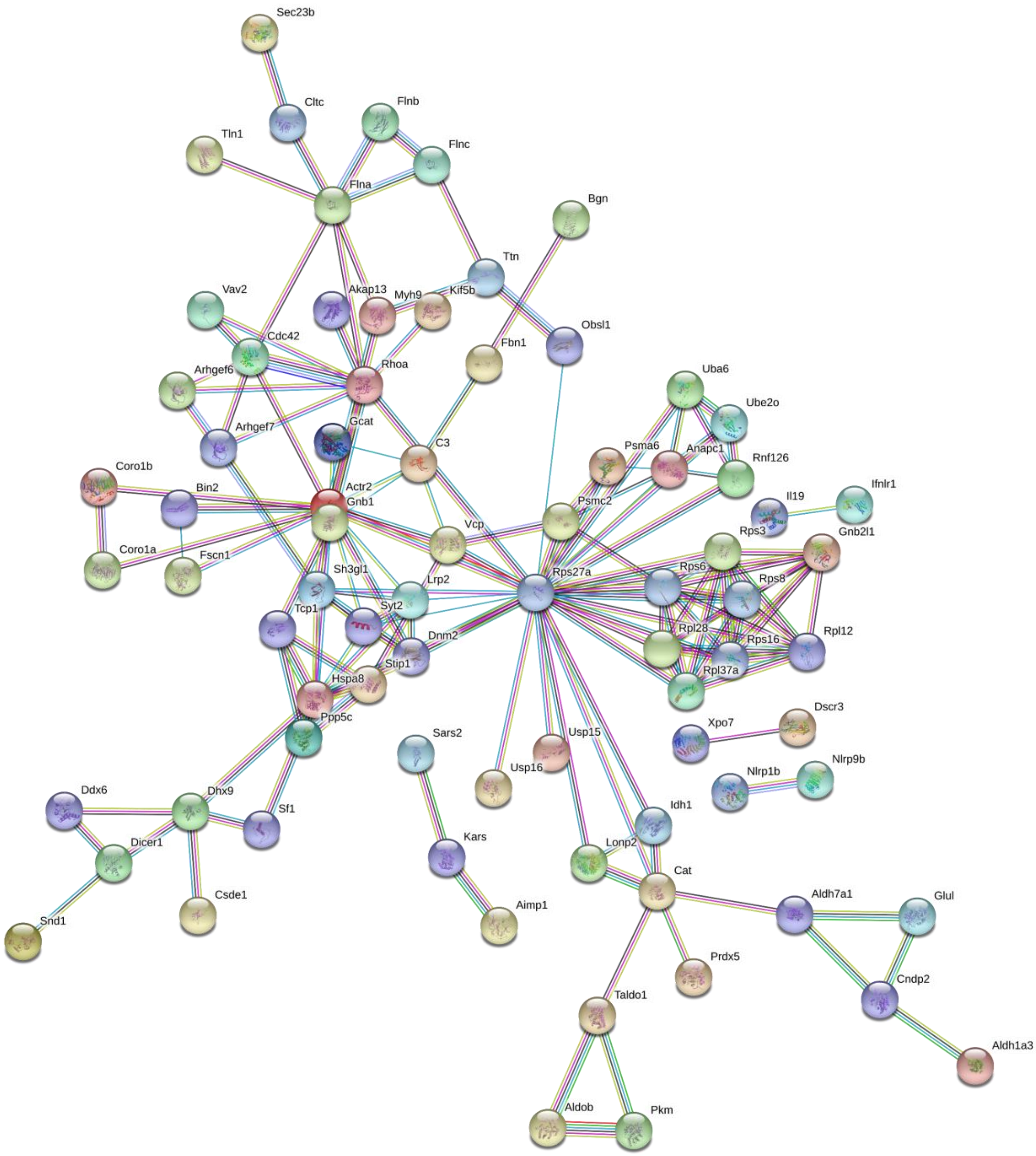

**Fig. S2.** Pathway analysis of list of proteins nitrosylated in GSNOR−/− placentas as compared to B6 placentas determined using dual-labelling mass spectrometry analysis was created using String-DB. All SNO-proteins were detected in at least 2 of 5 placentas/group. The number in the observation column is the number of placentas that showed expression of that particular SNO-protein for that group. These proteins correlate to data shown in Table S2

Table S1: List of proteins nitrosylated in GSNOR$^{-/-}$ placentas as compared to B6 placentas determined using dual-labelling mass spectrometry analysis.

Table S2: List of proteins nitrosylated in GSNOR$^{-/-}$ placentas as compared to B6 placentas and B6- and GSNOR$^{-/-}$ placentas treated with ascorbate.

Table S3: List of peptides identified using mass spectrometry analysis.

| Table S1: List of proteins nitrosylated in GSNOR$^{-/-}$ placentas as compared to B6 placentas determined using dual-labelling mass spectrometry analysis. All SNOylated proteins were detected in at least 2 of 5 placentas/group. The number in the observation column is the number of placentas that showed expression of that particular SNOylated protein for that particular group. | | | | | | | |
|---|---|---|---|---|---|---|---|
| label | Uniprot Accession | SNO site | Protein name | Log2 fold change vs background B6 | observations | Log2 fold change vs background GSNOR$^{-/-}$ | observations |
| HPDP | Q99L04 | C10 | Dehydrogenase/reductase SDR family member 1 | | 0 | 23.05415 | 2 |
| HPDP | Q99MN1 | C432 | Lysine--tRNA ligase | | 0 | 22.7331744 | 2 |
| HPDP | Q8C7R4 | C298 | Ubiquitin-like modifier-activating enzyme 6 (Ubiquitin-activating enzyme 6) | | 0 | 22.10295762 | 2 |
| HPDP | Q8VC70 | C217 | RNA-binding motif, single-stranded-interacting | | 0 | 21.71700633 | 2 |
| HPDP | Q9D1I5 | C168 | Methylmalonyl-CoA epimerase, mitochondrial | | 0 | 21.24558726 | 2 |
| HPDP | Q9JIG4 | C419 | Protein phosphatase 1 regulatory subunit 3F (R3F) | | 0 | 20.95864848 | 2 |
| HPDP | Q9EPK7 | C43 | Exportin-7 (Exp7) (Ran-binding protein 16) | | 0 | 20.94875308 | 2 |
| HPDP | Q9QUI0 | C164 | Transforming protein RhoA | | 0 | 20.91330254 | 2 |
| HPDP | Q64514 | C967 | Tripeptidyl-peptidase 2 (TPP-2) | | 0 | 20.80956501 | 2 |
| HPDP | O70400 | C73 | PDZ and LIM domain protein 1 (C-terminal LIM domain protein 1) (Elfin) (LIM domain protein CLP- | | 0 | 20.77362179 | 2 |
| HPDP | P24270 | C232 | Catalase | | 0 | 20.71680767 | 2 |
| HPDP | Q8BY89 | C401 | Choline transporter-like protein 2 (Solute carrier family 44 member 2) | | 0 | 20.70444271 | 2 |
| HPDP | P61982 | C112 | 14-3-3 protein gamma [Cleaved into: 14-3-3 protein gamma, N-terminally processed] | | 0 | 20.6712323 | 2 |
| HPDP | Q8VDD5 | C740 | Myosin-9 (Cellular myosin heavy chain, type A) (Myosin heavy chain 9) (Myosin heavy chain, non-muscle IIa) (Non-muscle myosin heavy chain A) (NMMHC-A) (Non-muscle myosin heavy chain IIa) (NMMHC II-a) (NMMHC-IIA) | | 0 | 20.40955957 | 2 |

| | | | | | | | |
|---|---|---|---|---|---|---|---|
| HPDP | P23198 | C177 | Chromobox protein homolog 3 (Heterochromatin protein 1 homolog gamma) (HP1 gamma) (M32) (Modifier 2 protein) | | 0 | 20.32635422 | 2 |
| HPDP | O88844 | C73 | Isocitrate dehydrogenase [NADP] cytoplasmic (IDH) | | 0 | 20.28660949 | 2 |
| HPDP | Q7TSI1 | C464 | Pleckstrin homology domain-containing family M member 1 (PH domain-containing family M member [illegible] | | 0 | 20.27430909 | 2 |
| HPDP | Q62419 | C277 | Endophilin-A2 (Endophilin-2) (SH3 domain protein 2B) (SH3 domain-containing GRB2-like protein 1) [illegible] | | 0 | 20.1597845 | 2 |
| HPDP | Q9Z2W0 | C411 | Aspartyl aminopeptidase | | 0 | 20.08429446 | 2 |
| HPDP | Q9JJL8 | C425 | Serine--tRNA ligase, mitochondrial | | 0 | 20.04438709 | 2 |
| HPDP | P21981 | C553 | Protein-glutamine gamma-glutamyltransferase 2 | | 0 | 19.98447735 | 2 |
| HPDP | Q91Y97 | C158 | Fructose-bisphosphate aldolase B | | 0 | 19.87182726 | 2 |
| HPDP | Q9DBN5 | C405 | Lon protease homolog 2, peroxisomal | | 0 | 19.84850213 | 2 |
| HPDP | Q8VC03 | C421 | Echinoderm microtubule-associated protein-like 3 (EMAP-3) | | 0 | 19.83187399 | 2 |
| HPDP | P62242 | C100 | 40S ribosomal protein S8 | | 0 | 19.81349615 | 2 |
| HPDP | A2ASS6 | C29432 | Titin | | 0 | 19.79460004 | 2 |
| HPDP | Q63ZW7 | C1406 | InaD-like protein (Inadl protein) (Channel-interacting PDZ domain-containing protein) (Pals1-associated tight junction protein) (Protein associated to tight | | 0 | 19.70935729 | 2 |
| HPDP | Q9CPV4 | C45 | Glyoxalase domain-containing protein 4 | | 0 | 19.70621887 | 2 |
| HPDP | Q68FD5 | C1266 | Clathrin heavy chain 1 | | 0 | 19.69269741 | 2 |
| HPDP | P54823 | C390 | Probable ATP-dependent RNA helicase DDX6 | | 0 | 19.49673091 | 2 |
| HPDP | Q91YL2 | C32 | E3 ubiquitin-protein ligase RNF126 | | 0 | 19.48361501 | 2 |
| HPDP | A6H8H2 | C1083 | DENN domain-containing protein 4C | | 0 | 19.41978765 | 3 |
| HPDP | Q01853 | C69,C77 | Transitional endoplasmic reticulum ATPase (TER ATPase) | | 0 | 19.40789288 | 2 |
| HPDP | B7ZMP1 | C491 | Xaa-Pro aminopeptidase 3 (X-Pro aminopeptidase 3) | | 0 | 19.29977564 | 2 |
| HPDP | A2ASS6 | C21780 | Titin | | 0 | 19.12293298 | 2 |
| HPDP | Q9ES28 | C427 | Rho guanine nucleotide exchange factor 7 (Beta-Pix) (PAK-interacting exchange factor beta) (p85SPR) | | 0 | 19.12161126 | 2 |

| | | | | | | | |
|---|---|---|---|---|---|---|---|
| HPDP | A2ARV4 | C2518 | Low-density lipoprotein receptor-related protein 2 (LRP-2) (Glycoprotein 330) (gp330) (Megalin) | | 0 | 19.03682762 | 2 |
| HPDP | Q6ZPJ3 | C365 | (E3-independent) E2 ubiquitin-conjugating enzyme UBE2O | | 0 | 18.97268389 | 3 |
| HPDP | Q61655 | C392 | ATP-dependent RNA helicase DDX19A | | 0 | 18.88880189 | 2 |
| HPDP | P14824 | C669 | Annexin A6 (67 kDa calelectrin) (Annexin VI) (Annexin-6) (Calphobindin-II) (CPB-II) (Chromobindin-20) (Lipocortin VI) (Protein III) (p68) [illegible] | | 0 | 18.8218629 | 2 |
| HPDP | Q5HZI1 | C823 | Microtubule-associated tumor suppressor 1 homolog (AT2 receptor-binding protein) (Angiotensin-II type 2 receptor-interacting protein) (Coiled-coiled tumor suppressor gene 1 protein) (Mitochondrial tumor suppressor 1 homolog) | | 0 | 18.76396509 | 2 |
| HPDP | P53996 | C141,C151 | Cellular nucleic acid-binding protein (CNBP) (Zinc finger protein 9) | | 0 | 18.75122145 | 2 |
| HPDP | Q921G6 | C454 | Leucine-rich repeat and calponin homology domain-containing protein 4 | | 0 | 18.56565726 | 4 |
| HPDP | P82343 | C250 | N-acylglucosamine 2-epimerase (AGE) | | 0 | 16.42343934 | 2 |
| HPDP | Q91W50 | C129 | Cold shock domain-containing protein E1 | | 0 | 16.0168855 | 2 |
| TMT | Q7TNJ0 | C89 | Dendritic cell-specific transmembrane protein (DC-STAMP) (mDC-STAMP) (Dendrocyte-expressed seven transmembrane protein) (Transmembrane 7 superfamily member 4) | | 0 | 13.72900332 | 4 |
| TMT | Q8R418 | C306 | Endoribonuclease Dicer | | 0 | 13.02362462 | 5 |
| TMT | Q5FW85 | C8 | Extracellular matrix protein 2 (Tenonectin) | | 0 | 13.00944639 | 4 |
| TMT | Q8BV57 | C6 | Soluble scavenger receptor cysteine-rich domain-containing protein SSC5D (Scavenger receptor cysteine-rich domain-containing protein LOC284297 [illegible] | | 0 | 12.76398811 | 5 |
| TMT | P70277 | C65 | Alpha-N-acetylgalactosaminide alpha-2,6-sialyltransferase 2 | | 0 | 12.6184686 | 5 |

| | | | | | | | |
|---|---|---|---|---|---|---|---|
| TMT | Q3UAW9 | C362 | Transcription factor IIIB 50 kDa subunit (B-related factor 2) (BRF-2) | | 0 | 12.585631 | 5 |
| TMT | Q8BLR9 | C236 | Hypoxia-inducible factor 1-alpha inhibitor | | 0 | 12.5665622 | 5 |
| TMT | Q9JMG4 | C8,C14 | Sodium/potassium-transporting ATPase subunit beta-1-interacting protein 4 (Na(+)/K(+)-transporting ATPase subunit beta-1-interacting protein 4) (Protein FAM77A) | | 0 | 12.39302865 | 5 |
| TMT | A1Z198 | C330 | NACHT, LRR and PYD domains-containing protein 1b allele 2 | | 0 | 12.36513178 | 5 |
| TMT | Q9DB60 | C44,C47 | Prostamide/prostaglandin F synthase (Prostamide/PG F synthase) (Prostamide/PGF synthase) | | 0 | 12.33380021 | 5 |
| TMT | P57110 | C677 | A disintegrin and metalloproteinase with thrombospondin motifs 8 (ADAM-TS 8) (ADAM- | | 0 | 12.30280006 | 5 |
| TMT | Q571F5 | C271 | SPRY domain-containing SOCS box protein 3 (SSB- | | 0 | 12.24478409 | 5 |
| TMT | O09118 | C17 | Netrin-1 | | 0 | 12.21630324 | 5 |
| TMT | Q8C0W1 | C818 | Ankyrin repeat and MYND domain-containing protein | | 0 | 12.20108475 | 5 |
| TMT | Q9DBP5 | C20 | UMP-CMP kinase | | 0 | 12.14459788 | 5 |
| TMT | O35963 | C48 | Ras-related protein Rab-33B | | 0 | 12.08737651 | 5 |
| TMT | A1L0T3 | C138 | Scavenger receptor cysteine-rich domain-containing group B protein (Four scavenger receptor cysteine-rich domains-containing protein) (S4D-SRCRB) | | 0 | 11.90176048 | 5 |
| TMT | Q8CGM1 | C1034 | Adhesion G protein-coupled receptor B2 (Brain-specific angiogenesis inhibitor 2) | | 0 | 11.8771311 | 5 |
| TMT | Q9WTN3 | C738,C753 | Sterol regulatory element-binding protein 1 (SREBP-1) (Sterol regulatory element-binding transcription factor 1) [Cleaved into: Processed sterol regulatory element-binding protein 1] | | 0 | 11.804481 | 5 |
| TMT | Q8CJ70 | C5,C18 | Interleukin-19 (IL-19) | | 0 | 11.69925873 | 5 |
| TMT | Q6DFV8 | C217 | von Willebrand factor D and EGF domain-containing protein | | 0 | 11.67417859 | 4 |

| | | | | | | | |
|---|---|---|---|---|---|---|---|
| TMT | Q8CGK5 | C81 | Interferon lambda receptor 1 (IFN-lambda R1) (Cytokine receptor class-II member 12) (Cytokine receptor family 2 member 12) (CRF2-12) (Interleukin-28 receptor subunit alpha) (IL-28 receptor subunit alpha) (IL-28R-alpha) (IL-28RA) | | 0 | 11.61269071 | 5 |
| TMT | Q4VAE3 | C31 | Transmembrane protein 65 | | 0 | 11.41109714 | 5 |
| TMT | Q9JLL3 | C25 | Tumor necrosis factor receptor superfamily member 19 (TRADE) (Toxicity and JNK inducer) | | 0 | 11.35771618 | 5 |
| TMT | Q8K400 | C293 | Syntaxin-binding protein 5 (Lethal(2) giant larvae protein homolog 3) (Tomosyn-1) | | 0 | 11.21114736 | 5 |
| TMT | Q9DC22 | C200,C211 | DDB1- and CUL4-associated factor 6 (IQ motif and WD repeat-containing protein 1) (Nuclear receptor interaction protein) (NRIP) | | 0 | 11.00286202 | 5 |
| HPDP | Q9WUM3 | C25 | Coronin-1B (Coronin-2) | | 0 | 7.079871538 | 2 |
| HPDP | Q8K274 | C11 | Ketosamine-3-kinase | | 0 | 5.271225169 | 2 |
| HPDP | P46471 | C389 | 26S proteasome regulatory subunit 7 (26S proteasome AAA-ATPase subunit RPT1) (Proteasome 26S subunit ATPase 2) (Protein MSS1) | | 0 | 5.062936789 | 2 |
| HPDP | P62754 | C12 | 40S ribosomal protein S6 (Phosphoprotein NP33) | | 0 | 4.714241907 | 2 |
| HPDP | Q8R5H1 | C264 | Ubiquitin carboxyl-terminal hydrolase 15 | | 0 | 4.569012373 | 2 |
| HPDP | Q9JJ28 | C1069 | Protein flightless-1 homolog | | 0 | 4.407051634 | 2 |
| HPDP | Q60992 | C196,C197 | Guanine nucleotide exchange factor VAV2 (VAV-2) | | 0 | 4.179681175 | 2 |
| HPDP | Q9D662 | C425 | Protein transport protein Sec23B (SEC23-related protein B) | | 0 | 3.985939588 | 2 |
| HPDP | O70439 | C28 | Syntaxin-7 | | 0 | 3.955148625 | 3 |
| HPDP | Q61768 | C632 | Kinesin-1 heavy chain (Conventional kinesin heavy chain) (Ubiquitous kinesin heavy chain) (UKHC) | | 0 | 3.760958689 | 2 |
| HPDP | Q99JY3 | C61 | GTPase IMAP family member 4 (Immunity-associated nucleotide 1 protein) (IAN-1) (Immunity-associated protein 4) | | 0 | 3.614276688 | 2 |
| HPDP | P99029 | C200 | Peroxiredoxin-5, mitochondrial | | 0 | 3.604855607 | 3 |

| | | | | | | | |
|---|---|---|---|---|---|---|---|
| HPDP | Q8BZB2 | C7 | Phosphopantothenoylcysteine decarboxylase (PPC- | | 0 | 3.603461526 | 2 |
| TMT | Q80VD1 | C52 | Protein FAM98B | | 0 | 3.259995258 | 5 |
| TMT | Q80XD8 | C9 | Proline-rich acidic protein 1 (Pregnancy-specific uterine protein) (Uterine-specific proline-rich acidic | | 0 | 3.172829061 | 5 |
| TMT | Q9ESD6 | C54 | CKLF-like MARVEL transmembrane domain-containing protein 7 (Chemokine-like factor superfamily member 7) (LNV) | | 0 | 2.984893682 | 5 |
| HPDP | Q9CYN2 | C26 | Signal peptidase complex subunit 2 | | 0 | 2.967611159 | 2 |
| HPDP | P14131 | C25 | 40S ribosomal protein S16 | | 0 | 2.96535426 | 2 |
| TMT | Q9JIP3 | C12 | Interleukin-17 receptor B (IL-17 receptor B) (IL-17RB) (IL-17 receptor homolog 1) (IL-17ER) (IL-17Rh1) (IL17Rh1) (Interleukin-17B receptor) (IL-17B receptor) | | 0 | 2.927411041 | 5 |
| TMT | Q8BKK5 | C263 | Zinc finger protein 689 | | 0 | 2.920765805 | 5 |
| HPDP | Q99LG0 | C24 | Ubiquitin carboxyl-terminal hydrolase 16 | | 0 | 2.800809396 | 2 |
| HPDP | Q9Z1Z0 | C802 | General vesicular transport factor p115 (Protein USO1 homolog) (Transcytosis-associated protein) (TAP) (Vesicle-docking protein) | | 0 | 2.748838009 | 2 |
| TMT | O08738 | C259 | Caspase-6 (CASP-6) | | 0 | 2.742606331 | 5 |
| TMT | O88282 | C381,C384 | B-cell CLL/lymphoma 6 member B protein (Bcl6-associated zinc finger protein) | | 0 | 2.722924669 | 5 |
| TMT | Q9Z0L3 | C371 | Otoconin-90 (Oc90) (Otoconin-95) (Oc95) | | 0 | 2.71298463 | 5 |
| TMT | E9PZZ1 | C653 | PR domain zinc finger protein 13 | | 0 | 2.547554782 | 4 |
| HPDP | Q8VHX6 | C2661 | Filamin-C (FLN-C) (ABP-280-like protein) (ABP-L) (Actin-binding-like protein) (Filamin-2) (Gamma- | | 0 | 2.489804487 | 2 |
| TMT | Q8BYA0 | C665 | Tubulin-specific chaperone D (Beta-tubulin cofactor D) (Tubulin-folding cofactor D) | | 0 | 2.48712789 | 5 |
| HPDP | P53996 | C120 | Cellular nucleic acid-binding protein (CNBP) (Zinc finger protein 9) | | 0 | 2.455112234 | 2 |
| HPDP | Q8CGB6 | C548 | Tensin-2 | | 0 | 2.399443288 | 3 |
| HPDP | P15105 | C183 | Glutamine synthetase (GS) | | 0 | 2.388341937 | 2 |

| | | | | | | | |
|---|---|---|---|---|---|---|---|
| HPDP | Q9CQ58 | C101 | Prolactin-8A9 (Placental prolactin-like protein C2) (PLP-C2) (PRL-like protein C2) (Prolactin-like protein C-beta) (PLP C-beta) | | 0 | 2.373532499 | 3 |
| TMT | Q60676 | C221 | Serine/threonine-protein phosphatase 5 (PP5) | | 0 | 2.35208233 | 5 |
| HPDP | Q80XN0 | C209 | D-beta-hydroxybutyrate dehydrogenase, mitochondrial | | 0 | 2.33471439 | 3 |
| HPDP | Q93092 | C250 | Transaldolase | | 0 | 2.325417107 | 2 |
| TMT | Q5F2L2 | C13 | Alpha-(1,3)-fucosyltransferase 10 | | 0 | 2.317276958 | 5 |
| TMT | O09008 | C18 | Beta-1,3-N-acetylglucosaminyltransferase manic fringe | | 0 | 2.305226515 | 4 |
| HPDP | P17742 | C67 | Peptidyl-prolyl cis-trans isomerase A (PPIase A) | | 0 | 2.291684824 | 2 |
| HPDP | A2ASS6 | C21834 | Titin | | 0 | 2.248744617 | 2 |
| HPDP | O08573 | C258 | Galectin-9 (Gal-9) | | 0 | 2.210136624 | 2 |
| HPDP | Q9D1Q6 | C92 | Endoplasmic reticulum resident protein 44 (ER protein 44) (ERp44) (Thioredoxin domain-containing [illegible] | | 0 | 2.147055114 | 2 |
| HPDP | P53995 | C988 | Anaphase-promoting complex subunit 1 (APC1) (Cyclosome subunit 1) (Mitotic checkpoint regulator) (Testis-specific gene 24 protein) | | 0 | 2.039981245 | 2 |
| HPDP | P70441 | C201 | Na(+)/H(+) exchange regulatory cofactor NHE-RF1 (NHERF-1) (Ezrin-radixin-moesin-binding phosphoprotein 50) (EBP50) (Regulatory cofactor of Na(+)/H(+) exchanger) (Sodium-hydrogen exchanger regulatory factor 1) (Solute carrier family 9 isoform A3 regulatory factor 1) | | 0 | 1.988263764 | 2 |
| TMT | Q9QXW9 | C209 | Large neutral amino acids transporter small subunit 2 (L-type amino acid transporter 2) (mLAT2) (Solute carrier family 7 member 8) | | 0 | 1.963679629 | 5 |
| HPDP | P60766 | C157 | Cell division control protein 42 homolog (G25K GTP-binding protein) | | 0 | 1.950445964 | 2 |
| TMT | Q66X22 | C891,C907,C908 | NACHT, LRR and PYD domains-containing protein 9B (NALP-delta) | | 0 | 1.935676951 | 4 |

| | | | | | | | |
|---|---|---|---|---|---|---|---|
| HPDP | P01027 | C559 | Complement C3 (HSE-MSF) [Cleaved into: Complement C3 beta chain; C3-beta-c (C3bc); Complement C3 alpha chain; C3a anaphylatoxin; Acylation stimulating protein (ASP) (C3adesArg); Complement C3b alpha' chain; Complement C3c alpha' chain fragment 1; Complement C3dg fragment; Complement C3g fragment; Complement C3d fragment; Complement C3f fragment; Complement C3c alpha' chain fragment 2] | | 0 | 1.93095258 | 2 |
| HPDP | O88986 | C26 | 2-amino-3-ketobutyrate coenzyme A ligase, mitochondrial (AKB ligase) | | 0 | 1.921218785 | 2 |
| HPDP | P35979 | C141 | 60S ribosomal protein L12 | | 0 | 1.912774567 | 2 |
| HPDP | P63017 | C603 | Heat shock cognate 71 kDa protein (Heat shock 70 kDa protein 8) | | 0 | 1.900311126 | 2 |
| TMT | Q8BLY7 | C180 | Hermansky-Pudlak syndrome 6 protein homolog (Ruby-eye protein) (Ru) | | 0 | 1.878707928 | 5 |
| HPDP | Q7TMW6 | C270 | Cytosolic iron-sulfur assembly component 3 (Cytosolic Fe-S cluster assembly factor NARFL) (Iron-only hydrogenase-like protein 1) (IOP1) (Nuclear prelamin A recognition factor-like protein) | | 0 | 1.847995032 | 2 |
| HPDP | Q60864 | C461 | Stress-induced-phosphoprotein 1 (STI1) (mSTI1) (Hsc70/Hsp90-organizing protein) (Hop) | | 0 | 1.832341765 | 2 |
| HPDP | A2ASS6 | C13473 | Titin | | 0 | 1.794763146 | 2 |
| TMT | Q8K4G1 | C1393,C1403 | Latent-transforming growth factor beta-binding protein 4 (LTBP-4) | | 0 | 1.71923574 | 5 |
| HPDP | O89053 | C24 | Coronin-1A (Coronin-like protein A) (Clipin-A) (Coronin-like protein p57) (Tryptophan aspartate-containing coat protein) (TACO) | | 0 | 1.710727777 | 2 |
| TMT | Q7TPG7 | C107 | Protein FAM19A2 (Chemokine-like protein TAFA-2) | | 0 | 1.706388912 | 5 |
| HPDP | Q9QUM9 | C167 | Proteasome subunit alpha type-6 | | 0 | 1.696998107 | 2 |
| HPDP | Q80X90 | C1434 | Filamin-B (FLN-B) (ABP-280-like protein) (Actin-binding-like protein) (Beta-filamin) | | 0 | 1.668503957 | 2 |

| | | | | | | | |
|---|---|---|---|---|---|---|---|
| HPDP | A2ASS6 | C33458 | Titin | | 0 | 1.640392354 | 2 |
| HPDP | Q61553 | C481 | Fascin (Singed-like protein) | | 0 | 1.636609875 | 2 |
| HPDP | P61514 | C48 | 60S ribosomal protein L37a | | 0 | 1.593487828 | 2 |
| HPDP | P61161 | C221 | Actin-related protein 2 (Actin-like protein 2) | | 0 | 1.554988721 | 2 |
| TMT | P46097 | C91 | Synaptotagmin-2 (Inositol polyphosphate-binding protein) (IP4-binding protein) (IP4BP) (Synaptotagmin II) (SytII) | | 0 | 1.554328226 | 5 |
| HPDP | Q8VDF3 | C347 | Death-associated protein kinase 2 (DAP kinase 2) | | 0 | 1.52416734 | 2 |
| HPDP | Q78PY7 | C152 | Staphylococcal nuclease domain-containing protein 1 | | 0 | 1.520546542 | 2 |
| HPDP | Q8BTM8 | C2582 | Filamin-A (FLN-A) (Actin-binding protein 280) (ABP-280) (Alpha-filamin) (Endothelial actin-binding protein) (Filamin-1) (Non-muscle filamin) | | 0 | 1.416572906 | 2 |
| HPDP | Q64514 | C150 | Tripeptidyl-peptidase 2 (TPP-2) | | 0 | 1.399757173 | 2 |
| HPDP | Q09324 | C100 | Beta-1,3-galactosyl-O-glycosyl-glycoprotein beta-1,6-N-acetylglucosaminyltransferase | | 0 | 1.397345074 | 2 |
| HPDP | Q9CY97 | C12 | RNA polymerase II subunit A C-terminal domain phosphatase SSU72 (CTD phosphatase SSU72) | | 0 | 1.39424147 | 2 |
| HPDP | P39054 | C607 | Dynamin-2 | | 0 | 1.374247771 | 2 |
| HPDP | P68040 | C153 | Receptor of activated protein C kinase 1 (12-3) (Guanine nucleotide-binding protein subunit beta-2-like 1) (Receptor for activated C kinase) (Receptor of activated protein kinase C 1) (p205) [Cleaved into: Receptor of activated protein C kinase 1, N-terminally processed (Guanine nucleotide-binding protein subunit beta-2-like 1, N-terminally processed)] | | 0 | 1.359971038 | 3 |
| HPDP | P26039 | C956 | Talin-1 | | 0 | 1.354647422 | 2 |
| HPDP | O70133 | C440 | ATP-dependent RNA helicase A | | 0 | 1.344512262 | 2 |
| HPDP | Q80X90 | C991 | Filamin-B (FLN-B) (ABP-280-like protein) (Actin-binding-like protein) (Beta-filamin) | | 0 | 1.337916373 | 2 |

| | | | | | | | |
|---|---|---|---|---|---|---|---|
| HPDP | Q99K30 | C546 | Epidermal growth factor receptor kinase substrate 8-like protein 2 (EPS8-like protein 2) (Epidermal growth factor receptor pathway substrate 8-related protein 2) (EPS8-related protein 2) | | 0 | 1.332404247 | 2 |
| HPDP | P62908 | C134 | 40S ribosomal protein S3 | | 0 | 1.319576402 | 2 |
| HPDP | E9Q394 | C609 | A-kinase anchor protein 13 (AKAP-13) (AKAP-Lbc) | | 0 | 1.277028431 | 3 |
| TMT | P02798 | C33 | Metallothionein-2 (MT-2) (Metallothionein-II) (MT- | | 0 | 1.266697299 | 4 |
| HPDP | P62908 | C119 | 40S ribosomal protein S3 | | 0 | 1.254357417 | 2 |
| HPDP | Q99LR1 | C40 | Monoacylglycerol lipase ABHD12 | | 0 | 1.23650774 | 2 |
| HPDP | P46471 | C377 | 26S proteasome regulatory subunit 7 (26S proteasome AAA-ATPase subunit RPT1) (Proteasome 26S subunit ATPase 2) (Protein MSS1) | | 0 | 1.228249064 | 2 |
| HPDP | P28653 | C77 | Biglycan (Bone/cartilage proteoglycan I) (PG-S1) | | 0 | 1.222294589 | 2 |
| HPDP | O35075 | C243 | Down syndrome critical region protein 3 homolog (Down syndrome critical region protein A homolog) | | 0 | 1.212332349 | 2 |
| HPDP | P61161 | C11 | Actin-related protein 2 (Actin-like protein 2) | | 0 | 1.199108066 | 2 |
| HPDP | P01837 | C106 | Immunoglobulin kappa constant (Ig kappa chain C region MOPC 21) | | 0 | 1.19005698 | 2 |
| TMT | Q61554 | C1099 | Fibrillin-1 [Cleaved into: Asprosin] | | 0 | 1.179059728 | 3 |
| HPDP | Q80X90 | C1280 | Filamin-B (FLN-B) (ABP-280-like protein) (Actin-binding-like protein) (Beta-filamin) | | 0 | 1.169029965 | 2 |
| HPDP | D3YYU8 | C149 | Obscurin-like protein 1 | | 0 | 1.102544141 | 2 |
| HPDP | P53996 | C159 | Cellular nucleic acid-binding protein (CNBP) (Zinc finger protein 9) | | 0 | 1.087825704 | 2 |
| TMT | P56546 | C18 | C-terminal-binding protein 2 (CtBP2) | | 0 | 1.074827722 | 3 |
| HPDP | Q9JJ28 | C576 | Protein flightless-1 homolog | | 0 | 1.067369682 | 2 |
| HPDP | Q80X90 | C604 | Filamin-B (FLN-B) (ABP-280-like protein) (Actin-binding-like protein) (Beta-filamin) | | 0 | 1.066918314 | 2 |
| TMT | Q811Q4 | C430 | Disintegrin and metalloproteinase domain-containing protein 29 (ADAM 29) | | 0 | 1.066068555 | 3 |
| HPDP | A2ASS6 | C20340 | Titin | | 0 | 1.062863942 | 2 |

| | | | | | | | |
|---|---|---|---|---|---|---|---|
| HPDP | Q9DBF1 | C522 | Alpha-aminoadipic semialdehyde dehydrogenase (Alpha-AASA dehydrogenase) | | 0 | 1.062682493 | 2 |
| HPDP | P52480 | C49 | Pyruvate kinase PKM | | 0 | 1.060038804 | 2 |
| HPDP | Q9D1A2 | C300 | Cytosolic non-specific dipeptidase | | 0 | 1.044536941 | 2 |
| TMT | Q6R5P0 | C743 | Toll-like receptor 11 (Toll-like receptor 12) | | 0 | 1.020796529 | 4 |
| HPDP | Q8K4I3 | C563 | Rho guanine nucleotide exchange factor 6 (Alpha-PIX) (Rac/Cdc42 guanine nucleotide exchange factor | | 0 | 1.015633881 | 2 |
| HPDP | P31230 | C159 | Aminoacyl tRNA synthase complex-interacting multifunctional protein 1 (Multisynthase complex auxiliary component p43) [Cleaved into: Endothelial monocyte-activating polypeptide 2 (EMAP-2) (Endothelial monocyte-activating polypeptide II) (EMAP-II) (Small inducible cytokine subfamily E [illegible] | | 0 | 0.995152319 | 2 |
| HPDP | Q8VD04 | C104 | GRIP1-associated protein 1 (GRASP-1) (HCMV-interacting protein) [Cleaved into: GRASP-1 C-terminal chain (30kDa C-terminus form)] | | 0 | 0.963548687 | 3 |
| HPDP | Q8R146 | C309 | Acylamino-acid-releasing enzyme (AARE) | | 0 | 0.921391689 | 2 |
| TMT | P15920 | C315 | V-type proton ATPase 116 kDa subunit a isoform 2 (V-ATPase 116 kDa isoform a2) (Immune suppressor factor J6B7) (ISF) (Lysosomal H(+)-transporting ATPase V0 subunit a2) (ShIF) (Vacuolar proton translocating ATPase 116 kDa subunit a isoform 2) | | 0 | 0.912327311 | 2 |
| HPDP | P12399 | C103 | Protein CTLA-2-alpha (Cytotoxic T-lymphocyte-associated protein 2-alpha) | | 0 | 0.908725648 | 2 |
| HPDP | P41105 | C13 | 60S ribosomal protein L28 | | 0 | 0.866404857 | 2 |
| HPDP | Q8BGF6 | C98 | ELMO domain-containing protein 2 | | 0 | 0.863404858 | 2 |
| HPDP | Q9DCM0 | C34 | Persulfide dioxygenase ETHE1, mitochondrial | | 0 | 0.859102687 | 2 |
| HPDP | O54988 | C1136 | STE20-like serine/threonine-protein kinase (STE20-like kinase) (mSLK) | | 0 | 0.854916117 | 2 |
| HPDP | D3Z6Q9 | C425 | Bridging integrator 2 | | 0 | 0.828034499 | 2 |

| | | | | | | | |
|---|---|---|---|---|---|---|---|
| HPDP | P11983 | C357 | T-complex protein 1 subunit alpha (TCP-1-alpha) (CCT-alpha) (Tailless complex polypeptide 1A) (TCP-1-A) (Tailless complex polypeptide 1B) (TCP-1-B) | | 0 | 0.815170441 | 2 |
| HPDP | P62874 | C25 | Guanine nucleotide-binding protein G(I)/G(S)/G(T) subunit beta-1 (Transducin beta chain 1) | | 0 | 0.777801095 | 2 |
| HPDP | Q9JHW9 | C164 | Aldehyde dehydrogenase family 1 member A3 | | 0 | 0.775764306 | 2 |
| HPDP | Q64213 | C279 | Splicing factor 1 (CW17) (Mammalian branch point-binding protein) (BBP) (mBBP) (Transcription factor ZFM1) (mZFM) (Zinc finger gene in MEN1 locus) (Zinc finger protein 162) | | 0 | 0.727433087 | 2 |
| HPDP | Q80X90 | C450,C455 | Filamin-B (FLN-B) (ABP-280-like protein) (Actin-binding-like protein) (Beta-filamin) | | 0 | 0.723299502 | 2 |
| HPDP | P21981 | C370 | Protein-glutamine gamma-glutamyltransferase 2 | | 0 | 0.702284056 | 2 |
| HPDP | P62983 | C144,C155 | Ubiquitin-40S ribosomal protein S27a (Ubiquitin carboxyl extension protein 80) [Cleaved into: Ubiquitin; 40S ribosomal protein S27a] | | 0 | 0.697668035 | 2 |
| HPDP | Q8VDP3 | C82 | [F-actin]-monooxygenase MICAL1 | | 0 | 0.684887184 | 2 |
| HPDP | Q91W34 | C12 | RUS1 family protein C16orf58 homolog | | 0 | 0.643663222 | 2 |

Table S2: List of proteins nitrosylated in GSNOR$^{-/-}$ placentas as compared to B6 placentas and B6- and GSNOR$^{-/-}$ placentas treated with ascorbate. All SNOylated proteins were detected in at least 2 of 5 placentas/group. The number in the observation column is the number of placentas that showed expression of that particular SNOylated protein for that particular group.

| label | Uniprot Accession | SNO site | Protein name | Log2 fold change vs background in all 3 other groups | observations | Log2 fold change vs background GSNOR$^{-/-}$ | observations |
|---|---|---|---|---|---|---|---|
| TMT | Q8R418 | C306 | Endoribonuclease Dicer | | 0 | 13.0236246 | 5 |
| TMT | Q5FW85 | C8 | Extracellular matrix protein 2 (Tenonectin) | | 0 | 13.0094464 | 4 |
| TMT | Q8BV57 | C6 | Soluble scavenger receptor cysteine-rich domain-containing protein SSC5D (Scavenger receptor cysteine-rich domain-containing protein LOC284297 homolog) | | 0 | 12.7639881 | 5 |
| TMT | P70277 | C65 | Alpha-N-acetylgalactosaminide alpha-2,6-sialyltransferase 2 | | 0 | 12.6184686 | 5 |
| TMT | Q9JMG4 | C8,C14 | Sodium/potassium-transporting ATPase subunit beta-1-interacting protein 4 (Na(+)/K(+)-transporting ATPase subunit beta-1-interacting protein 4) (Protein FAM77A) | | 0 | 12.3930287 | 5 |
| TMT | A1Z198 | C330 | NACHT, LRR and PYD domains-containing protein 1b allele 2 | | 0 | 12.3651318 | 5 |
| TMT | Q9DB60 | C44,C47 | Prostamide/prostaglandin F synthase (Prostamide/PG F synthase) (Prostamide/PGF synthase) | | 0 | 12.3338002 | 5 |
| TMT | P57110 | C677 | A disintegrin and metalloproteinase with thrombospondin motifs 8 (ADAM-TS 8) (ADAM-TS8) (ADAMTS-8) | | 0 | 12.3028001 | 5 |
| TMT | Q571F5 | C271 | SPRY domain-containing SOCS box protein 3 (SSB-3) | | 0 | 12.2447841 | 5 |
| TMT | O09118 | C17 | Netrin-1 | | 0 | 12.2163032 | 5 |
| TMT | Q8C0W1 | C818 | Ankyrin repeat and MYND domain-containing protein 1 | | 0 | 12.2010848 | 5 |
| TMT | Q9DBP5 | C20 | UMP-CMP kinase | | 0 | 12.1445979 | 5 |
| TMT | O35963 | C48 | Ras-related protein Rab-33B | | 0 | 12.0873765 | 5 |
| TMT | A1L0T3 | C138 | Scavenger receptor cysteine-rich domain-containing group B protein (Four scavenger receptor cysteine-rich domains-containing protein) (S4D-SRCRB) | | 0 | 11.9017605 | 5 |

| | | | | | | | |
|---|---|---|---|---|---|---|---|
| TMT | Q8CGM1 | C1034 | Adhesion G protein-coupled receptor B2 (Brain-specific angiogenesis inhibitor 2) | | 0 | 11.8771311 | 5 |
| TMT | Q9WTN3 | C738,C753 | Sterol regulatory element-binding protein 1 (SREBP-1) (Sterol regulatory element-binding transcription factor 1) [Cleaved into: Processed sterol regulatory element-binding protein 1] | | 0 | 11.804481 | 5 |
| TMT | Q8CJ70 | C5,C18 | Interleukin-19 (IL-19) | | 0 | 11.6992587 | 5 |
| TMT | Q6DFV8 | C217 | von Willebrand factor D and EGF domain-containing protein | | 0 | 11.6741786 | 4 |
| TMT | Q8CGK5 | C81 | Interferon lambda receptor 1 (IFN-lambda R1) (Cytokine receptor class-II member 12) (Cytokine receptor family 2 member 12) (CRF2-12) (Interleukin-28 receptor subunit alpha) (IL-28 receptor subunit alpha) (IL-28R-alpha) (IL-28RA) | | 0 | 11.6126907 | 5 |
| TMT | Q4VAE3 | C31 | Transmembrane protein 65 | | 0 | 11.4110971 | 5 |
| TMT | Q9JLL3 | C25 | Tumor necrosis factor receptor superfamily member 19 (TRADE) (Toxicity and JNK inducer) | | 0 | 11.3577162 | 5 |
| TMT | Q8K400 | C293 | Syntaxin-binding protein 5 (Lethal(2) giant larvae protein homolog 3) (Tomosyn-1) | | 0 | 11.2111474 | 5 |
| TMT | Q9DC22 | C200,C211 | DDB1- and CUL4-associated factor 6 (IQ motif and WD repeat-containing protein 1) (Nuclear receptor interaction protein) (NRIP) | | 0 | 11.002862 | 5 |
| TMT | Q80VD1 | C52 | Protein FAM98B | | 0 | 3.25999526 | 5 |
| TMT | Q80XD8 | C9 | Proline-rich acidic protein 1 (Pregnancy-specific uterine protein) (Uterine-specific proline-rich acidic protein) | | 0 | 3.17282906 | 5 |
| TMT | Q9ESD6 | C54 | CKLF-like MARVEL transmembrane domain-containing protein 7 (Chemokine-like factor superfamily member 7) (LNV) | | 0 | 2.98489368 | 5 |
| TMT | Q9JIP3 | C12 | Interleukin-17 receptor B (IL-17 receptor B) (IL-17RB) (IL-17 receptor homolog 1) (IL-17ER) (IL-17Rh1) (IL17Rh1) (Interleukin-17B receptor) (IL-17B receptor) | | 0 | 2.92741104 | 5 |
| TMT | Q8BKK5 | C263 | Zinc finger protein 689 | | 0 | 2.9207658 | 5 |
| TMT | O08738 | C259 | Caspase-6 (CASP-6) | | 0 | 2.74260633 | 5 |
| TMT | O88282 | C381,C384 | B-cell CLL/lymphoma 6 member B protein (Bcl6-associated zinc finger protein) | | 0 | 2.72292467 | 5 |

| | | | | | | | |
|---|---|---|---|---|---|---|---|
| TMT | Q9Z0L3 | C371 | Otoconin-90 (Oc90) (Otoconin-95) (Oc95) | | 0 | 2.71298463 | 5 |
| TMT | E9PZZ1 | C653 | PR domain zinc finger protein 13 | | 0 | 2.54755478 | 4 |
| TMT | Q8BYA0 | C665 | Tubulin-specific chaperone D (Beta-tubulin cofactor D) (Tubulin-folding cofactor D) | | 0 | 2.48712789 | 5 |
| TMT | Q60676 | C221 | Serine/threonine-protein phosphatase 5 (PP5) | | 0 | 2.35208233 | 5 |
| TMT | Q5F2L2 | C13 | Alpha-(1,3)-fucosyltransferase 10 | | 0 | 2.31727696 | 5 |
| TMT | O09008 | C18 | Beta-1,3-N-acetylglucosaminyltransferase manic fringe | | 0 | 2.30522651 | 4 |
| HPDP | A2ASS6 | C21834 | Titin | | 0 | 2.24874462 | 2 |
| TMT | Q9QXW9 | C209 | Large neutral amino acids transporter small subunit 2 (L-type amino acid transporter 2) (mLAT2) (Solute carrier family 7 [illegible] | | 0 | 1.96367963 | 5 |
| TMT | Q66X22 | C891,C907,C908 | NACHT, LRR and PYD domains-containing protein 9B (NALP-delta) | | 0 | 1.93567695 | 4 |
| TMT | Q8BLY7 | C180 | Hermansky-Pudlak syndrome 6 protein homolog (Ruby-eye protein) (Ru) | | 0 | 1.87870793 | 5 |
| TMT | Q8K4G1 | C1393,C1403 | Latent-transforming growth factor beta-binding protein 4 (LTBP-4) | | 0 | 1.71923574 | 5 |
| TMT | Q7TPG7 | C107 | Protein FAM19A2 (Chemokine-like protein TAFA-2) | | 0 | 1.70638891 | 5 |
| TMT | P46097 | C91 | Synaptotagmin-2 (Inositol polyphosphate-binding protein) (IP4-binding protein) (IP4BP) (Synaptotagmin II) (SytII) | | 0 | 1.55432823 | 5 |
| TMT | P02798 | C33 | Metallothionein-2 (MT-2) (Metallothionein-II) (MT-II) | | 0 | 1.2666973 | 4 |
| HPDP | P01837 | C106 | Immunoglobulin kappa constant (Ig kappa chain C region MOPC 21) | | 0 | 1.19005698 | 2 |
| TMT | Q61554 | C1099 | Fibrillin-1 [Cleaved into: Asprosin] | | 0 | 1.17905973 | 3 |
| TMT | P56546 | C18 | C-terminal-binding protein 2 (CtBP2) | | 0 | 1.07482772 | 3 |
| TMT | Q811Q4 | C430 | Disintegrin and metalloproteinase domain-containing protein 29 (ADAM 29) | | 0 | 1.06606856 | 3 |
| TMT | Q6R5P0 | C743 | Toll-like receptor 11 (Toll-like receptor 12) | | 0 | 1.02079653 | 4 |

| | | | | | | | |
|---|---|---|---|---|---|---|---|
| TMT | P15920 | C315 | V-type proton ATPase 116 kDa subunit a isoform 2 (V-ATPase 116 kDa isoform a2) (Immune suppressor factor J6B7) (ISF) (Lysosomal H(+)-transporting ATPase V0 subunit a2) (ShIF) (Vacuolar proton translocating ATPase 116 kDa subunit a isoform 2) | | 0 | 0.91232731 | 2 |

| Table S3: List of peptides identified using mass spectrometry analysis. | | | | |
|---|---|---|---|---|
| Gene | Protein | SNO site | Peptide Modified Sequence | orginal label |
| 1433G_MOUSE | P61982 | C112 | NC[+57]SETQYESK | HPDP |
| A16A1_MOUSE | Q571I9 | C249 | VAFC[+57]GAVEEGR | HPDP |
| AAK1_MOUSE | Q3UHJ0 | C319 | EC[+57]PVPNVQNSPIPAK | HPDP |
| AATM_MOUSE | P05202 | C295 | VGAFTVVC[+329]K | IodoTMT6 |
| ABD12_MOUSE | Q99LR1 | C40 | C[+125]AASGSSSSGSAAAALDADC[+57]SLK | HPDP |
| ABI1_MOUSE | Q8CBW3 | C33 | VADYC[+57]ENNYIQATDK | HPDP |
| ABLM1_MOUSE | Q8K4G5 | C146 | NGDYLC[+57]TLDYQR | HPDP |
| ACINU_MOUSE | Q9JIX8 | C513 | SSLPEC[+57]STQK | HPDP |
| ACOC_MOUSE | P28271 | C392 | DFESC[+57]LGAK | HPDP |
| ACTG_MOUSE | P63260 | C17 | EEEIAALVIDNGSGMC[+329]K | IodoTMT6 |
| ACTN1_MOUSE | Q7TPR4 | C480 | IC[+57]DQWDNLGALTQK | HPDP |
| ACTN4_MOUSE | Q7TPR4 | C860 | ELPPDQAEYC[+329]IAR | IodoTMT6 |
| ADA29_MOUSE | Q811Q4 | C430 | EQC[+125]DC[+125]GSLRNC[+125]TNDLC[+329]CMSNCTLST( | IodoTMT6 |
| AEDO_MOUSE | Q6PDY2 | C225 | EASGSAC[+57]DLPR | HPDP |
| AGK_MOUSE | Q9ESW4 | C72 | ATVFLNPAAC[+57]K | HPDP |
| AGM1_MOUSE | Q9CYR6 | C200 | AFVDLTNQVSC[+57]SGDVK | HPDP |
| AGM1_MOUSE | Q9CYR6 | C348 | VPVYC[+57]TK | HPDP |
| AGRB2_MOUSE | Q8CGM1 | C1034 | FLC[+329]LGWGLPALVVAVSVGFTRTK | IodoTMT6 |
| AGRG4_MOUSE | B7ZCC9 | C2279,C2281 | GQGM[+16]DAIFHVPYSC[+329]AC[+329]WVIIKAKSSLESVEL | IodoTMT6 |
| AHDC1_MOUSE | Q6PAL7 | C788 | NC[+57]GFQGTEAR | HPDP |
| AIFM1_MOUSE | Q9Z0X1 | C440 | SNIWVAGDAAC[+57]FYDIK | HPDP |
| AIMP1_MOUSE | P31230 | C159 | IGC[+57]IVTAK | HPDP |
| AIMP2_MOUSE | Q8R010 | C168 | C[+57]FGEQAR | HPDP |
| AKA10_MOUSE | O88845 | C110 | SC[+57]LDYQTQETK | HPDP |
| AKA12_MOUSE | Q9WTQ5 | C1113 | ATTC[+57]QVIK | HPDP |
| AKP13_MOUSE | E9Q394 | C1644 | QQGFNYC[+57]TSAISSPLTK | HPDP |
| AKP13_MOUSE | E9Q394 | C417 | EGLPSC[+57]GNR | HPDP |
| AKP13_MOUSE | E9Q394 | C609 | VLGGQEPDTSIAGFC[+57]K | HPDP |
| AL1A3_MOUSE | Q9JHW9 | C164 | TIPTDDNVVC[+57]FTR | HPDP |
| AL3A2_MOUSE | P47739 | C229 | DC[+125]DLDVAC[+329]R | IodoTMT6 |

| | | | | |
|---|---|---|---|---|
| AL7A1_MOUSE | Q9DBF1 | C522 | STC[+57]TINYSTSLPLAQGIK | HPDP |
| ALBU_MOUSE | P07724 | C289 | YMC[+57]ENQATISSK | HPDP |
| ALBU_MOUSE | P07724 | C294 | YM[+16]C[+57]ENQATISSK | HPDP |
| ALBU_MOUSE | P07724 | C416 | TNC[+57]DLYEK | HPDP |
| ALBU_MOUSE | P07724 | C58 | C[+57]SYDEHAK | HPDP |
| ALBU_MOUSE | P07724 | C591 | DTC[+57]FSTEGPNLVTR | HPDP |
| ALBU_MOUSE | P07724 | C77, C86 | TC[+57]VADESAANC[+57]DK | HPDP |
| ALBU_MOUSE | P07724 | C289 | YMC[+329]ENQATISSK | IodoTMT6 |
| ALBU_MOUSE | P07724 | C591 | AADKDTC[+329]FSTEGPNLVTR | IodoTMT6 |
| ALBU_MOUSE | P07724 | C77,C86 | TC[+329]VADESAANC[+329]DK | IodoTMT6 |
| ALDH2_MOUSE | P47738 | C68 | TFPTVNPSTGEVIC[+57]QVAEGNK | HPDP |
| ALDOA_MOUSE | P05064 | C339 | ALANSLAC[+57]QGK | HPDP |
| ALDOA_MOUSE | P05064 | C339 | RALANSLAC[+57]QGK | HPDP |
| ALDOB_MOUSE | Q91Y97 | C158 | IADQC[+57]PSSLAIQENANALAR | HPDP |
| ALDOB_MOUSE | Q91Y97 | C269 | TVPAAVPGIC[+57]FLSGGMSEEDATLNLNAINR | HPDP |
| AMPL_MOUSE | Q9CPY7 | C445 | QVIDC[+57]QLADVNNLGK | HPDP |
| ANKL2_MOUSE | Q6P1H6 | C462 | TPEEVIC[+57]ER | HPDP |
| ANMY1_MOUSE | Q8C0W1 | C818 | ALYLSKRAELAPC[+329]HR | IodoTMT6 |
| ANR17_MOUSE | Q99NH0 | C2059 | NSPLDC[+57]GSASPNK | HPDP |
| ANX11_MOUSE | P97384 | C224 | GFGTDEQAIIDC[+57]LGSR | HPDP |
| ANXA4_MOUSE | P97429 | C141 | SLEEDIC[+329]SDTSFMFQR | IodoTMT6 |
| ANXA6_MOUSE | P14824 | C669 | ALLALC[+57]GGED | HPDP |
| AP1B1_MOUSE | O35643 | C863 | DC[+57]PLNTEAASNK | HPDP |
| AP1G1_MOUSE | P22892 | C400 | ADC[+57]ASGIFLAAEK | HPDP |
| AP2A2_MOUSE | P17427 | C902 | TTQIGC[+57]LLR | HPDP |
| APC1_MOUSE | P53995 | C988 | QAC[+57]EGNLPR | HPDP |
| APEH_MOUSE | Q8R146 | C309 | C[+125]ELLSDESLAVC[+57]SPR | HPDP |
| AQP1_MOUSE | Q02013 | C189 | DLGGSAPLAIGLSVALGHLLAIDYTGC[+329]GINPAR | IodoTMT6 |
| ARAF_MOUSE | P04627 | C595 | TQADELPAC[+57]LLSAAR | HPDP |
| ARFG1_MOUSE | Q9EPJ9 | C350 | SPSSDSWTC[+57]ADASTGR | HPDP |
| ARHG6_MOUSE | Q8K4I3 | C25 | TVC[+57]DPEEFLK | HPDP |
| ARHG6_MOUSE | Q8K4I3 | C563 | TSSSSC[+57]STHSSFSSTGQPR | HPDP |

| | | | | |
|---|---|---|---|---|
| ARHG7_MOUSE | Q9ES28 | C427 | NLSAQC[+57]QEVR | HPDP |
| ARHG7_MOUSE | Q9ES28 | C700 | VIEAYC[+57]TSAK | HPDP |
| ARHGC_MOUSE | Q8R4H2 | C1325 | TGTGDIATC[+57]DSPR | HPDP |
| ARP2_MOUSE | P61161 | C11 | VVVC[+57]DNGTGFVK | HPDP |
| ARP2_MOUSE | P61161 | C11 | KVVVC[+57]DNGTGFVK | HPDP |
| ARP2_MOUSE | P61161 | C221 | LC[+57]YVGYNIEQEQK | HPDP |
| ARP3_MOUSE | Q99JY9 | C408 | DYEEIGPSIC[+57]R | HPDP |
| ARP3_MOUSE | Q99JY9 | C408 | KDYEEIGPSIC[+57]R | HPDP |
| ARP3_MOUSE | Q99JY9 | C8, C12 | LPAC[+57]VVDC[+57]GTGYTK | HPDP |
| ARP3_MOUSE | Q99JY9 | C408 | KDYEEIGPSIC[+329]R | IodoTMT6 |
| ARP3_MOUSE | Q99JY9 | C408 | DYEEIGPSIC[+329]R | IodoTMT6 |
| ARRB1_MOUSE | Q8BWG8 | C150 | AFC[+57]AENLEEK | HPDP |
| AS3MT_MOUSE | Q91WU5 | C33 | TSADLQTNAC[+329]VTR | IodoTMT6 |
| ASAH1_MOUSE | Q9WV54 | C291 | SGEGC[+57]VITR | HPDP |
| AT131_MOUSE | Q9EPE9 | C333 | SPQENLVPC[+57]DVLLLR | HPDP |
| AT1A1_MOUSE | Q8VDN2 | C211 | IISANGC[+57]K | HPDP |
| AT2A2_MOUSE | O55143 | C377 | VEGDTC[+57]SLNEFSITGSTYAPIGEVQK | HPDP |
| AT2A2_MOUSE | O55143 | C669 | DAC[+57]LNAR | HPDP |
| ATS8_MOUSE | P57110 | C677 | GQC[+329]VKAGCDHVVNSPKK | IodoTMT6 |
| BAG3_MOUSE | Q9JLV1 | C185 | SQSPAASDC[+57]SSSSSSASLPSSGR | HPDP |
| BAG3_MOUSE | Q9JLV1 | C154 | QC[+329]GQMPATATTAAAQPPTAHGPER | IodoTMT6 |
| BCL6B_MOUSE | O88282 | C381,C384 | IHSGEKPYKC[+329]ETC[+329]GSRFVQVAHLR | IodoTMT6 |
| BDH_MOUSE | Q80XN0 | C209 | SPYC[+57]ITK | HPDP |
| BIN2_MOUSE | D3Z6Q9 | C425 | ASGSGSC[+57]NAPGSPEGSSQLC[+125]SPR | HPDP |
| BOLA1_MOUSE | Q9D8S9 | C126 | ENPQLDISPPC[+57]LGGSK | HPDP |
| BRF2_MOUSE | Q3UAW9 | C362 | RASPTPLLPPC[+329]MLKPPKR | IodoTMT6 |
| CA123_MOUSE | Q8BHG2 | C102 | TIVEFEC[+57]R | HPDP |
| CACL1_MOUSE | Q8R0X2 | C370 | AGDELAYNSPSAC[+57]ASSR | HPDP |
| CALR_MOUSE | P14211 | C105 | HEQNIDC[+57]GGGYVK | HPDP |
| CALR_MOUSE | P14211 | C137 | DMHGDSEYNIMFGPDIC[+329]GPGTKK | IodoTMT6 |
| CALR_MOUSE | P14211 | C137 | DMHGDSEYNIMFGPDIC[+329]GPGTK | IodoTMT6 |
| CAND2_MOUSE | Q6ZQ73 | C1138 | LATLC[+329]PAPVLQRVDRLIEPLR | IodoTMT6 |

| | | | | |
|---|---|---|---|---|
| CAP1_MOUSE | P40124 | C426 | NSLDC[+57]EIVSAK | HPDP |
| CAPZB_MOUSE | P47757 | C62 | DYLLC[+57]DYNR | HPDP |
| CARL1_MOUSE | Q6EDY6 | C712 | AC[+57]GGDAIQEDLK | HPDP |
| CASP6_MOUSE | O08738 | C259 | QVPC[+329]FASM[+16]LTKKLHFCPKPSK | IodoTMT6 |
| CATA_MOUSE | P24270 | C232 | LVNADGEAVYC[+57]K | HPDP |
| CATA_MOUSE | P24270 | C425 | SALEHSVQC[+57]AVDVK | HPDP |
| CATA_MOUSE | P24270 | C393 | DGPMC[+329]MHDNQGGAPNYYPNSFSAPEQQR | IodoTMT6 |
| CATB_MOUSE | P10605 | C211 | SC[+57]EAGYSPSYK | HPDP |
| CATB_MOUSE | P10605 | C211 | SC[+329]EAGYSPSYKEDK | IodoTMT6 |
| CATB_MOUSE | P10605 | C211 | SC[+329]EAGYSPSYK | IodoTMT6 |
| CATZ_MOUSE | Q9WUU7 | C156 | HGIPDETC[+329]NNYQAK | IodoTMT6 |
| CBX3_MOUSE | P23198 | C177 | LTWHSC[+57]PEDEAQ | HPDP |
| CDC23_MOUSE | Q8BGZ4 | C532 | LWDEASTC[+57]AQK | HPDP |
| CDC42_MOUSE | P60766 | C157 | YVEC[+57]SALTQK | HPDP |
| CEAM5_MOUSE | Q3UKK2 | C909 | C[+57]QLSIDPVWR | HPDP |
| CERU_MOUSE | Q61147 | C239 | TFC[+57]SEPEK | HPDP |
| CERU_MOUSE | Q61147 | C694 | GTFDVEC[+57]LTTDHYTGGM[+16]K | HPDP |
| CERU_MOUSE | Q61147 | C694 | GTFDVEC[+57]LTTDHYTGGMK | HPDP |
| CERU_MOUSE | Q61147 | C713 | YTVNQC[+57]QR | HPDP |
| CERU_MOUSE | Q61147 | C173 | ADDKVLPGQQYVYVLHANEPSPGEGDSNC[+329]VTR | IodoTMT6 |
| CERU_MOUSE | Q61147 | C173 | VLPGQQYVYVLHANEPSPGEGDSNC[+329]VTR | IodoTMT6 |
| CERU_MOUSE | Q61147 | C239 | TFC[+329]SEPEK | IodoTMT6 |
| CERU_MOUSE | Q61147 | C239 | TFC[+329]SEPEKVDKDNEDFQESNR | IodoTMT6 |
| CERU_MOUSE | Q61147 | C694 | GTFDVEC[+329]LTTDHYTGGMK | IodoTMT6 |
| CERU_MOUSE | Q61147 | C713 | YTVNQC[+329]QR | IodoTMT6 |
| CH082_MOUSE | Q8VE95 | C132 | LSYC[+57]GGGEALAIPFEPAR | HPDP |
| CHPT1_MOUSE | Q8C025 | C386 | TSC[+57]QQAPEQVYK | HPDP |
| CHRD1_MOUSE | Q9D1P4 | C211 | RKTSDFNTFLAQEGC[+329]TR | IodoTMT6 |
| CHRD1_MOUSE | Q9D1P4 | C211 | KTSDFNTFLAQEGC[+329]TR | IodoTMT6 |
| CHRD1_MOUSE | Q9D1P4 | C211 | TSDFNTFLAQEGC[+329]TR | IodoTMT6 |
| CK5P3_MOUSE | Q99LM2 | C136 | C[+57]QQLQQEYSR | HPDP |
| CKAP4_MOUSE | Q8BMK4 | C79 | SSAATANASSASC[+57]SR | HPDP |

| | | | | |
|---|---|---|---|---|
| CKAP4_MOUSE | Q8BMK4 | C79 | SSAATANASSASC[+329]SR | IodoTMT6 |
| CKLF7_MOUSE | Q9ESD6 | C54 | VAQMVTLLIAFIC[+329]VR | IodoTMT6 |
| CLCB_MOUSE | Q6IRU5 | C199 | VAQLC[+57]DFNPK | HPDP |
| CLH1_MOUSE | Q68FD5 | C1266 | EVC[+125]FAC[+57]VDGK | HPDP |
| CLH1_MOUSE | Q68FD5 | C1102 | C[+329]NEPAVWSQLAK | IodoTMT6 |
| CLIC1_MOUSE | Q9Z1Q5 | C223 | EEFASTC[+57]PDDEEIELAYEQVAR | HPDP |
| CMTR1_MOUSE | Q9DBC3 | C503 | SNESYC[+57]SLQIK | HPDP |
| CMTR1_MOUSE | Q9DBC3 | C534 | EC[+57]LQLWK | HPDP |
| CNBP_MOUSE | P53996 | C120 | C[+57]YSC[+125]GEFGHIQK | HPDP |
| CNBP_MOUSE | P53996 | C141 | C[+57]GETGHVAINC[+125]SK | HPDP |
| CNBP_MOUSE | P53996 | C141, C151 | C[+57]GETGHVAINC[+57]SK | HPDP |
| CNBP_MOUSE | P53996 | C157 | C[+125]GETGHVAINC[+57]SK | HPDP |
| CNBP_MOUSE | P53996 | C159 | TSEVNC[+57]YR | HPDP |
| CNBP_MOUSE | P53996 | C162 | C[+57]GESGHLAR | HPDP |
| CNDP2_MOUSE | Q9D1A2 | C300 | DVGAETLLHSC[+57]K | HPDP |
| CNDP2_MOUSE | Q9D1A2 | C300 | DVGAETLLHSC[+57]KK | HPDP |
| CNN2_MOUSE | Q08093 | C164 | AGQC[+57]VIGLQM[+16]GTNK | HPDP |
| CNN2_MOUSE | Q08093 | C164 | AGQC[+57]VIGLQMGTNK | HPDP |
| CNN2_MOUSE | Q08093 | C215 | C[+57]ASQVGM[+16]TAPGTR | HPDP |
| CO3_MOUSE | P01027 | C559 | DSC[+57]IGTLVVK | HPDP |
| CO4A1_MOUSE | P02463 | C1460 | HSQTTDDPLC[+329]PPGTK | IodoTMT6 |
| CO4A1_MOUSE | P02463 | C1493 | AHGQDLGTAGSC[+329]LR | IodoTMT6 |
| CO4A2_MOUSE | P08122 | C1532 | AHNQDLGLAGSC[+329]LAR | IodoTMT6 |
| COAC_MOUSE | Q8BZB2 | C7 | APC[+57]PAAVPSEER | HPDP |
| COF1_MOUSE | P18760 | C39 | AVLFC[+57]LSEDK | HPDP |
| COF1_MOUSE | P18760 | C39 | AVLFC[+57]LSEDKK | HPDP |
| COPB2_MOUSE | O55029 | C56 | TFEVC[+57]DLPVR | HPDP |
| COPB2_MOUSE | O55029 | C351 | DMGSC[+329]EIYPQTIQHNPNGR | IodoTMT6 |
| COR1A_MOUSE | O89053 | C24 | ADQC[+57]YEDVR | HPDP |
| COR1B_MOUSE | Q9WUM3 | C25 | NDQC[+57]YEDIR | HPDP |
| COR1C_MOUSE | Q9WUM4 | C190 | NGSLIC[+57]TASK | HPDP |
| COR1C_MOUSE | Q9WUM4 | C424 | KSELSC[+57]APK | HPDP |

| | | | | |
|---|---|---|---|---|
| CPNE1_MOUSE | Q8C166 | C52 | NC[+57]SSPEFSK | HPDP |
| CPNS1_MOUSE | O88456 | C145 | TDGFGIDTC[+57]R | HPDP |
| CPSF1_MOUSE | Q9EPU4 | C1042 | VYAVATSTNTPC[+57]TR | HPDP |
| CPSM_MOUSE | Q8C196 | C225 | VVAVDC[+57]GIK | HPDP |
| CPZIP_MOUSE | Q3UZA1 | C244 | NTC[+57]NSTEKPEELVR | HPDP |
| CRIP2_MOUSE | Q9DCT8 | C126 | ASSVTTFTGEPNMC[+329]PR | IodoTMT6 |
| CSDE1_MOUSE | Q91W50 | C129 | SPAAPGQSPTGSVC[+57]YER | HPDP |
| CSK22_MOUSE | O54833 | C336 | EQSQPC[+57]AENTVLSSGLTAAR | HPDP |
| CSRP1_MOUSE | P97315 | C122 | C[+57]SQAVYAAEK | HPDP |
| CSRP1_MOUSE | P97315 | C167 | GLESTTLADKDGEIYC[+329]K | IodoTMT6 |
| CSRP1_MOUSE | P97315 | C167 | C[+125]GKGLESTTLADKDGEIYC[+329]K | IodoTMT6 |
| CSRP1_MOUSE | P97315 | C58 | KNLDSTTVAVHGEEIYC[+329]K | IodoTMT6 |
| CSRP1_MOUSE | P97315 | C58 | NLDSTTVAVHGEEIYC[+329]K | IodoTMT6 |
| CSRP2_MOUSE | P97314 | C167 | EGEIYC[+57]K | HPDP |
| CSRP2_MOUSE | P97314 | C167 | SLESTTLTEKEGEIYC[+57]K | HPDP |
| CSRP2_MOUSE | P97314 | C167 | SLESTTLTEKEGEIYC[+329]K | IodoTMT6 |
| CSRP2_MOUSE | P97314 | C58 | NLDSTTVAIHDEEIYC[+329]K | IodoTMT6 |
| CTBP2_MOUSE | P56546 | C18 | LDRIC[+329]EGIRPQIM[+16]NGPLHPRPLVALLDGRDC[+125]T | IodoTMT6 |
| CTCF_MOUSE | Q61164 | C472 | YC[+329]DAVFHER | IodoTMT6 |
| CTL2_MOUSE | Q8BY89 | C401 | VVDDTAC[+57]PLLR | HPDP |
| CTL2A_MOUSE | P12399 | C103 | TNC[+57]YGNSLNR | HPDP |
| CTNA1_MOUSE | P26231 | C116 | SAAGEFADDPC[+57]SSVK | HPDP |
| CTNA1_MOUSE | P26231 | C116 | SAAGEFADDPC[+329]SSVK | IodoTMT6 |
| CTNA1_MOUSE | P26231 | C116 | SAAGEFADDPC[+329]SSVKR | IodoTMT6 |
| CUBN_MOUSE | Q9JLB4 | C1466 | IAQLC[+57]SR | HPDP |
| CUBN_MOUSE | Q9JLB4 | C1510 | AVPGGC[+57]GGIFQVSR | HPDP |
| CUBN_MOUSE | Q9JLB4 | C1927 | LIGTYC[+57]GTQR | HPDP |
| CUBN_MOUSE | Q9JLB4 | C2054 | LSQQLAVLC[+57]GR | HPDP |
| CUBN_MOUSE | Q9JLB4 | C817 | ADYQVAC[+57]GGELR | HPDP |
| CUL4B_MOUSE | A2A432 | C74 | SVC[+57]PGTSGFSSPNPSAASAAAQEVR | HPDP |
| CXA1_MOUSE | P23242 | C260 | SDPYHATTGPLSPSKDC[+329]GSPK | IodoTMT6 |
| CY24B_MOUSE | Q61093 | C126 | NLTFHKM[+16]VAWMIALHTAIHTIAHLFNVEWC[+329]VNAR | IodoTMT6 |

| | | | | |
|---|---|---|---|---|
| CYFP1_MOUSE | Q7TMB8 | C428 | DC[+57]PDNAEEYER | HPDP |
| D19L4_MOUSE | A2AJQ3 | C355 | VFEFYLLC[+329]TLPVTLNLIVK | IodoTMT6 |
| DAPK2_MOUSE | Q8VDF3 | C347 | NC[+57]ESDTEENIAR | HPDP |
| DCAF6_MOUSE | Q9DC22 | C200,C211 | DDILINC[+125]RRAATSVAIC[+329]PPVPYYLAVGC[+329]SDS | IodoTMT6 |
| DCSTP_MOUSE | Q7TNJ0 | C89 | RARC[+57]FILLAVLSC[+329]GLR | IodoTMT6 |
| DD19A_MOUSE | Q61655 | C392 | VLVTTNVC[+57]AR | HPDP |
| DDX1_MOUSE | Q91VR5 | C571 | FLIC[+57]TDVAAR | HPDP |
| DDX6_MOUSE | P54823 | C390 | NLVC[+57]TDLFTR | HPDP |
| DEN4C_MOUSE | A6H8H2 | C1083 | ILTAALTC[+57]PK | HPDP |
| DESM_MOUSE | P31001 | C332 | HQIQSYTC[+329]EIDALK | IodoTMT6 |
| DESM_MOUSE | P31001 | C332 | HQIQSYTC[+329]EIDALKGTNDSLMR | IodoTMT6 |
| DHE3_MOUSE | P26443 | C327 | C[+329]VGVGESDGSIWNPDGIDPK | IodoTMT6 |
| DHPR_MOUSE | Q8BVI4 | C82 | VDAILC[+57]VAGGWAGGNAK | HPDP |
| DHRS1_MOUSE | Q99L04 | C10 | GQVC[+57]VVTGASR | HPDP |
| DHSO_MOUSE | Q64442 | C106 | EVDEYC[+57]K | HPDP |
| DHX36_MOUSE | Q8VHK9 | C277 | AESC[+57]GNGNSTGYQIR | HPDP |
| DHX9_MOUSE | O70133 | C440 | AAEC[+57]NIVVTQPR | HPDP |
| DI3L2_MOUSE | Q8CI75 | C376 | DC[+57]IFTIDPSTAR | HPDP |
| DIAC_MOUSE | Q8R242 | C342 | GIGMWNANC[+329]LDYSDDALAR | IodoTMT6 |
| DIAP1_MOUSE | O08808 | C1210 | AGC[+57]AVTSLLASELTK | HPDP |
| DICER_MOUSE | Q8R418 | C306 | QILSDC[+329]RAVLVVLGPWC[+57]ADKVAGM[+16]M[+16]V | IodoTMT6 |
| DLGP4_MOUSE | B1AZP2 | C726 | DTDSDTQDANDSSC[+329]K | IodoTMT6 |
| DNJA2_MOUSE | Q9QYJ0 | C308 | VIEPGC[+57]VR | HPDP |
| DNJB6_MOUSE | O54946 | C243 | SLTINGVADENALAEEC[+329]QR | IodoTMT6 |
| DNPEP_MOUSE | Q9Z2W0 | C411 | NDSPC[+57]GTTIGPILASR | HPDP |
| DOPD_MOUSE | O35215 | C57 | STEPC[+329]AHLLVSSIGVVGTAEQNR | IodoTMT6 |
| DP13A_MOUSE | Q8K3H0 | C615 | IC[+57]DSVGLAK | HPDP |
| DPM3_MOUSE | Q9D1Q4 | C67 | VATFHDC[+57]EDAAR | HPDP |
| DPYL2_MOUSE | O08553 | C248 | SITIANQTNC[+57]PLYVTK | HPDP |
| DPYL2_MOUSE | O08553 | C248 | SITIANQTNC[+329]PLYVTK | IodoTMT6 |
| DPYL2_MOUSE | O08553 | C248 | SITIANQTNC[+329]PLYVTKVMSK | IodoTMT6 |
| DPYL2_MOUSE | O08553 | C439 | THNSALEYNIFEGM[+16]EC[+329]R | IodoTMT6 |

| | | | | |
|---|---|---|---|---|
| DPYL2_MOUSE | O08553 | C439 | THNSALEYNIFEGMEC[+329]R | IodoTMT6 |
| DSCR3_MOUSE | O35075 | C243 | DATEIQNIQIADGDIC[+57]R | HPDP |
| DYHC1_MOUSE | Q9JHU4 | C631 | VQYPQSQAC[+57]K | HPDP |
| DYN2_MOUSE | P39054 | C607 | QIELAC[+57]DSQEDVDSWK | HPDP |
| ECM2_MOUSE | Q5FW85 | C8 | LAVLFC[+329]FILLIVLQTDC[+125]ERGTR | IodoTMT6 |
| EDC3_MOUSE | Q8K2D3 | C137 | SQDVAISPQQQQC[+57]SK | HPDP |
| EF1D_MOUSE | P57776 | C217 | SSILLDVKPWDDETDMAQLETC[+329]VR | IodoTMT6 |
| EF2_MOUSE | P58252 | C369 | C[+57]ELLYEGPPDDEAAMGIK | HPDP |
| EF2_MOUSE | P58252 | C591 | ETVSEESNVLC[+57]LSK | HPDP |
| EF2_MOUSE | P58252 | C693 | EGALC[+57]EENMR | HPDP |
| EF2_MOUSE | P58252 | C693 | EGALC[+57]EENM[+16]R | HPDP |
| EF2_MOUSE | P58252 | C591 | ETVSEESNVLC[+329]LSK | IodoTMT6 |
| EF2_MOUSE | P58252 | C728 | C[+329]LYASVLTAQPR | IodoTMT6 |
| EFTU_MOUSE | Q8BFR5 | C290 | KGDEC[+57]ELLGHNK | HPDP |
| EIF3A_MOUSE | P23116 | C478 | HC[+57]DLQVR | HPDP |
| EIF3F_MOUSE | Q9DCH4 | C260 | TC[+57]FSPNR | HPDP |
| ELMD2_MOUSE | Q8BGF6 | C98 | TC[+57]LLQITGYK | HPDP |
| EMAL3_MOUSE | Q8VC03 | C421 | DSSC[+57]IVTSGK | HPDP |
| ENOA_MOUSE | P17182 | C337, C339 | SC[+57]NC[+57]LLLK | HPDP |
| ENOA_MOUSE | P17182 | C357 | VNQIGSVTESLQAC[+57]K | HPDP |
| EPN3_MOUSE | Q91W69 | C461 | SPSTVELDPFGDSSPSC[+329]K | IodoTMT6 |
| ERP44_MOUSE | Q9D1Q6 | C92 | VDC[+57]DQHSDIAQR | HPDP |
| ES8L2_MOUSE | Q99K30 | C546 | SGQAGYVPC[+57]NILAEAR | HPDP |
| ESTD_MOUSE | Q9R0P3 | C206 | AYDATC[+57]LVK | HPDP |
| ETHE1_MOUSE | Q9DCM0 | C170 | TDFQQGC[+57]AK | HPDP |
| ETHE1_MOUSE | Q9DCM0 | C34 | SC[+57]TYTYLLGDR | HPDP |
| EVI5_MOUSE | P97366 | C479 | LSEAESQC[+57]ALK | HPDP |
| EXOC8_MOUSE | Q6PGF7 | C419 | AC[+57]ELFLR | HPDP |
| F16P1_MOUSE | Q9QXD6 | C93 | SSYATC[+57]VLVSEENTNAIIIEPEK | HPDP |
| F19A2_MOUSE | Q7TPG7 | C107 | WWCHMQPC[+57]LEGEEC[+329]KVLPDR | IodoTMT6 |
| F19A2_MOUSE | Q7TPG7 | C96 | WWC[+329]HMQPC[+57]LEGEECKVLPDR | IodoTMT6 |
| FA98B_MOUSE | Q80VD1 | C52 | AAEGGLSSPEFSELC[+329]IWLGSQIK | IodoTMT6 |

| | | | | |
|---|---|---|---|---|
| FABP4_MOUSE | P04117 | C118 | LVVEC[+57]VMK | HPDP |
| FABP4_MOUSE | P04117 | C118 | LVVEC[+57]VM[+16]K | HPDP |
| FABP4_MOUSE | P04117 | C118 | DGDKLVVEC[+57]VMK | HPDP |
| FABP4_MOUSE | P04117 | C118 | DGDKLVVEC[+329]VMK | IodoTMT6 |
| FABP4_MOUSE | P04117 | C118 | LVVEC[+329]VMK | IodoTMT6 |
| FABPL_MOUSE | P12710 | C69 | NEFTLGEEC[+57]ELETMTGEK | HPDP |
| FABPL_MOUSE | P12710 | C69 | NEFTLGEEC[+57]ELETM[+16]TGEK | HPDP |
| FAK1_MOUSE | P34152 | C597 | NVLVSSNDC[+57]VK | HPDP |
| FARP1_MOUSE | F8VPU2 | C524 | QASPLISPLLNDQAC[+57]PR | HPDP |
| FAS_MOUSE | P19096 | C1181 | LLAAAC[+57]QLQLNGNLQLELGEALAQER | HPDP |
| FAS_MOUSE | P19096 | C223 | SFDDSGSGYC[+57]R | HPDP |
| FBN1_MOUSE | Q61554 | C1960 | C[+57]NEGYEVAPDGR | HPDP |
| FBN1_MOUSE | Q61554 | C1099 | GQCVNTPGDFEC[+57]KC[+329]DEGYESGFMM[+16]M[+16]K | IodoTMT6 |
| FBN1_MOUSE | Q61554 | C1960 | C[+329]NEGYEVAPDGR | IodoTMT6 |
| FERM2_MOUSE | Q8CIB5 | C426 | GC[+57]EVTPDVNISGQK | HPDP |
| FETA_MOUSE | P02772 | C144 | TAPASVPPFQFPEPAESC[+329]K | IodoTMT6 |
| FETUA_MOUSE | P29699 | C336 | VGQPGAAGPVSPMC[+329]PGR | IodoTMT6 |
| FGF5_MOUSE | P15656 | C200 | GC[+329]SPRVKPQHVSTHFLPR | IodoTMT6 |
| FHL1_MOUSE | P97447 | C255 | C[+57]SVNLANKR | HPDP |
| FHL2_MOUSE | O70433 | C71 | C[+57]GSSLVDKPFAAK | HPDP |
| FHL5_MOUSE | Q9WTX7 | C222,C225 | KC[+329]AAC[+329]TKPITGLRGAK | IodoTMT6 |
| FHOD1_MOUSE | Q6P9Q4 | C539 | AEPIQEPPTC[+57]VPK | HPDP |
| FINC_MOUSE | P11276 | C136 | ISC[+57]TIANR | HPDP |
| FINC_MOUSE | P11276 | C232 | C[+57]NDQDTR | HPDP |
| FKBP5_MOUSE | Q64378 | C215 | EEQC[+57]ILYLGPR | HPDP |
| FLII_MOUSE | Q9JJ28 | C1069 | TNGSALC[+57]TR | HPDP |
| FLII_MOUSE | Q9JJ28 | C560 | AC[+57]SAIHAVNLR | HPDP |
| FLII_MOUSE | Q9JJ28 | C576 | NYLGAEC[+57]R | HPDP |
| FLNA_MOUSE | Q8BTM8 | C1157 | AHVAPC[+57]FDASK | HPDP |
| FLNA_MOUSE | Q8BTM8 | C1453 | C[+57]SGPGLSPGMVR | HPDP |
| FLNA_MOUSE | Q8BTM8 | C2102 | VDINTEDLEDGTC[+57]R | HPDP |
| FLNA_MOUSE | Q8BTM8 | C2582 | SNFTVDC[+57]SK | HPDP |

| | | | | |
|---|---|---|---|---|
| FLNA_MOUSE | Q8BTM8 | C2601 | TPC[+57]EEILVK | HPDP |
| FLNA_MOUSE | Q8BTM8 | C8 | C[+57]GQSAAVASPGGSIDSR | HPDP |
| FLNA_MOUSE | Q8BTM8 | C1312 | VANPSGNLTDTYVQDC[+329]GDGTYK | IodoTMT6 |
| FLNA_MOUSE | Q8BTM8 | C2102 | VDINTEDLEDGTC[+329]R | IodoTMT6 |
| FLNA_MOUSE | Q8BTM8 | C2102 | DAGYGGLSLSIEGPSKVDINTEDLEDGTC[+329]R | IodoTMT6 |
| FLNA_MOUSE | Q8BTM8 | C2293 | DGSC[+329]GVAYVVQEPGDYEVSVK | IodoTMT6 |
| FLNA_MOUSE | Q8BTM8 | C2476 | MDC[+329]QEC[+125]PEGYR | IodoTMT6 |
| FLNA_MOUSE | Q8BTM8 | C8 | C[+329]GQSAAVASPGGSIDSR | IodoTMT6 |
| FLNB_MOUSE | Q80X90 | C1280 | AQITNPSGASTEC[+57]FVK | HPDP |
| FLNB_MOUSE | Q80X90 | C1326 | VAVTEGC[+57]QPSR | HPDP |
| FLNB_MOUSE | Q80X90 | C1434 | IAGPGLSSC[+57]VR | HPDP |
| FLNB_MOUSE | Q80X90 | C1868 | AEISC[+57]IDNK | HPDP |
| FLNB_MOUSE | Q80X90 | C1876 | DGTC[+57]TVTYLPTLPGDYSILVK | HPDP |
| FLNB_MOUSE | Q80X90 | C2057 | VDIQTEDLEDGTC[+57]K | HPDP |
| FLNB_MOUSE | Q80X90 | C2537 | SSFLVDC[+57]SK | HPDP |
| FLNB_MOUSE | Q80X90 | C450, C455 | SPFGVQIGEAC[+57]NPNAC[+57]R | HPDP |
| FLNB_MOUSE | Q80X90 | C604 | IEYDDQNDGSC[+57]DVK | HPDP |
| FLNB_MOUSE | Q80X90 | C660 | SGC[+57]TINNPAEFIVDPK | HPDP |
| FLNB_MOUSE | Q80X90 | C991 | VVPC[+57]LVAPVAGR | HPDP |
| FLNB_MOUSE | Q80X90 | C991 | KVVPC[+57]LVAPVAGR | HPDP |
| FLNB_MOUSE | Q80X90 | C2057 | VDIQTEDLEDGTC[+329]K | IodoTMT6 |
| FLNB_MOUSE | Q80X90 | C2057 | DAGYGGISLAVEGPSKVDIQTEDLEDGTC[+329]K | IodoTMT6 |
| FLNB_MOUSE | Q80X90 | C2248 | NGSC[+329]GVSYIAQEPGNYEVSIK | IodoTMT6 |
| FLNB_MOUSE | Q80X90 | C2333 | VHSPSGAVEEC[+329]HVSELEPDKYAVR | IodoTMT6 |
| FLNB_MOUSE | Q80X90 | C2431 | MDC[+329]QEIPEGYK | IodoTMT6 |
| FLNB_MOUSE | Q80X90 | C2537 | SSFLVDC[+329]SK | IodoTMT6 |
| FLNB_MOUSE | Q80X90 | C450 | SPFGVQIGEAC[+329]NPNAC[+125]R | IodoTMT6 |
| FLNB_MOUSE | Q80X90 | C455 | SPFGVQIGEAC[+125]NPNAC[+329]R | IodoTMT6 |
| FLNB_MOUSE | Q80X90 | C660 | SGC[+329]TINNPAEFIVDPK | IodoTMT6 |
| FLNB_MOUSE | Q80X90 | C769 | ANEPTHFTVDC[+329]TEAGEGDVSVGIK | IodoTMT6 |
| FLNB_MOUSE | Q80X90 | C769 | SGLKANEPTHFTVDC[+329]TEAGEGDVSVGIK | IodoTMT6 |
| FLNB_MOUSE | Q8VHX6 | C1104 | GAGTGGLGLTVEGPC[+329]EAK | IodoTMT6 |

| | | | | |
|---|---|---|---|---|
| FLNC_MOUSE | Q8VHX6 | C2661 | NSFTVDC[+57]SK | HPDP |
| FLNC_MOUSE | Q8VHX6 | C2680 | TPC[+329]EEVYVK | IodoTMT6 |
| FN3C1_MOUSE | Q6DFV6 | C1112 | TKPLPPEPPQLNC[+329]VVYGHQSLR | IodoTMT6 |
| FRPD1_MOUSE | A2AKB4 | C1360 | AYSC[+57]TTPLSR | HPDP |
| FSCN1_MOUSE | Q61553 | C481 | AC[+57]AETIDPASLWEY | HPDP |
| FSCN1_MOUSE | Q61553 | C89 | EVPDGDC[+57]R | HPDP |
| FUMH_MOUSE | P97807 | C431 | LLGDASVSFTDNC[+57]VVGIQANTER | HPDP |
| FUT10_MOUSE | Q5F2L2 | C13 | LLASC[+329]LCVTATVFLM[+16]VTLQVVVELGKFER | IodoTMT6 |
| FXL20_MOUSE | Q9CZV8 | C283 | C[+57]SQLTDVGFTTLAR | HPDP |
| FXR1_MOUSE | Q61584 | C77 | ANDQEPC[+57]GWWLAK | HPDP |
| FXR2_MOUSE | Q9WVR4 | C282 | IYGETPEAC[+57]R | HPDP |
| G3P_MOUSE | P16858 | C150 | IVSNASC[+57]TTNC[+125]LAPLAK | HPDP |
| G3P_MOUSE | P16858 | C150, C154 | IVSNASC[+57]TTNC[+57]LAPLAK | HPDP |
| G3P_MOUSE | P16858 | C160 | IVSNASC[+125]TTNC[+57]LAPLAK | HPDP |
| GAB1_MOUSE | Q9QYY0 | C406 | DASSQDC[+57]YDIPR | HPDP |
| GALK1_MOUSE | Q9R0N0 | C243 | QC[+57]EEVAQALGK | HPDP |
| GALT_MOUSE | Q03249 | C75 | HDPLNPLC[+57]PGATR | HPDP |
| GATM_MOUSE | Q9D964 | C87 | AENAC[+57]VPPFTVEVK | HPDP |
| GATM_MOUSE | Q9D964 | C64 | DC[+329]PVSSYNEWDPLEEVIVGR | IodoTMT6 |
| GATM_MOUSE | Q9D964 | C87 | AENAC[+329]VPPFTVEVK | IodoTMT6 |
| GBB1_MOUSE | P62874 | C25 | AC[+57]ADATLSQITNNIDPVGR | HPDP |
| GBB1_MOUSE | P62874 | C25 | KAC[+57]ADATLSQITNNIDPVGR | HPDP |
| GCNT1_MOUSE | Q09324 | C100 | DC[+57]ASFIR | HPDP |
| GFPT1_MOUSE | P47856 | C262 | VDSTTC[+57]LFPVEEK | HPDP |
| GFPT1_MOUSE | Q9Z2Z9 | C461 | ETDC[+329]GVHINAGPEIGVASTK | IodoTMT6 |
| GIMA4_MOUSE | Q99JY3 | C61 | VFNSGIC[+57]AK | HPDP |
| GLNA_MOUSE | P15105 | C183 | AC[+57]LYAGVK | HPDP |
| GLNA_MOUSE | P15105 | C49 | TLDC[+57]EPK | HPDP |
| GLOD4_MOUSE | Q9CPV4 | C45 | AAC[+57]NGPYDGK | HPDP |
| GMPR1_MOUSE | Q9DCZ1 | C186 | VGVGPGSVC[+57]TTR | HPDP |
| GON7_MOUSE | P0C8B4 | C21 | VSC[+57]EASGDADPLQSLSAGVVR | HPDP |
| GORS2_MOUSE | Q99JX3 | C434 | VSDC[+57]TPAVEKPVSDADASEPS | HPDP |

| | | | | |
|---|---|---|---|---|
| GP126_MOUSE | Q6F3F9 | C38 | C[+125]C[+57]PWRLKPSALLFLFVLC[+57]VTCVPLSVC[+125]( | IodoTMT6 |
| GP142_MOUSE | Q7TQN9 | C233 | LLKWAHCLIVYFIPC[+329]NVFLVTNSAIILR | IodoTMT6 |
| GPD1L_MOUSE | Q3ULJ0 | C216 | NIVAVGAGFC[+57]DGLR | HPDP |
| GPX41_MOUSE | O70325 | C195 | YGPMEEPQVIEKDLPC[+329]YL | IodoTMT6 |
| GRAP1_MOUSE | Q8VD04 | C104 | LC[+57]SQLEQLELENR | HPDP |
| GRB10_MOUSE | Q60760 | C173 | NQC[+329]PTDTVNPVAR | IodoTMT6 |
| GRM1B_MOUSE | Q80TI0 | C20 | STPAC[+57]SPILR | HPDP |
| GRP75_MOUSE | P38647 | C66 | GAVVGIDLGTTNSC[+57]VAVMEGK | HPDP |
| GRP78_MOUSE | P20029 | C42 | KEDVGTVVGIDLGTTYSC[+57]VGVFK | HPDP |
| GRP78_MOUSE | P20029 | C42 | EDVGTVVGIDLGTTYSC[+57]VGVFK | HPDP |
| GSHR_MOUSE | P47791 | C256 | NFDSLISSNC[+57]TEELENAGVEVLK | HPDP |
| GSHR_MOUSE | P47791 | C85 | AAVVESHKLGGTC[+125]VNVGC[+329]VPK | IodoTMT6 |
| GSHR_MOUSE | P47791 | C85 | LGGTC[+125]VNVGC[+329]VPK | IodoTMT6 |
| GSTK1_MOUSE | Q9DCM2 | C176 | LIENTDAAC[+57]K | HPDP |
| HA1K_MOUSE | P14428 | C317 | GGDYALAPGSQTSDLSLPDC[+329]K | IodoTMT6 |
| HDAC1_MOUSE | O09106 | C408 | ISIC[+57]SSDKR | HPDP |
| HEMO_MOUSE | Q91X72 | C458 | SLPQPQKVNSILGC[+57]SQ | HPDP |
| HEMO_MOUSE | Q91X72 | C458 | VNSILGC[+57]SQ | HPDP |
| HEMO_MOUSE | Q91X72 | C364 | ELGSPPGISLETIDAAFSC[+329]PGSSR | IodoTMT6 |
| HEMO_MOUSE | Q91X72 | C458 | SLPQPQKVNSILGC[+329]SQ | IodoTMT6 |
| HEMO_MOUSE | Q91X72 | C458 | VNSILGC[+329]SQ | IodoTMT6 |
| HIF1N_MOUSE | Q8BLR9 | C236 | RC[+125]ILFPPDQFEC[+57]LYPYPVHHPC[+329]DR | IodoTMT6 |
| HIF1N_MOUSE | Q8BLR9 | C236 | RC[+125]ILFPPDQFEC[+57]LYPYPVHHPC[+329]DR | IodoTMT6 |
| HMCS2_MOUSE | P54869 | C96, C106 | MGFC[+57]SVQEDINSLC[+57]LTVVQR | HPDP |
| HNRL1_MOUSE | Q8VDM6 | C533 | KAIVIC[+329]PTDEDLKDR | IodoTMT6 |
| HNRL1_MOUSE | Q8VDM6 | C533 | AIVIC[+329]PTDEDLKDR | IodoTMT6 |
| HNRPD_MOUSE | Q60668 | C126 | FGEVVDC[+57]TLK | HPDP |
| HNRPL_MOUSE | Q8R081 | C578 | LC[+57]FSTAQHAS | HPDP |
| HNRPQ_MOUSE | Q7TMK9 | C289 | GFC[+57]FLEYEDHK | HPDP |
| HNRPU_MOUSE | Q8VEK3 | C583 | KAVVVC[+57]PK | HPDP |
| HNRPU_MOUSE | Q8VEK3 | C583 | AVVVC[+57]PKDEDYK | HPDP |
| HPRT_MOUSE | P00493 | C106 | SYC[+57]NDQSTGDIK | HPDP |

| | | | | |
|---|---|---|---|---|
| HPRT_MOUSE | P00493 | C206 | DLNHVC[+57]VISETGK | HPDP |
| HPS6_MOUSE | Q8BLY7 | C180 | TLETSGEAGTKLGC[+329]THILLHHCPLFGLIASR | IodoTMT6 |
| HPS6_MOUSE | Q8BLY7 | C692 | LLLAEFAQHRRLDAHLPLLC[+329]R | IodoTMT6 |
| HSP72_MOUSE | P17156 | C606 | VC[+57]NPIISK | HPDP |
| HSP74_MOUSE | Q61316 | C167 | SVMDATQIAGLNC[+329]LR | IodoTMT6 |
| HSP7C_MOUSE | P63017 | C17 | GPAVGIDLGTTYSC[+57]VGVFQHGK | HPDP |
| HSP7C_MOUSE | P63017 | C603 | VC[+57]NPIITK | HPDP |
| HXK2_MOUSE | O08528 | C375 | IC[+57]QIVSTR | HPDP |
| I17RB_MOUSE | Q9JIP3 | C12 | MLLVLLILAASC[+329]RSALPR | IodoTMT6 |
| ICAL_MOUSE | P51125 | C408 | C[+329]GEDEDTVPAEYR | IodoTMT6 |
| IDHC_MOUSE | O88844 | C73 | C[+57]ATITPDEK | HPDP |
| IF2G_MOUSE | Q9Z0N1 | C105 | SC[+57]GSSTPDEFPTDIPGTK | HPDP |
| IF2G_MOUSE | Q9Z0N1 | C105 | SC[+329]GSSTPDEFPTDIPGTK | IodoTMT6 |
| IF4B_MOUSE | Q8BGD9 | C543 | VDVVGATQGQAGSC[+57]SR | HPDP |
| IF4B_MOUSE | Q8BGD9 | C543 | DGNKVDVVGATQGQAGSC[+57]SR | HPDP |
| IF5_MOUSE | P59325 | C122 | KQTIGNSC[+329]K | IodoTMT6 |
| IGHM_MOUSE | P01872 | C453 | STGKPTLYNVSLIM[+16]SDTGGTC[+329]Y | IodoTMT6 |
| IGHM_MOUSE | P01872 | C453 | STGKPTLYNVSLIMSDTGGTC[+329]Y | IodoTMT6 |
| IGHM_MOUSE | P01872 | C88 | SILEGSDEYLVC[+329]K | IodoTMT6 |
| IGKC_MOUSE | P01837 | C106 | SFNRNEC[+57] | HPDP |
| IGKC_MOUSE | P01837 | C86 | HNSYTC[+57]EATHK | HPDP |
| IL19_MOUSE | Q8CJ70 | C5,C18 | KTQC[+329]ASTWLLGM[+16]TLILC[+329]SVHIYSLRR | IodoTMT6 |
| INADL_MOUSE | Q63ZW7 | C1406 | ESESPDSAAC[+57]QIK | HPDP |
| INLR1_MOUSE | Q8CGK5 | C81 | TGWRPVEHCAGIKALVC[+329]PLMCLKK | IodoTMT6 |
| INSM1_MOUSE | Q63ZV0 | C482 | GAQERHLRLLHAAQVFPC[+329]K | IodoTMT6 |
| IRGQ_MOUSE | Q8VIM9 | C370 | AGIGDSGC[+57]TAAR | HPDP |
| KBL_MOUSE | O88986 | C26 | C[+57]ILDSELEGIR | HPDP |
| KBTBD_MOUSE | Q8C828 | C337 | GRLFVCLWRPADITAVVEYVVQMDKWLPVAELC[+329]R | IodoTMT6 |
| KCY_MOUSE | Q9DBP5 | C20 | KPLVVFVLGGPGAGKGTQC[+329]AR | IodoTMT6 |
| KINH_MOUSE | Q61768 | C632 | ELAAC[+57]QLR | HPDP |
| KNG1_MOUSE | O08677 | C339 | ESNTELAEDC[+57]EIK | HPDP |
| KNG1_MOUSE | O08677 | C125 | ENEFFIVTQTC[+329]K | IodoTMT6 |

| | | | | |
|---|---|---|---|---|
| KNG1_MOUSE | O08677 | C369 | C[+329]QALDMTEMAR | IodoTMT6 |
| KPCI_MOUSE | Q62074 | C190 | LVTIEC[+57]GR | HPDP |
| KPYM_MOUSE | P52480 | C423, C424 | C[+57]C[+57]SGAIIVLTK | HPDP |
| KPYM_MOUSE | P52480 | C49 | NTGIIC[+57]TIGPASR | HPDP |
| KPYM_MOUSE | P52480 | C423 | C[+329]C[+125]SGAIIVLTK | IodoTMT6 |
| KPYM_MOUSE | P52480 | C423 | C[+329]C[+125]SGAIIVLTKSGR | IodoTMT6 |
| KPYM_MOUSE | P52480 | C424 | C[+125]C[+329]SGAIIVLTK | IodoTMT6 |
| KPYM_MOUSE | P52480 | C49 | NTGIIC[+329]TIGPASR | IodoTMT6 |
| KPYR_MOUSE | P53657 | C360,C369 | VFLAQKMMIGRC[+329]NLAGKPVVC[+329]ATQMLESMITK | IodoTMT6 |
| KT3K_MOUSE | Q8K274 | C11 | ELGC[+57]SSVK | HPDP |
| LAC3_MOUSE | P01844 | C103 | SLSPAEC[+329]L | IodoTMT6 |
| LAMA5_MOUSE | Q61001 | C69 | ITASATC[+57]GEEAPTR | HPDP |
| LAMA5_MOUSE | Q61001 | C69 | ITASATC[+329]GEEAPTR | IodoTMT6 |
| LAMB1_MOUSE | P02469 | C643 | C[+329]GNTVPDDDNQVVSLSPGSR | IodoTMT6 |
| LAMC1_MOUSE | P02468 | C349 | SQEC[+329]YFDPELYR | IodoTMT6 |
| LAS1L_MOUSE | A2BE28 | C488 | VC[+57]SIYTQNGENGLAK | HPDP |
| LAT2_MOUSE | Q9QXW9 | C209 | LLALALIIIM[+16]GIVQIC[+329]K | IodoTMT6 |
| LAT4_MOUSE | Q8CGA3 | C295 | LC[+57]LSTVDLEVK | HPDP |
| LCAP_MOUSE | Q8C129 | C305 | SAFPC[+57]FDEPAFK | HPDP |
| LDHA_MOUSE | P06151 | C35 | ITVVGVGAVGMAC[+57]AISILMK | HPDP |
| LDHB_MOUSE | P16125 | C36 | ITVVGVGQVGMAC[+57]AISILGK | HPDP |
| LEG2_MOUSE | Q9CQW5 | C57 | FDESTIVC[+57]NTSEGGR | HPDP |
| LEG9_MOUSE | O08573 | C258 | C[+57]GGDIAFHLNPR | HPDP |
| LIPB2_MOUSE | O35711 | C398 | C[+57]VDGNQLSPVGEPK | HPDP |
| LKHA4_MOUSE | P24527 | C147 | AILPC[+57]QDTPSVK | HPDP |
| LKHA4_MOUSE | P24527 | C147 | AILPC[+329]QDTPSVK | IodoTMT6 |
| LKHA4_MOUSE | P24527 | C17 | PEVADTC[+125]SLASPASVC[+329]R | IodoTMT6 |
| LMCD1_MOUSE | Q8VEE1 | C246 | EVEYVC[+125]ELC[+329]K | IodoTMT6 |
| LMNA_MOUSE | P48678 | C522 | AQNTWGC[+57]GSSLR | HPDP |
| LMNA_MOUSE | P48678 | C590, C593 | TVLC[+57]GTC[+57]GQPADK | HPDP |
| LMNB2_MOUSE | P21619 | C190 | C[+57]QSLQEELAFSK | HPDP |
| LONP2_MOUSE | Q9DBN5 | C405 | IALGGVC[+57]DQSDIR | HPDP |

| | | | | |
|---|---|---|---|---|
| LR16A_MOUSE | Q6EDY6 | C1360 | C[+329]SDSGEEAEKEFIFV | IodoTMT6 |
| LRC47_MOUSE | Q505F5 | C544 | DGQC[+57]PLVVEQVR | HPDP |
| LRC59_MOUSE | Q922Q8 | C131 | VAGDC[+57]LDEK | HPDP |
| LRCH4_MOUSE | Q921G6 | C454 | AAGAGASAPSTQATC[+57]NGPPK | HPDP |
| LRP1B_MOUSE | Q9JI18 | C1540 | GPC[+329]SHLCLINHNRSAACAC[+125]PHLM[+16]KLSSDK | IodoTMT6 |
| LRP2_MOUSE | A2ARV4 | C2518 | AIVLDPC[+57]R | HPDP |
| LRP2_MOUSE | A2ARV4 | C2713 | C[+57]ISQDWK | HPDP |
| LRP2_MOUSE | A2ARV4 | C2830 | C[+329]QTTNIC[+57]VPR | IodoTMT6 |
| LSM7_MOUSE | Q9CQQ8 | C76 | QLGLVVC[+329]R | IodoTMT6 |
| LSM7_MOUSE | Q9CQQ8 | C76 | LTEDTRQLGLVVC[+329]R | IodoTMT6 |
| LTBP4_MOUSE | Q8K4G1 | C1393,C1403 | RC[+125]VSNESQSLDDNLGVC[+329]WQEVGPDLVC[+329]SR | IodoTMT6 |
| LY6C2_MOUSE | P0CW02 | C53 | ASDGFC[+329]IAQNIELIEDSQR | IodoTMT6 |
| MA7D1_MOUSE | A2AJI0 | C363 | THPSAAVPVC[+57]PR | HPDP |
| MAGI3_MOUSE | Q9EQJ9 | C1439 | AGC[+57]TPQSSSLVK | HPDP |
| MAOX_MOUSE | P06801 | C415 | AEC[+57]SAEQC[+125]YK | HPDP |
| MAOX_MOUSE | P06801 | C415, C420 | AEC[+57]SAEQC[+57]YK | HPDP |
| MAOX_MOUSE | P06801 | C415 | AEC[+329]SAEQC[+125]YK | IodoTMT6 |
| MAP1B_MOUSE | P14873 | C1913 | SPC[+329]DSGYSYETIEK | IodoTMT6 |
| MAP2_MOUSE | O08663 | C121 | VQTDPPSVPIC[+57]DLYPNGVFPK | HPDP |
| MAP4_MOUSE | P27546 | C636 | ETPGSQPSEPC[+57]SGVSR | HPDP |
| MAP4_MOUSE | P27546 | C636 | ETPGSQPSEPC[+329]SGVSR | IodoTMT6 |
| MARC2_MOUSE | Q922Q1 | C301 | LC[+57]DPSVK | HPDP |
| MCEE_MOUSE | Q9D1I5 | C168 | DC[+57]GGVLVELEQA | HPDP |
| MD1L1_MOUSE | Q9WTX8 | C233 | LC[+57]LQEQDAAVVK | HPDP |
| MDHC_MOUSE | P14152 | C137 | KSVKVIVVGNPANTNC[+329]LTASK | IodoTMT6 |
| MDHC_MOUSE | P14152 | C137 | SVKVIVVGNPANTNC[+329]LTASK | IodoTMT6 |
| MDHC_MOUSE | P14152 | C137 | VIVVGNPANTNC[+329]LTASK | IodoTMT6 |
| MDHC_MOUSE | P14152 | C154 | SAPSIPKENFSC[+329]LTR | IodoTMT6 |
| MDR1A_MOUSE | P21447 | C638 | LVMTQTAGNEIELGNEAC[+329]K | IodoTMT6 |
| MET7B_MOUSE | Q9DD20 | C96 | VTC[+57]VDPNPNFEK | HPDP |
| MFAP3_MOUSE | Q922T2 | C241 | SVPLPPLILNC[+329]RAFVEEMFEAVR | IodoTMT6 |
| MFN1_MOUSE | Q811U4 | C681 | LC[+57]QQVDVTQK | HPDP |

| | | | | |
|---|---|---|---|---|
| MFNG_MOUSE | O09008 | C18 | HC[+57]RLFRGM[+16]AGALFTLLC[+329]VGLLSLR | IodoTMT6 |
| MICA1_MOUSE | Q8VDP3 | C82 | ASQPVYQQGQAC[+57]TNTK | HPDP |
| MPRI_MOUSE | Q07113 | C1910 | SYDEC[+329]VLEGR | IodoTMT6 |
| MPRIP_MOUSE | P97434 | C723 | EGYVLQATC[+57]ER | HPDP |
| MT1_MOUSE | P02802 | C44 | C[+329]AQGC[+125]VC[+125]K | IodoTMT6 |
| MT1_MOUSE | P02802 | C44 | C[+329]AQGC[+57]VC[+57]K | IodoTMT6 |
| MT1_MOUSE | P02802 | C50 | C[+125]AQGC[+125]VC[+329]K | IodoTMT6 |
| MT1_MOUSE | P02802 | C50 | C[+125]AQGC[+57]VC[+329]K | IodoTMT6 |
| MT2_MOUSE | P02798 | C44, C48, C50 | C[+57]SQGC[+57]IC[+57]K | HPDP |
| MT2_MOUSE | P02798 | C54, C56 | C[+125]SQGC[+57]IC[+57]K | HPDP |
| MT2_MOUSE | P02798 | C33 | SC[+329]C[+57]SC[+125]C[+57]PVGC[+57]AK | IodoTMT6 |
| MT2_MOUSE | P02798 | C33,C41 | SC[+329]C[+57]SC[+125]C[+57]PVGC[+329]AK | IodoTMT6 |
| MT2_MOUSE | P02798 | C33,C41 | SC[+329]C[+125]SC[+125]C[+57]PVGC[+329]AK | IodoTMT6 |
| MT2_MOUSE | P02798 | C34,C41 | SC[+125]C[+329]SC[+125]C[+125]PVGC[+329]AK | IodoTMT6 |
| MT2_MOUSE | P02798 | C36,C41 | SC[+57]C[+125]SC[+329]C[+57]PVGC[+329]AK | IodoTMT6 |
| MT2_MOUSE | P02798 | C37,C41 | SC[+57]C[+57]SC[+125]C[+329]PVGC[+329]AK | IodoTMT6 |
| MT2_MOUSE | P02798 | C41 | SC[+57]C[+57]SC[+125]C[+57]PVGC[+329]AK | IodoTMT6 |
| MT2_MOUSE | P02798 | C41 | SC[+57]C[+125]SC[+57]C[+57]PVGC[+329]AK | IodoTMT6 |
| MT2_MOUSE | P02798 | C41 | SC[+125]C[+57]SC[+125]C[+57]PVGC[+329]AK | IodoTMT6 |
| MT2_MOUSE | P02798 | C41 | SC[+57]C[+57]SC[+125]C[+125]PVGC[+329]AK | IodoTMT6 |
| MT2_MOUSE | P02798 | C41 | SC[+125]C[+125]SC[+57]C[+57]PVGC[+329]AK | IodoTMT6 |
| MT2_MOUSE | P02798 | C41 | SC[+125]C[+125]SC[+125]C[+57]PVGC[+329]AK | IodoTMT6 |
| MT2_MOUSE | P02798 | C44 | C[+329]SQGC[+125]IC[+125]K | IodoTMT6 |
| MT2_MOUSE | P02798 | C44 | C[+329]SQGC[+57]IC[+125]K | IodoTMT6 |
| MT2_MOUSE | P02798 | C50 | C[+125]SQGC[+125]IC[+329]K | IodoTMT6 |
| MT2_MOUSE | P02798 | C50 | C[+125]SQGC[+57]IC[+329]K | IodoTMT6 |
| MTCH2_MOUSE | Q791V5 | C79 | LC[+57]SGVLGTVVHGK | HPDP |
| MTMR2_MOUSE | Q9Z2D1 | C95 | DVTYIC[+57]PFTGAVR | HPDP |
| MTMR7_MOUSE | Q9Z2C9 | C158 | VC[+57]DSYPTELYVPR | HPDP |
| MTMR7_MOUSE | Q9Z2C9 | C68 | QATTATGC[+57]PLLIR | HPDP |
| MTMR9_MOUSE | Q9Z2D0 | C392 | C[+125]AQSAYC[+57]SSK | HPDP |
| MTMRA_MOUSE | Q7TPM9 | C703 | SGPLEAC[+57]YAELDQSR | HPDP |

| | | | | |
|---|---|---|---|---|
| MTNB_MOUSE | Q9WVQ5 | C187 | MAHAMNEYPDSC[+329]AVLVRR | IodoTMT6 |
| MTNB_MOUSE | Q9WVQ5 | C187 | MAHAMNEYPDSC[+329]AVLVR | IodoTMT6 |
| MTR1L_MOUSE | O88495 | C363 | AC[+57]VAVEGTPR | HPDP |
| MTUS1_MOUSE | Q5HZI1 | C823 | SLC[+57]IQTQTAPDVLSSER | HPDP |
| MYG_MOUSE | P04247 | C67 | HGC[+57]TVLTALGTILK | HPDP |
| MYH9_MOUSE | Q8VDD5 | C740 | QAC[+57]VLMIK | HPDP |
| MYH9_MOUSE | Q8VDD5 | C816 | NC[+57]AAYLR | HPDP |
| MYH9_MOUSE | Q8VDD5 | C896 | LQLQEQLQAETELC[+57]AEAEELR | HPDP |
| MYH9_MOUSE | Q8VDD5 | C988 | KLEEDQIIM[+16]EDQNC[+329]K | IodoTMT6 |
| MYH9_MOUSE | Q8VDD5 | C988 | LKKLEEDQIIMEDQNC[+329]K | IodoTMT6 |
| MYH9_MOUSE | Q8VDD5 | C988 | KLEEDQIIMEDQNC[+329]K | IodoTMT6 |
| MYH9_MOUSE | Q8VDD5 | C988 | LEEDQIIM[+16]EDQNC[+329]K | IodoTMT6 |
| MYH9_MOUSE | Q8VDD5 | C988 | LEEDQIIMEDQNC[+329]K | IodoTMT6 |
| MYL3_MOUSE | P09542 | C191 | LMAGQEDSNGC[+57]INYEAFVK | HPDP |
| MYL6_MOUSE | Q60605 | C32 | ILYSQC[+57]GDVM[+16]R | HPDP |
| MYL6_MOUSE | Q60605 | C32 | ILYSQC[+57]GDVMR | HPDP |
| MYO1E_MOUSE | E9Q634 | C960 | AAPAPPGC[+57]HQNGVIR | HPDP |
| MYOF_MOUSE | Q69ZN7 | C1540 | ELPDSVPQEC[+57]TVR | HPDP |
| MYOF_MOUSE | Q69ZN7 | C409 | VC[+57]TNIIER | HPDP |
| MYOF_MOUSE | Q69ZN7 | C409 | KVC[+57]TNIIER | HPDP |
| MYOM1_MOUSE | Q62234 | C656 | C[+57]DVGAENWQR | HPDP |
| MYOTI_MOUSE | Q9JIF9 | C321 | ASDAGPYAC[+57]VAR | HPDP |
| MYPC3_MOUSE | O70468 | C1204 | SIIAGYNAILC[+125]C[+57]AVR | HPDP |
| NAL9B_MOUSE | Q66X22 | C891,C907,C90 | QLC[+57]EALSHPNC[+329]NLEC[+57]LGLDLCEFTSDC[+329]( | IodoTMT6 |
| NARFL_MOUSE | Q7TMW6 | C270 | DVDC[+57]VLTTGEVFR | HPDP |
| NDKA_MOUSE | P15532 | C145 | SC[+329]AQNWIYE | IodoTMT6 |
| NDKB_MOUSE | Q01768 | C145 | SC[+329]AHDWVYE | IodoTMT6 |
| NDUV1_MOUSE | Q91YT0 | C125 | YLVVNADEGEPGTC[+329]K | IodoTMT6 |
| NET1_MOUSE | O09118 | C17 | M[+16]MRAVWEALAALAAVAC[+329]LVGAVR | IodoTMT6 |
| NEXN_MOUSE | Q7TPW1 | C585 | GETYC[+125]LYLPETFPEDGGEYMC[+329]K | IodoTMT6 |
| NHRF1_MOUSE | P70441 | C201 | IVEVNGVC[+57]M[+16]EGK | HPDP |
| NHRF1_MOUSE | P70441 | C201 | IVEVNGVC[+57]MEGK | HPDP |

| | | | | |
|---|---|---|---|---|
| NID1_MOUSE | P10493 | C1232 | C[+329]PDNTLGVDC[+57]IER | IodoTMT6 |
| NIPA_MOUSE | Q80YV2 | C405 | LC[+57]SSSSSDTSPR | HPDP |
| NIT1_MOUSE | Q8VDK1 | C247, C255 | AIESQC[+57]YVIAAAQC[+57]GR | HPDP |
| NKAI4_MOUSE | Q9JMG4 | C8,C14 | C[+329]TLLALC[+329]ALQLVTALER | IodoTMT6 |
| NL1B2_MOUSE | A1Z198 | C330 | QIFGIKALMMVESNPVLLTLCEVPWVCWLVC[+329]NC[+125]I | IodoTMT6 |
| NMDZ1_MOUSE | P35438 | C459 | KVIC[+125]TGPNDTSPGSPRHTVPQC[+125]C[+125]YGFC[+32 | IodoTMT6 |
| NOP58_MOUSE | Q6DFW4 | C439 | TYDPSGDSTLPTC[+57]SK | HPDP |
| NPAS4_MOUSE | Q8BGD7 | C149 | QQLTM[+16]PSALDADRLFRC[+329]R | IodoTMT6 |
| NRDC_MOUSE | Q8BHG1 | C685 | AFDC[+57]PETEYPAK | HPDP |
| NSF_MOUSE | P46460 | C11 | C[+57]PTDELSLSNC[+125]AVVNEK | HPDP |
| NSF_MOUSE | P46460 | C11, C21 | C[+57]PTDELSLSNC[+57]AVVNEK | HPDP |
| NTKL_MOUSE | Q9EQC5 | C241 | SLVTHYC[+329]ELVGANPKVRPNPARFLQNCR | IodoTMT6 |
| NUMA1_MOUSE | E9Q7G0 | C728 | AADALKEQQC[+57]R | HPDP |
| NUMB_MOUSE | Q9QZS3 | C176 | EC[+57]GVTATFDASR | HPDP |
| OBSCN_MOUSE | A2AAJ9 | C3100 | GTLTLQC[+57]EVSDPEAR | HPDP |
| OBSCN_MOUSE | A2AAJ9 | C3864 | SLTIADAGEYLC[+57]TC[+125]GQEK | HPDP |
| OBSCN_MOUSE | A2AAJ9 | C4816 | DLTVEDTGEYSC[+57]TC[+125]GQER | HPDP |
| OBSL1_MOUSE | D3YYU8 | C149 | GEEVVLTC[+57]QVGGLPEPK | HPDP |
| OC90_MOUSE | Q9Z0L3 | C371 | QVGC[+329]LHGRRSQSSVVCEDHMAK | IodoTMT6 |
| OSBL1_MOUSE | Q91XL9 | C520 | DC[+57]GGGDALSNGIK | HPDP |
| OSBP1_MOUSE | Q3B7Z2 | C222 | VEDLSTC[+57]NDLIAK | HPDP |
| OXSM_MOUSE | Q9D404 | C86 | NIPC[+57]SVAAYVPR | HPDP |
| P3H1_MOUSE | Q3V1T4 | C648 | TVTAEVQPQC[+57]GR | HPDP |
| PA2G4_MOUSE | P50580 | C49 | SLVEASSSGVSVLSLC[+57]EK | HPDP |
| PACN2_MOUSE | Q9WVE8 | C465 | IEDEDEQGWC[+329]K | IodoTMT6 |
| PACN2_MOUSE | Q9WVE8 | C465 | AGDELTKIEDEDEQGWC[+329]K | IodoTMT6 |
| PAFA_MOUSE | Q60963 | C290 | C[+329]GVALDPWMYPVNEELYSR | IodoTMT6 |
| PAFA_MOUSE | Q60963 | C290 | C[+329]GVALDPWM[+16]YPVNEELYSR | IodoTMT6 |
| PAFA_MOUSE | Q60963 | C290 | C[+329]GVALDPWMYPVNEELYSR | IodoTMT6 |
| PARK7_MOUSE | Q99LX0 | C53 | DVMIC[+57]PDTSLEDAK | HPDP |
| PARK7_MOUSE | Q99LX0 | C58 | DVM[+16]IC[+57]PDTSLEDAK | HPDP |
| PARK7_MOUSE | Q99LX0 | C53 | DVMIC[+329]PDTSLEDAK | IodoTMT6 |

| | | | | |
|---|---|---|---|---|
| PAXI_MOUSE | Q8VI36 | C108 | NSSASNTQDGVGSLC[+329]SR | IodoTMT6 |
| PCBP1_MOUSE | P60335 | C109 | LVVPATQC[+57]GSLIGK | HPDP |
| PCCB_MOUSE | Q99MN9 | C271 | AFDNDVDALC[+57]NLR | HPDP |
| PDC6I_MOUSE | Q9WU78 | C40 | FIQQTYPSGGEEQAQYC[+57]R | HPDP |
| PDIA5_MOUSE | Q921X9 | C449, C454 | IAC[+57]AAVDC[+57]VK | HPDP |
| PDIA5_MOUSE | Q921X9 | C449, C454 | KIAC[+57]AAVDC[+57]VK | HPDP |
| PDIA5_MOUSE | Q921X9 | C463 | DKNQDLC[+57]QQEAVK | HPDP |
| PDLI1_MOUSE | O70400 | C73 | GC[+57]ADNM[+16]TLTVSR | HPDP |
| PDLI1_MOUSE | O70400 | C73 | GC[+57]ADNMTLTVSR | HPDP |
| PDLI5_MOUSE | Q8CI51 | C73 | AC[+57]TGSLNMTLQR | HPDP |
| PDLI5_MOUSE | Q8CI51 | C73 | AC[+57]TGSLNM[+16]TLQR | HPDP |
| PEA15_MOUSE | Q62048 | C27 | SAC[+57]KEDIPSEK | HPDP |
| PEG10_MOUSE | Q7TN75 | C521 | SIVFNSDYC[+57]R | HPDP |
| PEG10_MOUSE | Q7TN75 | C521 | SIVFNSDYC[+329]R | IodoTMT6 |
| PEG3_MOUSE | Q3URU2 | C383 | EC[+57]GETFSR | HPDP |
| PELP1_MOUSE | Q9DBD5 | C202 | AC[+57]VTYFPR | HPDP |
| PELP1_MOUSE | Q9DBD5 | C523 | NANSDVC[+57]AAALR | HPDP |
| PEPD_MOUSE | Q11136 | C183 | FNVNNTILHPEIVEC[+329]R | IodoTMT6 |
| PEPD_MOUSE | Q11136 | C183 | EASFEGISKFNVNNTILHPEIVEC[+329]R | IodoTMT6 |
| PEPD_MOUSE | Q11136 | C482 | TVEEIEAC[+125]MAGC[+329]DK | IodoTMT6 |
| PEPD_MOUSE | Q11136 | C58 | YC[+329]TDTSIIFR | IodoTMT6 |
| PEPL1_MOUSE | Q6NSR8 | C357 | LVLADGVSYAC[+57]K | HPDP |
| PFD3_MOUSE | P61759 | C8 | DGC[+57]GLETAAGNGR | HPDP |
| PFKAP_MOUSE | Q9WUA3 | C410 | SNC[+57]NVAVINVGAPAAGMNAAVR | HPDP |
| PFKAP_MOUSE | Q9WUA3 | C410 | SNC[+329]NVAVINVGAPAAGMNAAVR | IodoTMT6 |
| PGBM_MOUSE | Q05793 | C1628 | TC[+57]ESLGAGGYR | HPDP |
| PGBM_MOUSE | Q05793 | C2456 | DITLEC[+57]ISSGEPR | HPDP |
| PGBM_MOUSE | Q05793 | C1530 | ALEVEEC[+329]R | IodoTMT6 |
| PGBM_MOUSE | Q05793 | C2456 | DITLEC[+329]ISSGEPR | IodoTMT6 |
| PGBM_MOUSE | Q05793 | C479 | EADQGAYTC[+329]EAMNSR | IodoTMT6 |
| PGBM_MOUSE | Q05793 | C892 | GSLGTSGETC[+329]R | IodoTMT6 |
| PGFS_MOUSE | Q9DB60 | C44,C47 | AC[+57]VVAGLRRFGC[+329]MVC[+329]R | IodoTMT6 |

| | | | | |
|---|---|---|---|---|
| PGS1_MOUSE | P28653 | C77 | VVQC[+57]SDLGLK | HPDP |
| PHAG1_MOUSE | Q3U1F9 | C423 | ESDYESIGDLQQC[+57]R | HPDP |
| PHAG1_MOUSE | Q3U1F9 | C423 | ESDYESIGDLQQC[+329]R | IodoTMT6 |
| PIPNB_MOUSE | P53811 | C187 | ELANTPDC[+57]PR | HPDP |
| PKHA7_MOUSE | Q3UIL6 | C969 | DREQGQC[+57]VNGDLK | HPDP |
| PKHM1_MOUSE | Q7TSI1 | C464 | SAAGLC[+57]TSPVQDTPESR | HPDP |
| PKP3_MOUSE | Q9QY23 | C129 | SAVDLTC[+57]SR | HPDP |
| PLEC_MOUSE | Q9QXS1 | C1386 | QEQIQAVPIANC[+57]QAAR | HPDP |
| PLEC_MOUSE | Q9QXS1 | C4267 | C[+125]ITDPQTGLC[+57]LLPLKEK | HPDP |
| PLEC_MOUSE | Q9QXS1 | C4267 | C[+125]ITDPQTGLC[+57]LLPLK | HPDP |
| PLRG1_MOUSE | Q922V4 | C208 | C[+57]IAVEPGNQWFVTGSADR | HPDP |
| PLRG1_MOUSE | Q922V4 | C263 | SPYLFSC[+329]GEDK | IodoTMT6 |
| PLRG1_MOUSE | Q922V4 | C337 | C[+329]QAAEPQIITGSHDTTIRLWDLVAGKTR | IodoTMT6 |
| PLRG1_MOUSE | Q922V4 | C337 | C[+329]QAAEPQIITGSHDTTIR | IodoTMT6 |
| PMGE_MOUSE | P15327 | C145 | VC[+57]DVPLDQLPR | HPDP |
| PP2AB_MOUSE | P63330 | C269 | C[+329]GNQAAIMELDDTLK | IodoTMT6 |
| PP4P1_MOUSE | Q3TWL2 | C94 | VC[+57]QSPINVEGK | HPDP |
| PP4R1_MOUSE | Q8K2V1 | C385 | LESLEGC[+57]AAK | HPDP |
| PP6R3_MOUSE | Q922D4 | C815 | C[+329]TAPLTPSSSPEQR | IodoTMT6 |
| PPBT_MOUSE | P09242 | C119 | TYNTNAQVPDSAGTATAYLC[+329]GVK | IodoTMT6 |
| PPIA_MOUSE | P17742 | C161 | KITISDC[+57]GQL | HPDP |
| PPIA_MOUSE | P17742 | C161 | ITISDC[+57]GQL | HPDP |
| PPIA_MOUSE | P17742 | C161 | TSKKITISDC[+57]GQL | HPDP |
| PPIA_MOUSE | P17742 | C67 | IIPGFM[+16]C[+57]QGGDFTR | HPDP |
| PPIA_MOUSE | P17742 | C161 | KITISDC[+329]GQL | IodoTMT6 |
| PPIA_MOUSE | P17742 | C62 | IIPGFMC[+329]QGGDFTR | IodoTMT6 |
| PPIG_MOUSE | A2AR02 | C174 | ILSC[+57]GELIPK | HPDP |
| PPIG_MOUSE | A2AR02 | C308 | EC[+57]NPPNSQPASYQR | HPDP |
| PPME1_MOUSE | Q8BVQ5 | C238 | QC[+57]EGITSPEGSK | HPDP |
| PPP5_MOUSE | Q60676 | C404 | GVSC[+57]QFGPDVTK | HPDP |
| PPP5_MOUSE | Q60676 | C221 | EVLC[+329]KLSTLVETTLK | IodoTMT6 |
| PPR3F_MOUSE | Q9JIG4 | C419 | ILPATC[+57]GLGGPPR | HPDP |

| | | | | |
|---|---|---|---|---|
| PR8A9_MOUSE | Q9CQ58 | C101 | AGTYC[+57]HSTLSNPPDR | HPDP |
| PRAP1_MOUSE | Q80XD8 | C9 | RFLLATC[+329]LVAALLWEAGAAPAHQVPVK | IodoTMT6 |
| PRD13_MOUSE | E9PZZ1 | C653 | THTGYKPLKC[+125]KVC[+329]LRPFGDPSNLNK | IodoTMT6 |
| PRDX4_MOUSE | O08807 | C54 | ENEC[+329]HFYAGGQVYPGEASR | IodoTMT6 |
| PRDX5_MOUSE | P99029 | C200 | ALNVEPDGTGLTC[+57]SLAPNILSQL | HPDP |
| PRDX5_MOUSE | P99029 | C96 | GVLFGVPGAFTPGC[+57]SK | HPDP |
| PRDX6_MOUSE | O08709 | C47 | DFTPVC[+57]TTELGR | HPDP |
| PRDX6_MOUSE | O08709 | C47 | DFTPVC[+329]TTELGR | IodoTMT6 |
| PRP19_MOUSE | Q99KP6 | C298 | IWSVPNTSC[+57]VQVVR | HPDP |
| PRS40_MOUSE | A6H6T1 | C60 | STLSLSEVC[+57]GK | HPDP |
| PRS6A_MOUSE | O88685 | C399 | C[+125]TDDFNGAQC[+329]K | IodoTMT6 |
| PRS7_MOUSE | P46471 | C377 | LC[+57]PNSTGAEIR | HPDP |
| PRS7_MOUSE | P46471 | C389 | SVC[+57]TEAGMFAIR | HPDP |
| PRS7_MOUSE | P46471 | C389 | SVC[+57]TEAGM[+16]FAIR | HPDP |
| PSA6_MOUSE | Q9QUM9 | C154, C161 | C[+57]DPAGYYC[+57]GFK | HPDP |
| PSA6_MOUSE | Q9QUM9 | C167 | C[+125]DPAGYYC[+57]GFK | HPDP |
| PSA6_MOUSE | Q9QUM9 | C161 | C[+125]DPAGYYC[+329]GFK | IodoTMT6 |
| PSD13_MOUSE | Q9WVJ2 | C114 | SSDEAVILC[+329]KTAIGALK | IodoTMT6 |
| PTGR1_MOUSE | Q91YR9 | C251 | TGPC[+57]PQGPAPEVVIYQQLR | HPDP |
| PTN1_MOUSE | P35821 | C32 | HEASDFPC[+57]K | HPDP |
| PUR2_MOUSE | Q64737 | C41 | QVLVAPGNAGTAC[+57]AGK | HPDP |
| PUR4_MOUSE | Q5SUR0 | C270 | FC[+57]DNSSAIQGK | HPDP |
| PUR9_MOUSE | Q9CWJ9 | C434 | YTQSNSVC[+57]YAK | HPDP |
| PURA2_MOUSE | P46664 | C58 | VVDLLAQDADIVC[+57]R | HPDP |
| PZP_MOUSE | Q61838 | C933 | EQTYNTLLC[+329]PQDTELQDNWSLELPPNVVEGSAR | IodoTMT6 |
| QCR1_MOUSE | Q9CZ13 | C268 | VYEEDAVPGLTPC[+329]R | IodoTMT6 |
| RABL6_MOUSE | Q5U3K5 | C501 | VAPQQC[+57]SEPETK | HPDP |
| RACK1_MOUSE | P68040 | C153 | YTVQDESHSEWVSC[+57]VR | HPDP |
| RAE1L_MOUSE | Q8C570 | C106 | VFTASC[+57]DK | HPDP |
| RB33B_MOUSE | O35963 | C48 | IIVIGDSNVGKTC[+329]LTYRFCAGR | IodoTMT6 |
| RB6I2_MOUSE | Q99MI1 | C258 | TGEPC[+57]VAELTEENFQR | HPDP |
| RBMS2_MOUSE | Q8VC70 | C217 | TPPGVAAPSDPLLC[+57]K | HPDP |

| | | | | |
|---|---|---|---|---|
| RBX1_MOUSE | P62878 | C94 | QVC[+57]PLDNR | HPDP |
| RENBP_MOUSE | P82343 | C250 | DGQVVLENVSEDGKELPGC[+57]LGR | HPDP |
| RFLB_MOUSE | Q5SVD0 | C88 | LC[+57]PLSFGEGVEFDPLPPK | HPDP |
| RHG01_MOUSE | Q5FWK3 | C91 | IIVFSAC[+57]R | HPDP |
| RHG10_MOUSE | Q6Y5D8 | C587 | TSPDTTFAEPTC[+57]LSASPPNAPPR | HPDP |
| RHG29_MOUSE | Q8CGF1 | C1152 | SSDSC[+57]PATAVR | HPDP |
| RHOA_MOUSE | Q9QUI0 | C164 | IGAFGYM[+16]EC[+57]SAK | HPDP |
| RL12_MOUSE | P35979 | C141 | EILGTAQSVGC[+57]NVDGR | HPDP |
| RL12_MOUSE | P35979 | C17 | C[+57]TGGEVGATSALAPK | HPDP |
| RL12_MOUSE | P35979 | C17 | C[+329]TGGEVGATSALAPK | IodoTMT6 |
| RL13A_MOUSE | P19253 | C38 | C[+57]EGINISGNFYR | HPDP |
| RL18A_MOUSE | P62717 | C64 | SSGEIVYC[+57]GQVFEK | HPDP |
| RL18A_MOUSE | P62717 | C64 | SSGEIVYC[+329]GQVFEKSPLR | IodoTMT6 |
| RL18A_MOUSE | P62717 | C64 | SSGEIVYC[+329]GQVFEK | IodoTMT6 |
| RL23_MOUSE | P62830 | C125 | EC[+57]ADLWPR | HPDP |
| RL27A_MOUSE | P14115 | C144 | GVGGAC[+57]VLVA | HPDP |
| RL28_MOUSE | P41105 | C13 | NC[+57]SSFLIK | HPDP |
| RL28_MOUSE | P41105 | C13 | NC[+57]SSFLIKR | HPDP |
| RL30_MOUSE | P62889 | C92 | VC[+57]TLAIIDPGDSDIIR | HPDP |
| RL30_MOUSE | P62889 | C52 | LVILANNC[+329]PALR | IodoTMT6 |
| RL30_MOUSE | P62889 | C92 | VC[+329]TLAIIDPGDSDIIR | IodoTMT6 |
| RL36_MOUSE | P47964 | C48 | EVC[+57]GFAPYER | HPDP |
| RL36A_MOUSE | P83882 | C72, C77 | LEC[+57]VEPNC[+57]R | HPDP |
| RL37A_MOUSE | P61514 | C48 | YTC[+125]SFC[+57]GK | HPDP |
| RL37A_MOUSE | P61514 | C42 | YTC[+125]SFC[+329]GK | IodoTMT6 |
| RL4_MOUSE | Q9D8E6 | C101 | SGQGAFGNM[+16]C[+57]R | HPDP |
| RL4_MOUSE | Q9D8E6 | C208 | GPC[+57]IIYNEDNGIIK | HPDP |
| RL7A_MOUSE | P12970 | C182 | MGVPYC[+57]IIK | HPDP |
| RL9_MOUSE | P51410 | C134 | TGVAC[+57]SVSQAQK | HPDP |
| RLA0_MOUSE | P14869 | C119 | AGAIAPC[+57]EVTVPAQNTGLGPEK | HPDP |
| RLA0_MOUSE | P14869 | C119 | AGAIAPC[+329]EVTVPAQNTGLGPEK | IodoTMT6 |
| RLA0_MOUSE | P14869 | C119 | AGAIAPC[+329]EVTVPAQNTGLGPEKTSFFQALGITTKISR | IodoTMT6 |

| | | | | |
|---|---|---|---|---|
| RLA0_MOUSE | P14869 | C119 | AGAIAPC[+329]EVTVPAQNTGLGPEKTSFFQALGITTK | IodoTMT6 |
| RN126_MOUSE | Q91YL2 | C32 | C[+57]ESGFIEELPEETR | HPDP |
| RPAP3_MOUSE | Q9D706 | C341 | DC[+57]TQAIVLDGSYSK | HPDP |
| RPB2_MOUSE | Q8CFI7 | C221 | YAYTGEC[+57]R | HPDP |
| RPB2_MOUSE | Q8CFI7 | C892 | DC[+57]STFLR | HPDP |
| RRAGC_MOUSE | Q99K70 | C376 | SC[+57]SHQTSAPSLK | HPDP |
| RRAS2_MOUSE | P62071 | C55 | QC[+57]VIDDR | HPDP |
| RRBP1_MOUSE | Q99PL5 | C1327 | EAEETQNSLQAEC[+57]DQYR | HPDP |
| RRBP1_MOUSE | Q99PL5 | C1327 | LREAEETQNSLQAEC[+57]DQYR | HPDP |
| RRBP1_MOUSE | Q99PL5 | C1198 | LKELESQVSC[+329]LEK | IodoTMT6 |
| RRBP1_MOUSE | Q99PL5 | C1327 | LREAEETQNSLQAEC[+329]DQYR | IodoTMT6 |
| RRBP1_MOUSE | Q99PL5 | C1327 | EAEETQNSLQAEC[+329]DQYR | IodoTMT6 |
| RS11_MOUSE | P62281 | C131 | DVQIGDIVTVGEC[+57]RPLSK | HPDP |
| RS11_MOUSE | P62281 | C60 | C[+329]PFTGNVSIR | IodoTMT6 |
| RS16_MOUSE | P14131 | C25 | TATAVAHC[+57]K | HPDP |
| RS17_MOUSE | P63276 | C35 | VC[+57]EEIAIIPSK | HPDP |
| RS17_MOUSE | P63276 | C35 | VC[+57]EEIAIIPSKK | HPDP |
| RS27A_MOUSE | P62983 | C144, C155 | C[+57]C[+125]LTYC[+57]FNKPEDK | HPDP |
| RS3_MOUSE | P62908 | C119 | AC[+57]YGVLR | HPDP |
| RS3_MOUSE | P62908 | C134 | GC[+57]EVVVSGK | HPDP |
| RS5_MOUSE | P97461 | C66 | AQC[+57]PIVER | HPDP |
| RS6_MOUSE | P62754 | C12 | LNISFPATGC[+57]QK | HPDP |
| RS8_MOUSE | P62242 | C100 | NC[+57]IVLIDSTPYR | HPDP |
| RSSA_MOUSE | P14206 | C163 | YVDIAIPC[+57]NNK | HPDP |
| RTCB_MOUSE | Q99LF4 | C193 | EGYAWAEDKEHC[+329]EEYGR | IodoTMT6 |
| RUS1_MOUSE | Q91W34 | C12 | APLC[+57]TEQFGSGAPR | HPDP |
| RUVB2_MOUSE | Q9WTM5 | C227 | FVQC[+57]PDGELQK | HPDP |
| RUXF_MOUSE | P62307 | C66 | C[+329]NNVLYIR | IodoTMT6 |
| S2533_MOUSE | Q3TZX3 | C30 | ATGTQQKENTLLHLFAGGC[+125]GGTVGAIFTC[+329]PLEVI | IodoTMT6 |
| S27A1_MOUSE | Q60714 | C80 | AGDTIPC[+57]IFQAVAR | HPDP |
| S7A6O_MOUSE | Q7TPE5 | C27 | NAEPAEALVLAC[+57]K | HPDP |
| SAC1_MOUSE | Q9EP69 | C445 | NAWADNANAC[+57]AK | HPDP |

| | | | | |
|---|---|---|---|---|
| SAFB1_MOUSE | Q80YR5 | C219 | ILDILGETC[+329]K | IodoTMT6 |
| SAHH_MOUSE | P50247 | C278 | EGNIFVTTTGC[+57]VDIILGR | HPDP |
| SAHH_MOUSE | P50247 | C297 | DDAIVC[+57]NIGHFDVEIDVK | HPDP |
| SAHH3_MOUSE | Q68FL4 | C189 | GSSDFC[+57]VK | HPDP |
| SAMH1_MOUSE | Q60710 | C342 | IC[+57]EVEYK | HPDP |
| SAMH1_MOUSE | Q60710 | C614 | TSSC[+57]LQEVSK | HPDP |
| SBP1_MOUSE | P17563 | C31 | C[+125]GPGYSTPLEAMKGPREEIVYLPC[+329]IYR | IodoTMT6 |
| SBP1_MOUSE | Q63836 | C31 | GPREEIVYLPC[+329]IYR | IodoTMT6 |
| SBP1_MOUSE | Q63836 | C31 | EEIVYLPC[+329]IYR | IodoTMT6 |
| SC23A_MOUSE | Q01405 | C74 | AVLNPLC[+57]QVDYR | HPDP |
| SC23B_MOUSE | Q9D662 | C425 | IAGAIGPC[+57]VSLNVK | HPDP |
| SC23B_MOUSE | Q9D662 | C434 | GPC[+57]VSENELGVGGTSQWK | HPDP |
| SC23B_MOUSE | Q9D662 | C74 | AILNPLC[+57]QVDYR | HPDP |
| SC31A_MOUSE | Q3UPL0 | C173 | TQPPEDISC[+57]IAWNR | HPDP |
| SC31A_MOUSE | Q3UPL0 | C60 | SC[+57]ATFSSSHR | HPDP |
| SEC62_MOUSE | Q8BU14 | C82 | ESVVDYC[+57]NR | HPDP |
| SF01_MOUSE | Q64213 | C279 | SITNTTVC[+57]TK | HPDP |
| SGT1_MOUSE | Q9CX34 | C79 | SLELNPNNC[+57]TALLR | HPDP |
| SH3G1_MOUSE | Q62419 | C277 | EPFELGELEQPNGGFPC[+57]APAPK | HPDP |
| SH3G1_MOUSE | Q62419 | C311 | SMPPLDQPSC[+329]K | IodoTMT6 |
| SHRM2_MOUSE | A2ALU4 | C886 | SLATSC[+57]GEILSDR | HPDP |
| SI1L1_MOUSE | Q8C0T5 | C585 | HSTARGLPLKEVLEHVIPELNVQC[+329]LR | IodoTMT6 |
| SIA7B_MOUSE | P70277 | C65 | KSRLC[+329]QHSLSLAIQK | IodoTMT6 |
| SIDT2_MOUSE | Q8CIF6 | C430 | QYLC[+57]VADLAR | HPDP |
| SLK_MOUSE | O54988 | C1136 | DLQLQC[+57]EANVR | HPDP |
| SMD2_MOUSE | P62317 | C46 | NNTQVLINC[+329]R | IodoTMT6 |
| SMD2_MOUSE | P62317 | C46 | EEEEFNTGPLSVLTQSVKNNTQVLINC[+329]R | IodoTMT6 |
| SMD2_MOUSE | P62317 | C46 | REEEEFNTGPLSVLTQSVKNNTQVLINC[+329]R | IodoTMT6 |
| SMD3_MOUSE | P62320 | C20 | VLHEAEGHIVTC[+329]ETNTGEVYR | IodoTMT6 |
| SMU1_MOUSE | Q3UKJ7 | C383 | TTEC[+57]SNTFK | HPDP |
| SND1_MOUSE | Q78PY7 | C152 | LSEC[+57]EEQAK | HPDP |
| SODC_MOUSE | P08228 | C147 | LAC[+57]GVIGIAQ | HPDP |

| | | | | |
|---|---|---|---|---|
| SODC_MOUSE | P08228 | C147 | TGNAGSRLAC[+57]GVIGIAQ | HPDP |
| SODC_MOUSE | P08228 | C7 | AVC[+329]VLKGDGPVQGTIHFEQK | IodoTMT6 |
| SPAS2_MOUSE | Q8K1N4 | C356 | FTC[+57]DVETLK | HPDP |
| SPCS2_MOUSE | Q9CYN2 | C26 | SGGGGGSSGAGGGPSC[+57]GTSSSR | HPDP |
| SPEG_MOUSE | Q62407 | C2710 | APC[+57]TYTLER | HPDP |
| SPNS1_MOUSE | Q8R0G7 | C44 | SGELEVPDC[+57]EGLQR | HPDP |
| SPSB3_MOUSE | Q571F5 | C271 | VIRSC[+329]ASSTSLQYLCCYRLR | IodoTMT6 |
| SPTN1_MOUSE | P16546 | C1930 | VNDVC[+57]TNGQDLIK | HPDP |
| SRB4D_MOUSE | A1L0T3 | C138 | QLGCGLALPVPRPLAFGQGRGPIFLDNVEC[+329]R | IodoTMT6 |
| SRBP1_MOUSE | Q9WTN3 | C738,C753 | QAC[+329]LAQSGSVPLAMQWLC[+329]HPVGHR | IodoTMT6 |
| SRC_MOUSE | P05480 | C408 | AANILVGENLVC[+329]K | IodoTMT6 |
| SRCRL_MOUSE | Q8BV57 | C6 | MRGLAC[+329]LLAM[+16]LVGIQAIER | IodoTMT6 |
| SRP09_MOUSE | P49962 | C48 | VTDDLVC[+57]LVYR | HPDP |
| SRRT_MOUSE | Q99MR6 | C489 | EC[+57]ELSPGVNR | HPDP |
| SRSF1_MOUSE | Q6PDM2 | C148 | EAGDVC[+57]YADVYR | HPDP |
| SRSF3_MOUSE | P84104 | C6, C10 | DSC[+57]PLDC[+57]K | HPDP |
| SSU72_MOUSE | Q9CY97 | C12 | VAVVC[+57]SSNQNR | HPDP |
| STA5B_MOUSE | P42232 | C688 | YYTPVPC[+57]EPATAK | HPDP |
| STIM1_MOUSE | P70302 | C49 | NTGASSGATSEESTEAEFC[+329]R | IodoTMT6 |
| STIP1_MOUSE | Q60864 | C461 | ALDLDSSC[+57]K | HPDP |
| STIP1_MOUSE | Q60864 | C461 | ALDLDSSC[+57]KEAADGYQR | HPDP |
| STK39_MOUSE | Q9Z1W9 | C461 | EGPC[+57]AVNLVLR | HPDP |
| STRN4_MOUSE | P58404 | C569 | LASC[+57]SADGTVR | HPDP |
| STX7_MOUSE | O70439 | C28 | ITQC[+57]SVEIQR | HPDP |
| STXB5_MOUSE | Q8K400 | C293 | KPEPC[+329]KPILKVELKTTR | IodoTMT6 |
| SUCB1_MOUSE | Q9Z2I9 | C164 | IC[+125]NQVLVC[+57]ER | HPDP |
| SUCB1_MOUSE | Q9Z2I9 | C430 | ILAC[+57]DDLDEAAK | HPDP |
| SYDC_MOUSE | Q922B2 | C76 | QC[+57]FLVLR | HPDP |
| SYEP_MOUSE | Q8CGC7 | C697 | EAPC[+57]ILIYIPDGHTK | HPDP |
| SYEP_MOUSE | Q8CGC7 | C910 | VAC[+57]QGEVVR | HPDP |
| SYK_MOUSE | Q99MN1 | C432 | AVEC[+57]PPPR | HPDP |
| SYNPO_MOUSE | Q8CC35 | C686 | ASPAAAEEAVPEWASC[+57]LK | HPDP |

| | | | | |
|---|---|---|---|---|
| SYSM_MOUSE | Q9JJL8 | C425 | YGEVTSASNC[+57]TDFQSR | HPDP |
| SYT2_MOUSE | P46097 | C91 | IPLPPWALIAM[+16]AVVAGLLLLTCC[+57]FCIC[+57]KKCC[+3 | IodoTMT6 |
| SYTC_MOUSE | Q9D0R2 | C266 | C[+125]GPLIDLC[+57]R | HPDP |
| TALDO_MOUSE | Q93092 | C250 | ALAGC[+57]DFLTISPK | HPDP |
| TARA_MOUSE | Q99KW3 | C1491 | SC[+57]TDVTEYAVQR | HPDP |
| TARA_MOUSE | Q99KW3 | C1921 | SFIASQGTGNSC[+57]GR | HPDP |
| TARA_MOUSE | Q99KW3 | C1930 | SSC[+57]ELEVLLR | HPDP |
| TARA_MOUSE | Q99KW3 | C1491 | SC[+329]TDVTEYAVQR | IodoTMT6 |
| TB182_MOUSE | P58871 | C259 | LAC[+57]SEAPTDVSK | HPDP |
| TBA1B_MOUSE | P68368 | C295 | AYHEQLSVAEITNAC[+329]FEPANQM[+16]VK | IodoTMT6 |
| TBA1B_MOUSE | P68368 | C295 | AYHEQLSVAEITNAC[+329]FEPANQMVK | IodoTMT6 |
| TBCD_MOUSE | Q8BYA0 | C665 | AVQSLKQIHQQLC[+329]DRHLYR | IodoTMT6 |
| TBPL1_MOUSE | P62340 | C68 | IIC[+57]TGATSEEEAK | HPDP |
| TCPA_MOUSE | P11983 | C147 | DC[+57]LINAAK | HPDP |
| TCPA_MOUSE | P11983 | C357 | IC[+57]DDELILIK | HPDP |
| TCPA_MOUSE | P11983 | C357 | IC[+57]DDELILIKNTK | HPDP |
| TCPD_MOUSE | P80315 | C295 | TGC[+57]NVLLIQK | HPDP |
| TCPG_MOUSE | P80318 | C403 | NLQDAM[+16]QVC[+57]R | HPDP |
| TCPH_MOUSE | P80313 | C370 | TC[+57]TIILR | HPDP |
| TERA_MOUSE | Q01853 | C105 | LGDVISIQPC[+57]PDVK | HPDP |
| TERA_MOUSE | Q01853 | C69, C77 | EAVC[+57]IVLSDDTC[+57]SDEK | HPDP |
| TERA_MOUSE | Q01853 | C184 | VVETDPSPYC[+125]IVAPDTVIHC[+329]EGEPIKR | IodoTMT6 |
| TERA_MOUSE | Q01853 | C69 | EAVC[+329]IVLSDDTC[+125]SDEKIR | IodoTMT6 |
| TERA_MOUSE | Q01853 | C69 | EAVC[+329]IVLSDDTC[+125]SDEK | IodoTMT6 |
| TERA_MOUSE | Q01853 | C77 | EAVC[+125]IVLSDDTC[+329]SDEK | IodoTMT6 |
| TERA_MOUSE | Q01853 | C77 | EAVC[+125]IVLSDDTC[+329]SDEKIR | IodoTMT6 |
| TGM2_MOUSE | P21981 | C27 | DHHTADLC[+57]QEK | HPDP |
| TGM2_MOUSE | P21981 | C370 | SEGTYC[+57]C[+125]GPVSVR | HPDP |
| TGM2_MOUSE | P21981 | C370, C371 | SEGTYC[+57]C[+57]GPVSVR | HPDP |
| TGM2_MOUSE | P21981 | C553 | YSGC[+57]LTESNLIK | HPDP |
| TGM2_MOUSE | P21981 | C10 | C[+329]DLEIQANGR | IodoTMT6 |
| THIM_MOUSE | Q8BWT1 | C382 | YAVGSAC[+57]IGGGQGIALIIQNTA | HPDP |

| | | | | |
|---|---|---|---|---|
| THIM_MOUSE | Q8BWT1 | C92, C103, C10 | LC[+57]GSGFQSIVSGC[+57]QEIC[+57]SK | HPDP |
| TIF1B_MOUSE | Q62318 | C628 | LASPSGSTSSGLEVVAPEVTSAPVSGPGILDDSATIC[+329]R | IodoTMT6 |
| TINAL_MOUSE | Q99JR5 | C445 | GTNEC[+57]DIETFVLGVWGR | HPDP |
| TINAL_MOUSE | Q99JR5 | C326 | C[+329]PNGQVDSNDIYQVTPAYR | IodoTMT6 |
| TIPRL_MOUSE | Q8BH58 | C87 | VAC[+57]AEEWQESR | HPDP |
| TITIN_MOUSE | A2ASS6 | C13473 | C[+57]EVSKDVPVK | HPDP |
| TITIN_MOUSE | A2ASS6 | C14323 | ILIIQNAQLEDAGSYNC[+57]R | HPDP |
| TITIN_MOUSE | A2ASS6 | C16418 | C[+57]NEHLVPVLTYTAK | HPDP |
| TITIN_MOUSE | A2ASS6 | C18063 | EC[+57]MYTIPK | HPDP |
| TITIN_MOUSE | A2ASS6 | C18869 | VPDLLEGC[+57]QYEFR | HPDP |
| TITIN_MOUSE | A2ASS6 | C20340 | C[+57]NAAAQLIR | HPDP |
| TITIN_MOUSE | A2ASS6 | C2115 | VVGKPDPEC[+57]EWYK | HPDP |
| TITIN_MOUSE | A2ASS6 | C21280 | C[+57]DPPVISNITK | HPDP |
| TITIN_MOUSE | A2ASS6 | C21561 | YILTLENSC[+57]GK | HPDP |
| TITIN_MOUSE | A2ASS6 | C21561 | YILTLENSC[+57]GKK | HPDP |
| TITIN_MOUSE | A2ASS6 | C21780 | DLPDLC[+57]YLAK | HPDP |
| TITIN_MOUSE | A2ASS6 | C21834 | VSVESTAVNTTLVVYDC[+57]QK | HPDP |
| TITIN_MOUSE | A2ASS6 | C24876 | SYAAVVTNC[+57]HK | HPDP |
| TITIN_MOUSE | A2ASS6 | C24969 | NTDKWSEC[+57]AR | HPDP |
| TITIN_MOUSE | A2ASS6 | C26841 | AAADEWTTC[+57]TPPSGLQGK | HPDP |
| TITIN_MOUSE | A2ASS6 | C28709 | ELQTNALVC[+57]VENSTDLASILIK | HPDP |
| TITIN_MOUSE | A2ASS6 | C29432 | YTVILDNAVC[+57]R | HPDP |
| TITIN_MOUSE | A2ASS6 | C33458 | EVYDYYC[+57]R | HPDP |
| TITIN_MOUSE | A2ASS6 | C34371 | FSC[+57]DTDGEPVPTVTWLR | HPDP |
| TITIN_MOUSE | A2ASS6 | C6675 | SSC[+57]TAVVDVSDR | HPDP |
| TITIN_MOUSE | A2ASS6 | C9704 | AEDQGQYTC[+57]K | HPDP |
| TJAP1_MOUSE | Q9DCD5 | C335 | NSPLPNC[+57]TYATR | HPDP |
| TKT_MOUSE | P40142 | C468 | AVELAANTKGIC[+329]FIR | IodoTMT6 |
| TLN1_MOUSE | P26039 | C1087 | C[+57]TQDLGNSTK | HPDP |
| TLN1_MOUSE | P26039 | C956 | ASAGPQPLLVQSC[+57]K | HPDP |
| TLR11_MOUSE | Q6R5P0 | C743 | TLLFSFLATNCPHGTEFWGFLTSFILLLLLIILPLISC[+329]PK | IodoTMT6 |
| TMM65_MOUSE | Q4VAE3 | C31 | SLRPGPAAAPRLPSWCC[+329]CGRGLLALGVPGGPR | IodoTMT6 |

| | | | | |
|---|---|---|---|---|
| TNNC1_MOUSE | P19123 | C35 | AAFDIFVLGAEDGC[+57]ISTK | HPDP |
| TNR19_MOUSE | Q9JLL3 | C189 | DTALAAVICSALATVLLALLILC[+329]VIYCK | IodoTMT6 |
| TNR19_MOUSE | Q9JLL3 | C25 | MALKVLPLHRTVLFAAILFLLHLAC[+329]K | IodoTMT6 |
| TNS2_MOUSE | Q8CGB6 | C548 | LLGGC[+57]GVASAGR | HPDP |
| TOM34_MOUSE | Q9CYG7 | C222 | YSESLLC[+57]SSLESATYSNR | HPDP |
| TPIS_MOUSE | P17751 | C177 | VSHALAEGLGVIAC[+57]IGEK | HPDP |
| TPIS_MOUSE | P17751 | C268 | IIYGGSVTGATC[+57]K | HPDP |
| TPIS_MOUSE | P17751 | C268 | SNVNDGVAQSTRIIYGGSVTGATC[+57]K | HPDP |
| TPIS_MOUSE | P17751 | C268 | IIYGGSVTGATC[+329]K | IodoTMT6 |
| TPM1_MOUSE | P58771 | C190 | C[+329]AELEEELK | IodoTMT6 |
| TPM1_MOUSE | P58771 | C190 | C[+329]AELEEELKTVTNNLK | IodoTMT6 |
| TPM2_MOUSE | P58774 | C190 | C[+329]GDLEEELKIVTNNLK | IodoTMT6 |
| TPM4_MOUSE | P58774 | C190 | C[+329]GDLEEELK | IodoTMT6 |
| TPM4_MOUSE | Q6IRU2 | C247 | EENVGLHQTLDQTLNELNC[+329]I | IodoTMT6 |
| TPP2_MOUSE | Q64514 | C150 | VALAEAC[+57]R | HPDP |
| TPP2_MOUSE | Q64514 | C209 | AC[+57]VDSNENGDLSK | HPDP |
| TPP2_MOUSE | Q64514 | C967 | GAGPGC[+57]YLAGSLTLSK | HPDP |
| TPSN_MOUSE | Q9R233 | C118 | SLSPEQNC[+57]PR | HPDP |
| TRFE_MOUSE | Q921I1 | C373 | TKC[+57]DEWSIISEGK | HPDP |
| TRFE_MOUSE | Q921I1 | C260 | KPVDQYEDC[+329]YLAR | IodoTMT6 |
| TRFE_MOUSE | Q921I1 | C350 | NQQEGVC[+329]PEGSIDNSPVK | IodoTMT6 |
| TRFE_MOUSE | Q921I1 | C373 | TKC[+329]DEWSIISEGK | IodoTMT6 |
| TRFE_MOUSE | Q921I1 | C373 | C[+329]DEWSIISEGK | IodoTMT6 |
| TRFE_MOUSE | Q921I1 | C386,C395 | IEC[+329]ESAETTEDC[+329]IEK | IodoTMT6 |
| TRFE_MOUSE | Q921I1 | C395 | IEC[+57]ESAETTEDC[+329]IEK | IodoTMT6 |
| TRFE_MOUSE | Q921I1 | C395 | TKC[+57]DEWSIISEGKIEC[+57]ESAETTEDC[+329]IEK | IodoTMT6 |
| TRFE_MOUSE | Q921I1 | C472 | SC[+329]HTGVDR | IodoTMT6 |
| TRFE_MOUSE | Q921I1 | C67 | KTSYPDC[+329]IK | IodoTMT6 |
| TRFE_MOUSE | Q921I1 | C67 | TSYPDC[+329]IK | IodoTMT6 |
| TRI42_MOUSE | Q9D2H5 | C18,C27 | ETAMCVC[+57]SPCC[+57]TWQRC[+125]C[+329]PRLFSCLCC[+ | IodoTMT6 |
| TXND5_MOUSE | Q91W90 | C107,C114 | VDC[+329]TADSDVC[+329]SAQGVR | IodoTMT6 |
| TXND5_MOUSE | Q91W90 | C114 | VDC[+57]TADSDVC[+329]SAQGVR | IodoTMT6 |

| | | | | |
|---|---|---|---|---|
| U3IP2_MOUSE | Q91WM3 | C460 | NSVC[+57]IIPLR | HPDP |
| UBA6_MOUSE | Q8C7R4 | C298 | TFC[+57]FEPLESQIK | HPDP |
| UBA6_MOUSE | Q8C7R4 | C347 | C[+57]QQDSDELLK | HPDP |
| UBA6_MOUSE | Q8C7R4 | C546 | VC[+57]PATESIYSDEFYTK | HPDP |
| UBAC1_MOUSE | Q8VDI7 | C134 | ATANLPAC[+57]STDR | HPDP |
| UBE2O_MOUSE | Q6ZPJ3 | C309 | SFC[+57]PGGTDSVSPPPSIITQENLGR | HPDP |
| UBE2O_MOUSE | Q6ZPJ3 | C365 | IAWEC[+57]PEK | HPDP |
| UBP15_MOUSE | Q8R5H1 | C264 | NSNYC[+57]LPSYTAYK | HPDP |
| UBP16_MOUSE | Q99LG0 | C24 | SAPDTVASESAEPVC[+57]R | HPDP |
| UBP4_MOUSE | P35123 | C758 | SLYFDEQESEAC[+329]EK | IodoTMT6 |
| UBP47_MOUSE | Q8BY87 | C856 | AGGDSGNVDDDC[+57]ER | HPDP |
| UBR1_MOUSE | O70481 | C279 | AGVYATC[+57]QEAK | HPDP |
| UBR1_MOUSE | O70481 | C996 | SC[+57]LVVATTSGLEC[+125]VK | HPDP |
| UBR2_MOUSE | Q6WKZ8 | C112 | VGEPTYSC[+57]R | HPDP |
| UPP1_MOUSE | P52624 | C132 | C[+57]SNITIIR | HPDP |
| USO1_MOUSE | Q9Z1Z0 | C802 | SQLC[+57]SQSLEITR | HPDP |
| UTP20_MOUSE | Q5XG71 | C2058 | KPAAPVPDARLPPQSC[+329]LLLPATPVRGGPK | IodoTMT6 |
| VAT1_MOUSE | Q62465 | C99 | AC[+57]GLNFADLM[+16]GR | HPDP |
| VATG1_MOUSE | Q9CR51 | C69 | EAAALGSHGSC[+57]SSEVEK | HPDP |
| VAV2_MOUSE | Q60992 | C196, C197 | SC[+57]C[+57]LLEIQETEAK | HPDP |
| VDAC1_MOUSE | Q60932 | C245 | YQVDPDAC[+329]FSAK | IodoTMT6 |
| VDAC2_MOUSE | Q60930 | C48 | SC[+329]SGVEFSTSGSSNTDTGK | IodoTMT6 |
| VDAC3_MOUSE | Q60931 | C36 | SC[+329]SGVEFSTSGHAYTDTGK | IodoTMT6 |
| VIGLN_MOUSE | Q8VDJ3 | C53 | AAC[+57]LESAQEPAGAWSNK | HPDP |
| VIME_MOUSE | P20152 | C328 | QVQSLTC[+329]EVDALK | IodoTMT6 |
| VIME_MOUSE | P20152 | C328 | QVQSLTC[+329]EVDALKGTNESLER | IodoTMT6 |
| VPP2_MOUSE | P15920 | C315 | KMKAIYHMLNMC[+329]SFDVTNK | IodoTMT6 |
| VPS8_MOUSE | Q0P5W1 | C1293 | EC[+57]TLEVEGQTR | HPDP |
| VTNC_MOUSE | P29788 | C179 | GQYC[+329]YELDETAVRPGYPK | IodoTMT6 |
| VTNC_MOUSE | P29788 | C473 | SIAQYWLGC[+329]PTSEK | IodoTMT6 |
| VWDE_MOUSE | Q6DFV8 | C217 | ISVELLGSLVFC[+125]RC[+329]TFDVSPTNTSVGFLIAWSR | IodoTMT6 |
| WASF2_MOUSE | Q8BH43 | C27 | QTLPSDTSELEC[+329]R | IodoTMT6 |

| | | | | |
|---|---|---|---|---|
| WDR1_MOUSE | O88342 | C225 | VC[+57]ALGESK | HPDP |
| WDR1_MOUSE | O88342 | C382 | M[+16]TVNESEQLVSC[+329]SMDDTVR | IodoTMT6 |
| WDR1_MOUSE | O88342 | C382 | MTVNESEQLVSC[+329]SMDDTVR | IodoTMT6 |
| WDR5_MOUSE | P61965 | C195 | DGSLIVSSSYDGLC[+57]R | HPDP |
| WDR5_MOUSE | P61965 | C195 | DGSLIVSSSYDGLC[+329]R | IodoTMT6 |
| WFD11_MOUSE | A2A5H7 | C8 | KPSWFPC[+329]LVFLC[+125]M[+16]LLLSALGGRK | IodoTMT6 |
| XDH_MOUSE | Q00519 | C970, C974 | C[+57]WDEC[+57]IASSQYQAR | HPDP |
| XIRP1_MOUSE | O70373 | C997 | ISGSTPC[+57]PPPSR | HPDP |
| XPO7_MOUSE | Q9EPK7 | C43 | ALVEFTNSPDC[+57]LSK | HPDP |
| XPP3_MOUSE | B7ZMP1 | C491 | IEDDVVVTQDSPLILSADC[+57]PK | HPDP |
| XRN2_MOUSE | Q9DBR1 | C276 | DC[+57]EGLPR | HPDP |
| YAP1_MOUSE | P46938 | C328 | C[+57]QELALR | HPDP |
| YIPF5_MOUSE | Q9EQQ2 | C42 | QYAGC[+57]DYSQQGR | HPDP |
| Z354C_MOUSE | Q571J5 | C248 | LHTGEKPYKC[+329]SECGKSFSHR | IodoTMT6 |
| ZFPL1_MOUSE | Q9DB43 | C56 | LC[+57]NTPLASR | HPDP |
| ZN106_MOUSE | O88466 | C7,C10 | KC[+329]ILC[+329]HIVYGSK | IodoTMT6 |
| ZN363_MOUSE | Q9CR50 | C243 | LC[+57]DSYNTAQAGGR | HPDP |
| ZN689_MOUSE | Q8BKK5 | C263 | TTHTGEKPHQC[+329]PSCGRRFAYPSLLAIHQR | IodoTMT6 |
| ZYX_MOUSE | Q62523 | C376 | QSVAVNESC[+57]GK | HPDP |
| ZYX_MOUSE | Q62523 | C379 | C[+57]NQPLAR | HPDP |
